# Inhalable Polymeric Nanoparticle Vaccine for Lysosome-targeting Co-delivery of Antigen and Adjuvant with Enhanced Immunoprotection

**DOI:** 10.64898/2026.01.21.700801

**Authors:** Chenxi Dai, Hanchen Zhang, Lingfei Hu, Xiaolin Song, Xi Zhang, Shengnan Fu, Zhixin Li, Haihua Xiao, Dongsheng Zhou

## Abstract

Conventional subunit vaccines suffer from premature clearance and poor synchronization of antigen and adjuvant, limiting coordinated immune activation. Here, we develop an amphipathic polymer (YAXA) with acid-labile imine bonds for acid-sensitive degradation and terminal NHS-activated esters for antigen conjugation. YAXA is co-assembled with hydrophobic TLR7 agonist 3M-052 and conjugated to protein antigen, forming nanoparticle vaccine YM3.7. Following aerosolized intratracheal inoculation into the lung, YM3.7 is efficiently internalized by antigen-presenting cell and trafficked into lysosome, where acidic conditions trigger its dissociation and co-release of antigen and adjuvant. This spatiotemporally synchronized delivery promotes robust immune activation, including cell maturation, germinal center formation, systemic and lung-resident B/T cell response, and IgG/sIgA production. In lethal pneumonia models induced by either *Pseudomonas aeruginosa* or *Staphylococcus aureus*, YM3.7 markedly improves survival over conventional antigen and adjuvant mixture. This study presents an inhalable nanoparticle platform that coordinately activates innate, humoral, mucosal, and cellular immunity for enhanced protection.

## 1. Introduction

Respiratory infections, including those caused by Gram-negative bacterium *Pseudomonas aeruginosa* (*P. aeruginosa*)^1^ and Gram-positive bacterium *Staphylococcus aureus* (*S. aureus*)^2, 3^, continue to be a major cause of global illness and death, highlighting the need for advanced vaccination strategies to target the pathogens entering the body through the respiratory tract^4^. Subunit vaccines, i.e. antigen and adjuvant mixture formulations, are generally safer than traditional methods like inactivated or live-attenuated vaccines^5^. Effective vaccination relies on an accurate and timely delivery of antigen and adjuvant to antigen-presenting cell (APC) in order to achieve a spatiotemporal coordination of antigen presentation and innate immune activation as fulfilled by APC^6, 7^. However, conventional antigen and adjuvant mixtures have the limitations such as premature dissipation of both components and inefficient APC stimulation^8^, thereby weakening the coordination of innate immunity and adaptive immunity particularly at mucosal sites like the lung^9^. Such coordination is particularly crucial for respiratory immunoprotection, involving not only peripheral humoral immunity and cell-mediated immunity but also mucosal immunity and tissue-resident memory immunity, all of which are essential for effective frontline defense^10^.

Nanotechnologies are being utilized to develop advanced vaccines, and among them, nanoparticles as a carrier for both antigen and adjuvant show great promise^11, 12, 13^. Particularly, this dual-delivery strategy of imparting both components into the lysosome can enhance antigen presentation via major histocompatibility complex class II (MHC-II)^14^, and improve action of adjuvant on its lysosomal Toll-like receptors (TLRs) such as TLR7^15^. Polymers offer a promising approach for developing nanoparticle vaccines due to their tunable physicochemical properties and functional versatility^16^. These polymeric vaccines can be inhaled and administered to the respiratory tract, which enhances mucosal immunity and tissue-resident memory immunity—often proving to be more effective than traditional injectable vaccines^17^. Additionally, several polymer materials have been designed to co-load antigens and adjuvants as tumor nanovaccines^18, 19^. Notably, pH-responsive linkages within these polymers are sensitive to acidic environment within the lysosome. This feature allows nanoparticles to dissociate, releasing encapsulated antigen and adjuvant into this organelle^20^. However, this dual-loading and co-delivery approach has rarely been applied to develop polymeric nanoparticle vaccines for infectious diseases, and the underlying cellular and molecular mechanisms for immune activation and protection remain largely unclear.

In this study, we engineer an amphiphilic polymer (YAXA) that contains acid-sensitive imine bonds within its main chain, along with NHS-activated esters at its terminal side (Fig. 1a). Imine bonds are degradable upon acidic conditions while NHS-activated esters are suitable for antigen conjugation, endowing enhanced retention during systemic circulation and facilitating rapid payload release upon lysosomal acidification. YAXA is self-assembled with a hydrophobic TLR7 agonist 3M-052^21^ to form nanoparticle YM3.5, which encapsulates 3M-052 within its hydrophobic core. Subsequent covalent conjugation of *P. aeruginosa* or *S. aureus* protein antigen onto the surface of YM3.5 via coupling reaction yields YM3.7, a nanoparticle vaccine co-loaded with protein antigen and 3M-052 adjuvant (Fig. 1b). After aerosolized intratracheal inoculation into the lung, YM3.7 is efficiently taken up by APC and trafficked into the lysosome. Under lysosomal acidic environments, acid-sensitive imine bonds break and weak-bond CO-NH coupling dissociates, leading to dissociation of nanoparticle YM3.7 along with simultaneous release of protein antigen and 3M-052 in the lysosome. This strategy of lysosome-targeting co-delivery and subsequent co-release of antigen and adjuvant (Fig. 1c) endows greatly improved efficacy to induce a spatiotemporally ordered immune response including APC maturation, germinal center (GC) formation, recirculating B, helper T (Th) and cytotoxic T (Tc) cell response, and lung-resident non-recirculating memory B/Th/Tc cell response, and immunoglobulin G (IgG) and secretory immunoglobulin A (sIgA) production. In a mouse model of lethal *P. aeruginosa* or *S. aureus* pneumonia, three-dose immunization with nanoparticle vaccine YM3.7 significantly improves survival compared to conventional antigen and adjuvant mixture (YM3.3). This work establishes a paradigm for developing next-generation inhalable nanoparticle vaccines, capable of spatiotemporally coordinating innate immunity, humoral immunity, mucosal immunity, and cell-mediated immunity to provide enhanced immunoprotection.

**Fig. 1.**
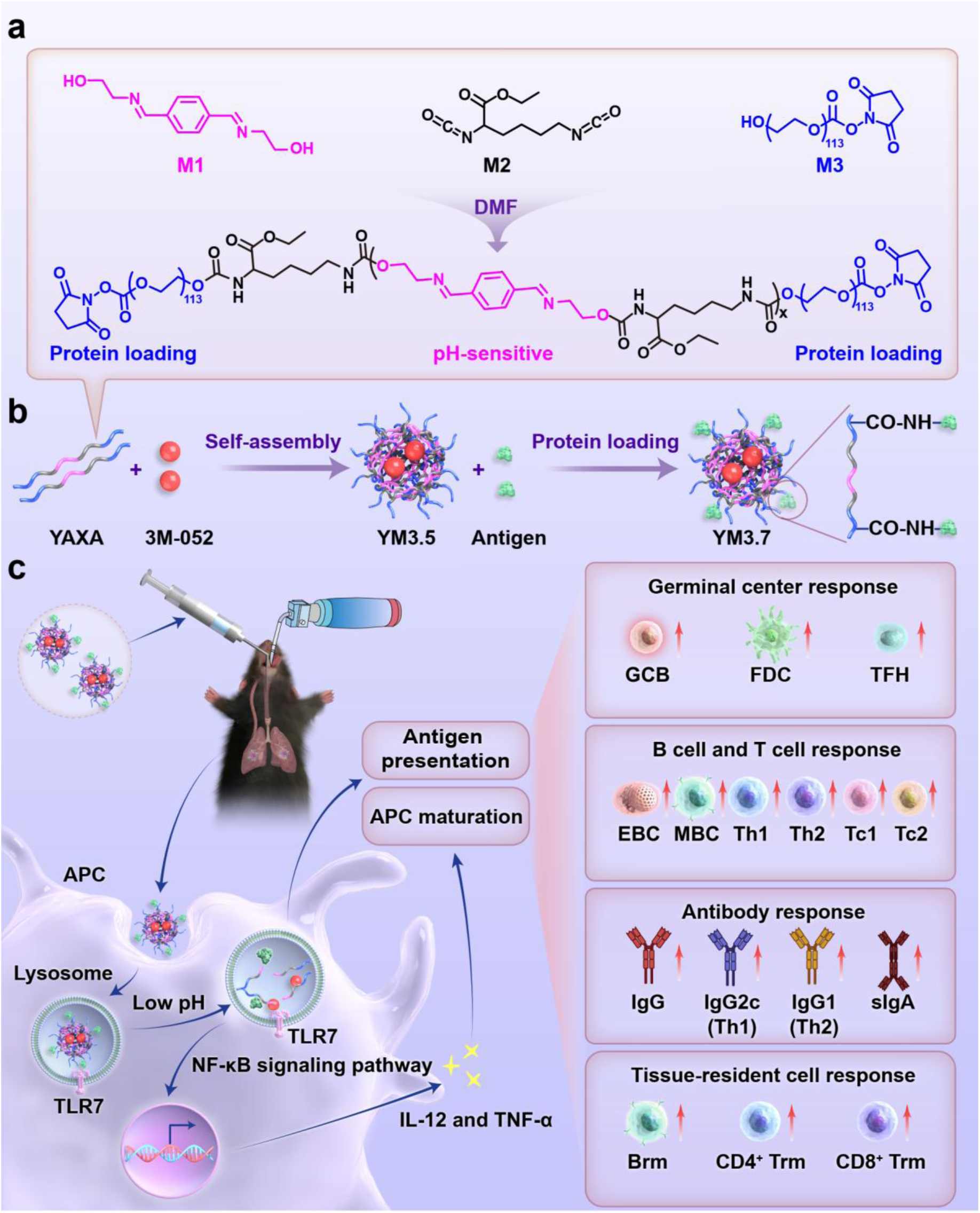
Development of nanoparticle vaccine YM3.7 for lysosome-targeting co-delivery of antigen and adjuvant. **a)** Synthesis of YAXA. The polymer YAXA is synthesized from the 3 monomers including M1, M2, and M3. Notably, M1 contains acid-sensitive imine bond. M2 provides isocyanate group for polymerization. M3 with NHS-activated ester functions as an end-capping agent in polymerization reaction. Moreover, NHS-activated esters can capture protein antigen via CO-NH coupling. **b)** Preparation of YM3.7. Self-assembly of amphiphilic YAXA with hydrophobic 3M-052 adjuvant results in nanoparticle YM3.5, and 3M-052 is encapsulated within its hydrophobic core. Further, YM3.7 is prepared by loading hydrophilic protein antigen onto YM3.5, and the amino acid residues of protein antigen are covalently conjugated to NHS-activated esters on the surface of YM3.7. **c)** Immunostimulatory and immunoprotective effects of YM3.7. YM3.7, loaded with protein antigen and 3M-052 adjuvant, is administered into the lung via aerosolized intratracheal inoculation, and then taken up by APC and trafficked into the lysosome. Acidic environments in the lysosome trigger the breakage of pH-sensitive imine bond and the cleavage of weak-bond CO-NH coupling, thereby bringing the dissociation of YM3.7 and the release of protein antigen and 3M-052. Released protein antigen is subjected to antigen presentation. Whereas, released 3M-052 binds to its receptor TLR7 located within lysosomal membrane, and then activates TLR7/NF-κB signaling pathway, thereby stimulating APC-mediated production of IL-12 and TNF-α. What’s more, secreted IL-12 and TNF-α promote APC maturation and antigen presentation. This strategy of lysosome-targeting co-delivery and subsequent co-release of antigen and adjuvant endows greatly improved efficacy to induce a spatiotemporally ordered immune response including APC maturation, GC formation, recirculating B/Th/Tc cell response, and lung-resident memory B/Th/Tc cell response, and IgG/sIgA production, thereby achieving an excellent immunoprotection against lethal infection.

## 2. Result and Discussion

### Preparation and characterizations of YM3.7

We prepared one polymer YAXA and 4 accordingly assembled nanoparticles YM3.4 to YM3.7 (Fig. 1 and Supplementary Fig. 1). YAXA was synthesized from the 3 monomers M1 to M3 via a condensation polymerization reaction (Fig. 1a). Briefly, M1 was 2,2’-(((1E,1’E)-1,4-phenylenebis(methaneylylidene))bis(azaneylylidene))bis(ethan-1-ol)) with its acid-sensitive imine bond^22^, which endowed the acidic-sensitive degradation of YAXA and thereby the controllable dissociation of nanoparticles. L-lysine diisocyanate (M2) acted as a linking monomer between M1 and M3 to form the polymer. HO-mPEG_5k_-NHS (M3) served as an end-capping agent in polymerization reaction, and its terminal NHS-activated esters mediated covalent conjugation to amino acid residues of protein antigen^23^, thereby enabling protein loading onto the nanoparticles.

Successful synthesis of M1 (Supplementary Fig. 2) and YAXA (Supplementary Fig. 3) was confirmed by proton nuclear magnetic resonance (^1^H NMR) and carbon-13 nuclear magnetic resonance (^13^C NMR) spectra. In the ^1^H NMR spectrum of YAXA, the characteristic peak at 3.5 ppm originated from methylene groups of the polyethylene glycol chain, peaks between 6.1 to 7.3 ppm were attributed to aromatic protons from M1, and the triplet peak at 1.1 ppm arose from methyl groups in M2 isocyanate residues (Supplementary Fig. 3).

YM3.4 was self-assembled from amphipathic YAXA (Supplementary Fig. S1), while YM3.5 was prepared by co-assembling YAXA with 3M-052 adjuvant (Fig. 1b). Following this, YM3.6 (Supplementary Fig. 1) and YM3.7 (Fig. 1b) were developed by covalent conjugation of protein antigen onto YM3.4 and YM3.5, respectively. It was believed that, via hydrophilic-hydrophobic interactions, hydrophobic 3M-052 was encapsulated within the hydrophobic core of YM3.5/7, while hydrophilic protein antigen was covalently conjugated to NHS-activated esters on the surface of YM3.6/7 via CO-NH coupling^23^.

We then tested 2 interdependent parameters: i) YAXA:3M-052 mass ratios determining adjuvant loading capacity; and ii) NHS-activated esters conferring protein loading capacity. First, high-performance liquid chromatography (HPLC) analysis of YM3.5 formulation revealed a dose-dependent increase in 3M-052 loading efficiency with increasing YAXA:3M-052 mass ratios, achieving almost 100% encapsulation of 3M-052 at ≥27.78:1 ratio (Supplementary Fig. 4). Second, *P. aeruginosa* protein antigen VacPAE1 was feuded from 200 to 800 µg/mL during YM3.7 formulation, and quantitative bicinchoninic acid (BCA) assay confirmed 100% antigen conjugation across all tested concentrations (Supplementary Fig. 5), indicating that NHS-activated esters were redundant for VacPAE1 loading. Accordingly, YAXA:3M-052 ratio of 27.78:1 was used for all subsequent experiments.

Next, we systematically characterized physicochemical features of the 4 nanoparticles YM3.4 to YM3.7, together with the 2 control groups YM3.2 and YM3.3 (Fig. 2a). Morphological analysis by transmission electron microscopy (TEM) and scanning electron microscopy (SEM) images revealed monodisperse spherical nanostructures for YM3.4 to YM3.6 (Supplementary Fig. 6-7) and YM3.7 (Fig. 2b-c). Dynamic light scattering (DLS) measurements indicated the hydrodynamic diameters of 117 nm for YM3.4, 135 nm for YM3.5, 175 nm for YM3.6, and 217 nm for YM3.7, each having a polydispersity index (PDI) of <0.2 (Fig. 2d, Supplementary Fig. 8). Zeta potential values were -7 mV for YM3.4, -2 mV for YM3.5, -6 mV for YM3.6, and -3 mV for YM3.7, indicating moderately negative surface charges (Fig. 2e). Subsequently, combined BCA and sodium dodecyl sulfate-polyacrylamide gel electrophoresis (SDS-PAGE) analysis confirmed 100% retention of VacPAE1 in YM3.2/3/6/7 (see Fig. 2f), with no detectable protein degradation observed (Fig. 2). Finally, DLS-based stability tracking showed that YM3.7 could maintain its overall colloidal integrity in 10% mouse serum for up to 24 h (see Fig. 2h). These results validated the successful preparation of YM3.4 to YM3.7 nanoparticles. To evaluate the generalizability of YM3.7 platform, we also prepared nanoparticles loaded with another antigen from *Staphylococcus aureus* (*S. aureus*), namely VacSAU4, and characterized their physicochemical properties (Supplementary Fig. 9a). DLS measurements confirmed the formation of monodisperse nanoparticles with hydrodynamic diameters of 141 nm for YM3.6 and 191 nm for YM3.7, and PDI<0.3 for YM3.6/7 (Supplementary Fig. 9b-c). Zeta potential values were approximately -6 mV for YM3.6 and -5 mV for YM3.7 (Supplementary Fig. 9d), consistent with moderately negative surface characteristics observed in VacPAE1-loaded nanoparticles. These findings supported the robustness of YM3.7 platform for loading different protein antigens.

**Fig. 2.**
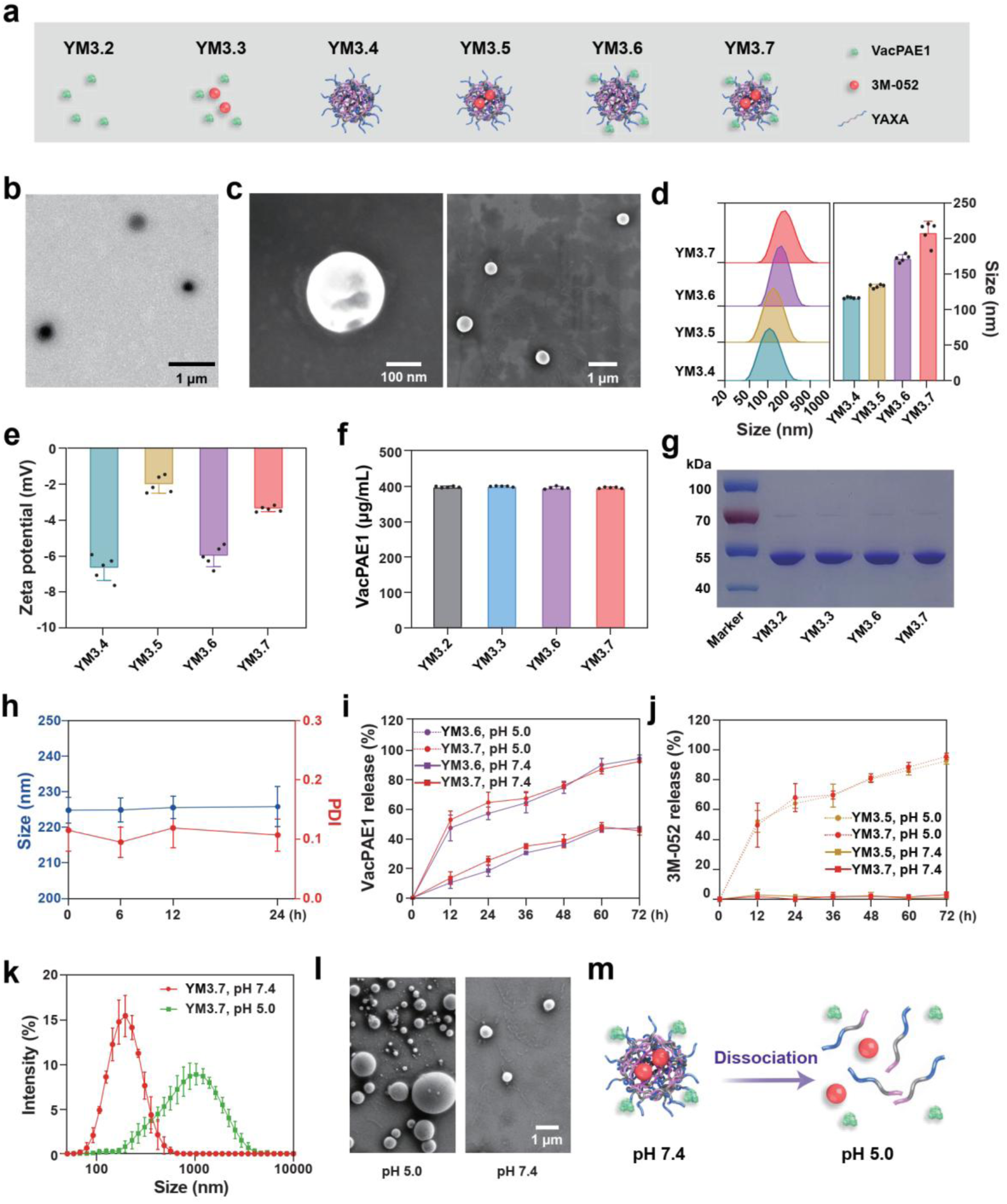
Characterization of nanoparticles YM3.4 to YM3.7. **a)** Schematic illustration of the 6 experimental groups: YM3.2 (VacPAE1), YM3.3 (mixture of VacPAE1 and 3M-052), YM3.4 (nanoparticle containing pure YAXA), YM3.5 (nanoparticle containing YAXA and 3M-052), YM3.6 (nanoparticle containing YAXA and VacPAE1), and YM3.7 (nanoparticle containing YAXA, 3M-052, and VacPAE1). **b)** TEM and **c)** SEM images of YM3.7. Experiments are repeated for 2 individual batches. DLS measurement of **d)** hydrodynamic size and **e)** zeta potential of YM3.4/5/6/7. n=5 biologically independent replicates. **f)** BCA assay of VacPAE1 content in YM3.2/3/6/7 (n=5 biologically independent replicates). **g)** SDS-PAGE analysis of VacPAE1 in YM3.2/3/6/7. Experiments are repeated for 2 individual batches. **h)** DLS measurement of hydrodynamic diameter and PDI value of YM3.7 in 10% mouse serum over 24 h (n=5 biologically independent replicates). **i)** BCA assay of release kinetics of VacPAE1 from YM3.6/7 at pH 5.0/7.4 (n=5 biologically independent replicates). **j)** HPLC determination of release kinetics of 3M-052 from YM3.5/7 at pH 5.0/7.4 (n=5 biologically independent replicates). **k)** DLS measurement of hydrodynamic diameter of YM3.7 at pH 5.0/7.4 over 72 h (n=5 biologically independent replicates). **l)** SEM image of YM3.7 at pH 5.0/7.4 over 72 h. Experiments are repeated for 2 individual batches. **m)** Schematic illustration of YM3.7 dissociation and 3M-052/VacPAE1 release under acidic conditions (pH 5.0). Data are presented as mean ± SD.

To evaluate in vitro pH-responsive characteristics of YM3.7, we quantitatively examined the release of VacPAE1 and 3M-052, as well as the dissociation of YM3.7. First, under neutral conditions (pH 7.4), YM3.6/7 exhibited VacPAE1 release rates of approximately 40% after 72 h; in contrast, under lysosome-mimicking acidic conditions (pH 5.0), the release of VacPAE1 was significantly accelerated, achieving over 90% after the same duration (Fig. 2i). For YM3.5/7, the release of 3M-052 was minimal at pH 7.4 (<1%) but nearly complete (>90%) at pH 5.0 after 72 h (Fig. 2j). Second, DLS analysis indicated that YM3.7 underwent acid-triggered structural disassembly and formed polydisperse aggregates at pH 5.0 over 72 h: a dramatic increase in hydrodynamic diameter from 200 nm to >1000 nm (Fig. 2k) plus an upshift in PDI from 0.2 to 0.9 (Supplementary Fig. 10). SEM imaging visually confirmed the dissociation of YM3.7 under acidic conditions, consistent with DLS results (Fig. 2l). In summary, YM3.7 loaded with VacPAE1 and 3M-052 could maintain its structural integrity under physiological conditions, while acidic environments would rapidly trigger the breakage of pH-sensitive imine bond and the cleavage of weak-bond CO-NH coupling, thereby leading to the dissociation of YM3.7 and the release of VacPAE1 and 3M-052 (Fig. 2m).

### Favored biosafety in vitro and in vivo

Prior to conducting in vivo immunoprotection studies, we performed biosafety evaluation experiments both in vitro and in vivo. First, Cell Counting Kit-8 (CCK-8) analysis and hemolysis assay were conducted to assess cytotoxicity of YM3.4 in vitro. For CCK-8 analysis, mouse embryonic fibroblast NIH-3T3, human ovarian epithelial cell IOSE-80, and mouse bone marrow-derived dendritic cell (BMDC) were exposed to YM3.4 ranging from 50 to 500 µg/mL for 24 h. All cell types exhibited >80% viability^24^ for all tested YM3.4 concentrations (Supplementary Fig. 11). Hemolysis assay revealed that even at the maximum concentration of 500 µg/mL, YM3.4 induced <5% hemolysis rate^24^ in mouse red blood cell after 4-h incubation (Supplementary Fig. 12). Second, biochemical blood test and histopathologic examination were conducted in mice to evaluate the biosafety of YM3.7 in vivo. There were no abnormal changes in the tested markers of hepatic function (alanine aminotransferase and aspartate aminotransferase) or renal function (blood urea nitrogen and creatinine) on 7 d post single-dose immunization (Supplementary Fig. 13). Additionally, Hematoxylin and Eosin (H&E) staining of the lung, spleen, liver, kidney, and heart tissues collected on 7 d and 28 post single-dose immunization showed no signs of inflammatory infiltration, necrosis, or structural distortion (Supplementary Fig. 14, Supplementary Fig. 15), indicating both short-term and long-term biosafety in vivo. Overall, this multi-dimensional biosafety evaluation paradigm demonstrated the highly favored biosafety of YM3.4 in vitro and YM3.7 in vivo.

### Enhanced antibody production and immunoprotection for YM3.7

This study aimed to develop YM3.7 as an optimized nanoparticle vaccine. To comprehensively evaluate its immunogenicity and protective efficacy, we performed a multi-tiered assessment in mice using *P. aeruginosa* antigen VacPAE1 and *S. aureus* antigen VacSAU4. The experimental design comprised 3 main aspects: i) comparison of YM3.7 (nanoparticle co-loaded with antigen and adjuvant) with YM3.3 (conventional antigen and adjuvant mixture) using VacPAE1; ii) comparison of YM3.7 with YM3.5+YM3.6 (physical mixture of nanoparticles YM3.5 and YM3.6) using VacPAE1; and iii) evaluation of YM3.7 with an alternative antigen VacSAU4.

First, to compare YM3.7 with YM3.3 using VacPAE1 (Fig. 3a), serum samples were analyzed by enzyme-linked immunosorbent assay (ELISA) to measure VacPAE1-specific IgG subclasses, including total IgG, Th1-associated IgG2c, and Th2-assciated IgG1^25^. Additionally, VacPAE1-specific IgG and sIgA in bronchoalveolar lavage fluid (BALF) specimens^10^ were also measured by ELISA. YM3.7 induced a balanced Th1 and Th2 IgG response as well as a robust sIgA response, and the titers of each antibody subset tested in the YM3.7 group were significantly higher than those in all other groups following three-dose immunizations (Fig. 3b-d, Supplementary Fig. 16). In addition, functional capacity of antibody elicited by YM3.7 was assessed in RAW264.7 cells infected with *P. aeruginosa* strains F291007 and PAO1. Both serum and BALF samples collected on 42 d post the first immunization demonstrated significantly higher neutralization activities (Fig. 3e, Supplementary Fig. 17) and enhanced opsonic phagocytosis (Fig. 3f, Supplementary Fig. 18) compared to all other groups. To evaluate the protection efficacy against lung infections, mice that received three-dose immunizations were challenged with a lethal dose of *P. aeruginosa* aerosols. The YM3.7 group exhibited a survival rate of 93.3%, significantly higher than that of any other group (Fig. 3g). No significant differences in body weight changes in survived mice were observed among the groups tested (Fig. S19). Furthermore, YM3.7-immunized mice demonstrated significantly lower levels of bacterial burden (Fig. 3h) and less pathological damage (Fig. 3i, Supplementary Fig. 20) in multiple organs.

**Fig. 3.**
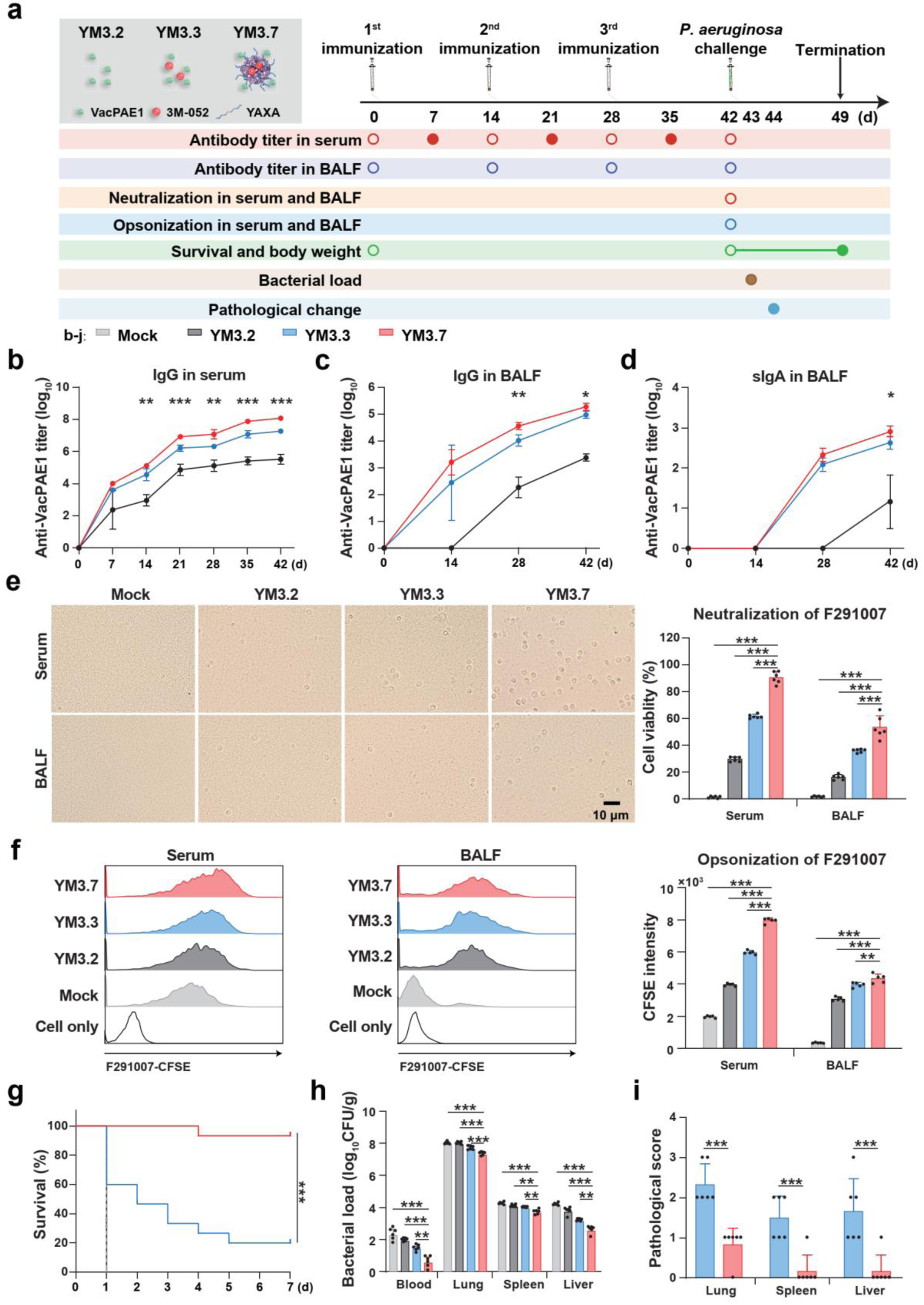
Antibody titers and immunoprotection induced by YM3.7 against *P. aeruginosa*. **a)** Schematic diagram of experimental design. C57BL/6J mice are immunized 3 times at 14-d intervals via aerosolized intratracheal inoculation with the following 4 experimental groups: Mock (PBS), YM3.2 (VacPAE1), YM3.3 (mixture of VacPAE1 and 3M-052), and YM3.7 (nanoparticle containing VacPAE1, 3M-052, and YAXA). 20 μg of VacPAE1, 18 μg of 3M-052, and 500 μg of YAXA are used per inoculation. The absolute lethal dose (LD_100_) pre-determined at 1.4×10^6^ colony-forming units (CFUs) of *P. aeruginosa* F291007 is used for aerosolized intratracheal challenge following three-dose immunizations. Hollow circles denote timepoints pre-immunization or pre-challenge. **b)** ELISA measurement of VacPAE1-specific IgG in sera (n=6 biologically independent replicates). ELISA measurement of VacPAE1-specific **c)** IgG and **d)** sIgA in BALF (n=5 biologically independent replicates). **e)** Neutralizing activity of sera and BALF against *P. aeruginosa* F291007 was assessed by CCK-8 assay (n=6 biologically independent replicates)^26^. Shown are representative microscopic images and quantification. **f)** Opsonophagocytic activity of sera and BALF was measured using Carboxyfluorescein diacetate succinimidyl ester (CFSE)-labeled *P. aeruginosa* F291007 (n=5 biologically independent replicates)^27, 28^. **g)** Survival curves of mice post *P. aeruginosa* challenge (n=15 biologically independent replicates). **i)** Plate counting of bacterial loads in the lung, spleen, liver, and blood at 6 h post *P. aeruginosa* challenge (n=6 biologically independent replicates). **j)** Pathological scores for the lung, spleen, and liver on 2 d post *P. aeruginosa* challenge (n=6 biologically independent replicates). Data are presented as mean ± SD. **Statistical significance test: b-d)** YM3.7 is compared with each of the other groups using two-way ANOVA with Dunn’s multiple comparison test; **e-f, h)** YM3.7 is compared with each of the other groups using one-way ANOVA with Dunn’s multiple comparison test; **g)** Survival analysis is conducted using Log-rank (Mantel-Cox) test; **i)** Unpaired two-tailed Student’s *t*-test is used to compare YM3.7 and YM3.3. *: *P*<0.05, **: *P*<0.01, ***: *P*<0.001.

Second, we compared YM3.7 with YM3.5+YM3.6 using VacPAE1 (Supplementary Fig. 21a). No significant differences were observed between these 2 groups with respect to VacPAE1-specific IgG titers in serum, or sIgA and IgG titers in BALF (Supplementary Fig. 21b-d). Following challenge with a lethal dose of *P. aeruginosa* aerosols, survival rates were 100% and 90% for the YM3.7 and YM3.5+YM3.6 groups, respectively, with no statistically significant difference between them (Supplementary Fig. 21e). In addition, body weight changes among survivors showed no significant intergroup differences (Supplementary Fig. 21f). Notably, although the protective efficacy was comparable, YM3.7 formulation offered distinct practical advantages for clinical translation, including streamlined manufacturing process and reduced complexity.

Third, to assess the generalizability of YM3.7, we evaluated YM3.7 loaded with an alternative antigen VacSAU4 (Fig. 4a). Consistently, YM3.7 induced a balanced Th1/Th2 IgG profile and a strong sIgA response, with all antibody subclass titers being significantly higher in the YM3.7 group than in other groups (Fig. 4b-f). Following challenge with a lethal dose of *S. aureus* aerosols, YM3.7 conferred 100% survival, significantly outperforming all other groups (Fig. 4g). Notably, starting from 2 d post *S. aureus* challenge, surviving mice in the YM3.7 group maintained significantly higher body weights compared to other groups (Fig. 4h). Furthermore, YM3.7 immunization also significantly reduced bacterial burdens in multiple organs (Fig. 4i).

**Fig. 4.**
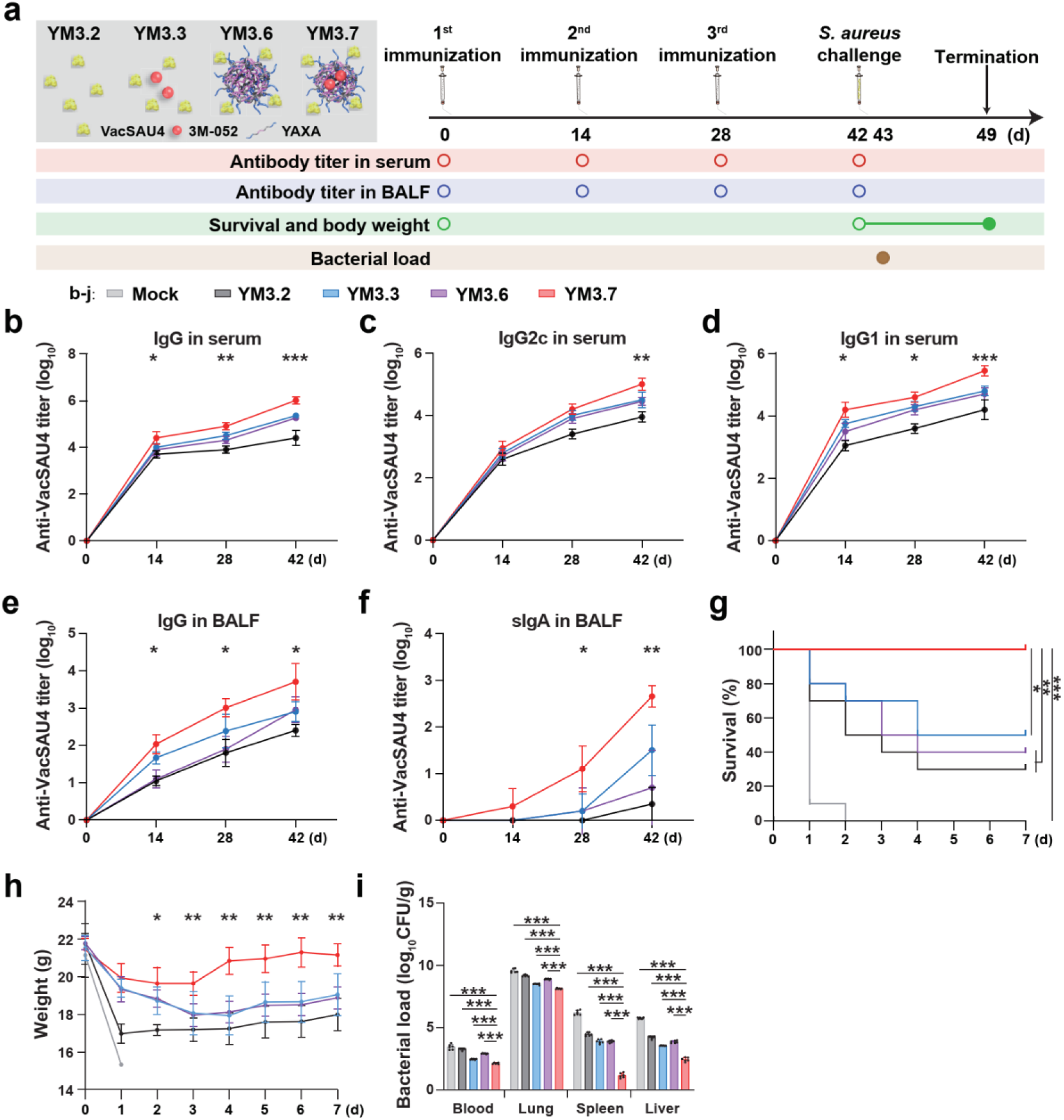
Antibody titers and immunoprotection induced by YM3.7 against *S. aureus*. **a)** Schematic diagram of experimental design. C57BL/6J mice are immunized 3 times at 14-d intervals via aerosolized intratracheal inoculation with the following 5 experimental groups: Mock (PBS), YM3.2 (VacSAU4), YM3.3 (mixture of VacSAU4 and 3M-052), YM3.6 (nanoparticle containing VacSAU4 and YAXA), and YM3.7 (nanoparticle containing VacSAU4, 3M-052, and YAXA). 20 μg of VacSAU4, 18 μg of 3M-052, or 500 μg of YAXA are used per inoculation. The LD_100_ value pre-determined at 4×10^8^ CFUs of *S. aureus* USA300-FPR3757 is used for aerosolized intratracheal challenge following three-dose immunizations. Hollow circles denote timepoints pre-immunization or pre-challenge. ELISA measurement of VacSAU4-specific **b)** IgG, **c)** IgG2c, and **d)** IgG1 in sera (n=6 biologically independent replicates). ELISA measurement of VacSAU4-specific **e)** IgG and **f)** sIgA in BALF (n=6 biologically independent replicates). **g)** Survival curves and **h)** body weight changes of mice post *S. aureus* challenge (n=10 biologically independent replicates). **i)** Plate counting of bacterial loads in the lung, spleen, liver, and blood at 6 h post *S. aureus* challenge (n=6 biologically independent replicates). Data are presented as mean ± SD. **Statistical significance test: b-f)** YM3.7 is compared with each of the other groups using two-way ANOVA with Dunn’s multiple comparison test; **g)** Survival analysis is conducted using Log-rank (Mantel-Cox) test; **i)** YM3.7 is compared with each of the other groups using one-way ANOVA with Dunn’s multiple comparison test. *: *P*<0.05, **: *P*<0.01, ***: *P*<0.001.

Collectively, these findings demonstrated that YM3.7 consistently elicited potent humoral and mucosal immunity across different protein antigens and provided superior protection against lethal bacterial pneumonia, underscoring its utility as a robust and versatile mucosal nanoparticle vaccine platform.

### Improved in vivo metabolism and intracellular localization for YM3.7

To dissect their features of in vivo metabolism and intracellular localization (Fig. 5a), we focused on comparing the nanoparticle vaccine (YM3.7) with the conventional antigen and adjuvant mixture (YM3.3), following a previously established framework^19^. First, real-time in vivo fluorescence imaging (IVFI) of the whole body revealed that YM3.7-Cy5.5 exhibited favorable pharmacokinetics in mice. The almost complete systemic clearance of YM3.7-Cy5.5 was achieved on 8 d post single-dose immunization, which was comparable to that of YM3.3-Cy5.5 (Supplementary Fig. 22a). Critically, no fluorescent signal was detected in the brain, indicating absence of cerebral accumulation and suggesting a low risk of neurotoxicity. Additionally, YM3.7-Cy5.5 displayed higher Cy5.5 fluorescence intensity during 1-3 d post single-dose immunization (Supplementary Fig. 22b), indicating a better ability for tissue residence. Second, quantitative IVFI analysis was conducted to evaluate the distribution of YM3.7-Cy5.5 and YM3.3-Cy5.5 in the pulmonary mediastinal lymph node (PMLN) and spleen. PMLN was the secondary lymphoid organ nearest to the lung^29^ while the spleen was the largest and also the primordial one in the body^30^, both of which were associated with the initiation of adaptive immune response^31^. The results revealed significantly greater accumulation of YM3.7-Cy5.5 within the PMLN (Fig. 5b) and spleen (Supplementary Fig. 23) on 1 d post single-dose immunization. Third, flow cytometry analysis of APC uptake in lung (Fig. 5c), PMLN (Fig. 5d), and spleen (Supplementary Fig. 24) on 1 d post single-dose immunization demonstrated a greater uptake of YM3.7-Cy5.5 by the following 4 major APC types (see Supplementary Fig. 25 for gating strategy): type 1 conventional dendritic cell (cDC1, CD11c^+^ MHC-II^+^ CD103^+^ CD11b^-^), type 2 cDC (cDC2, CD11c^+^ MHC-II^+^ CD103^-^CD11b^+^), plasmacytoid dendritic cell (pDC, pDCA1^+^), and macrophage (Mø, CD11b^+^ F4/80^+^)^32^. Fourth, immunofluorescence staining analysis was conducted to assess GC formation in the PMLN post three-dose immunizations. GC, situated at the core of secondary lymphoid organ, was the site of activation, proliferation, differentiation, and apoptosis of B cell, thereby serving as the foundation of humoral immunity^33^. The results showed that YM3.7 induced a greater GC formation in PMLN (Fig. 5e), which was characterized by both a larger number of B cell follicle and also a higher proportion of GC B cell (GCB, B220^+^ Ki67^+^) in total B cell (B220^+^)^34^. Fifth, confocal microscopy demonstrated that YM3.7-Cy5.5 exhibited greater uptake by BMDC (Supplementary Fig. 26), along with a higher co-localization with lysosome within BMDC (Fig. 5f), highlighting an improved capability of lysosome-targeting delivery. In summary, compared to YM3.3, YM3.7 displayed at least 4 key advantages: improved accumulation within secondary lymphoid organ, better APC uptake, enhanced GC formation, and elevated intracellular trafficking to lysosome.

**Fig. 5.**
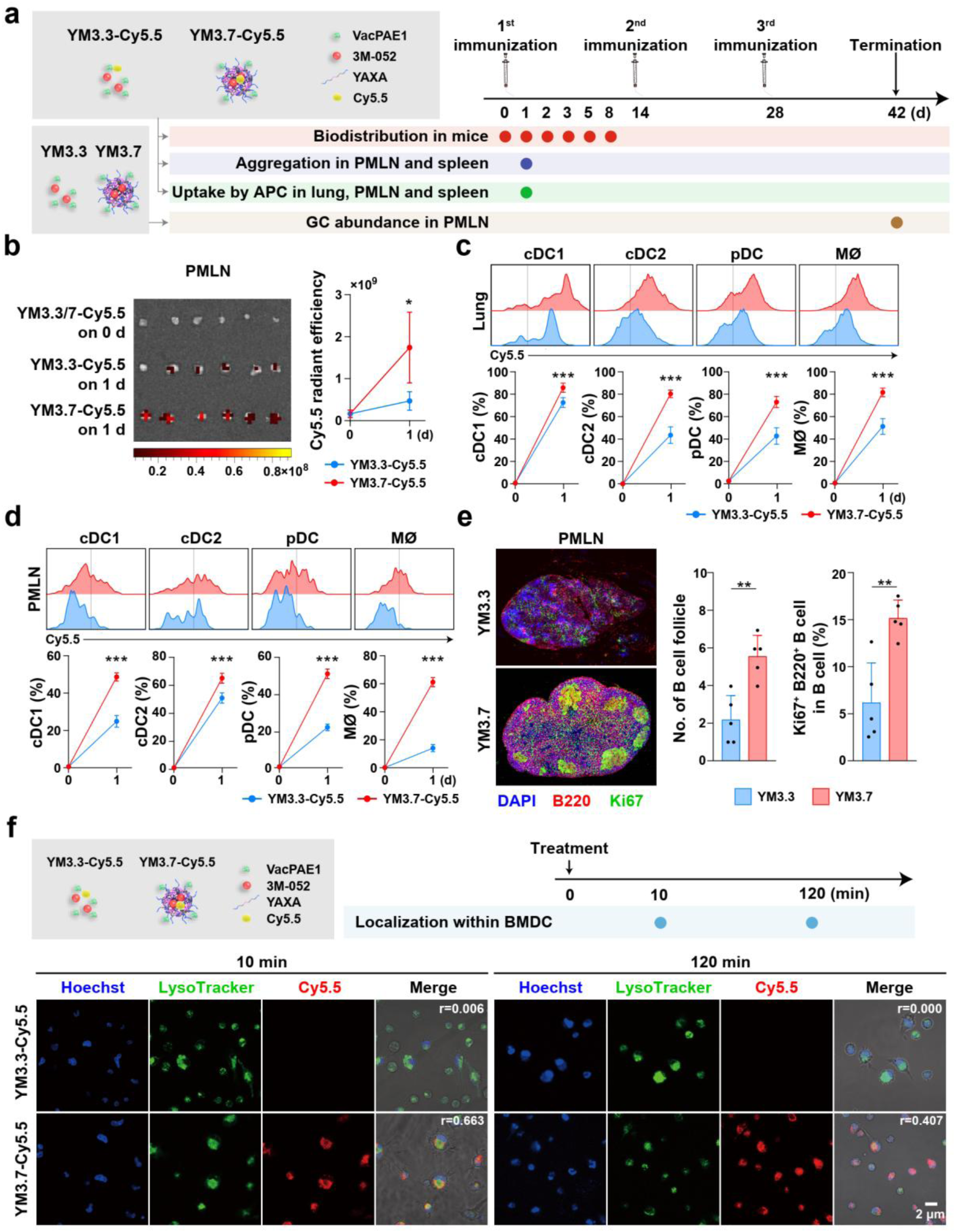
In vivo metabolism and intracellular localization for YM3.7. **a)** Schematic diagram of experimental design. YM3.7-Cy5.5 represents the nanoparticle containing VacPAE1, 3M-052, YAXA, and Cy5.5, while YM3.3-Cy5.5 stands for the mixture of VacPAE1, 3M-052, and Cy5.5. Mice are immunized with single-dose YM3.7-Cy5.5 or YM3.3-Cy5.5 unless otherwise stated. 20 μg of VacPAE1, 18 μg of 3M-052, 500 μg of YAXA, or 1 μg of Cy5.5 are used per inoculation. **b)** IVFI assessment of YM3.7-Cy5.5 or YM3.3-Cy5.5 accumulation in PMLN (n=6 biologically independent replicates). Flow cytometry analysis of uptake of YM3.7-Cy5.5 or YM3.3-Cy5.5 by APC in the **c)** lung and **d)** PMLN (n=5 biologically independent replicates). **e)** Immunofluorescence staining of GC abundance, and corresponding quantification of B cell follicles and Ki67^+^B220^+^ B cells in PMLN (n=5 biologically independent replicates). Mice are immunized 3 times at 14-d intervals using YM3.3/7 without Cy5.5 labeling. Total B cell is stained with B220 (red), and GCB is stained with B220 (red) and Ki67 (green)^34^. **f)** Confocal laser fluorescence imaging of intracellular localization of YM3.7-Cy5.5 or YM3.3-Cy5.5 within BMDC. BMDC is incubated with YM3.7-Cy5.5 or YM3.3-Cy5.5 for 10 min. 5.56 μg/mL VacPAE1, 138.89 μg/mL YAXA, 5 μg/mL 3M-052, and Cy5.5 0.278 μg/mL are used. Cell nucleus and lysosome are labeled with Hoechst (blue) and LysoTracker (green), respectively^35^. Co-localization coefficients: YM3.7 v.s. lysosome is 0.663 at 10 min and 0.407 at 120 min; YM3.3 v.s. lysosome is 0.006 at 10 min and 0.000 at 120 min (n=5 biologically independent replicates). Data are presented as mean ± SD. **Statistical significance test: b-d)** YM3.7 is compared with YM3.3 using two-way ANOVA with Dunn’s post hoc multiple comparison test; **e-f)** Unpaired two-tailed Student’s *t*-test is used to compare YM3.7 and YM3.3. **f)** Co-localization coefficient of Cy5.5 and LysoTracker is determined by Pearson correlation analysis. *: *P*<0.05, **: *P*<0.01, ***: *P*<0.001.

### Elevated cell-mediated immune response for YM3.7

To investigate cell-mediated immune response in the lung, PMLN, and spleen, we conducted a series of quantitative flow cytometry analyses to measure APC maturation, GC cell response, recirculating B/Th/Tc cell response, and lung-resident non-recirculating memory B/Th/Tc cell response induced by YM3.7 relative to YM3.3 (Fig. 6a). First, cell maturation of the 4 APC subsets cDC1, cDC2, pDC, and Mø (see Supplementary Fig. 27 for gating strategy) were evaluated on 1 d post single-dose immunization, and the results showed that YM3.7 led to a higher APC maturation in the lung (Fig. 6b), PMLN (Fig. 6c), and spleen (Supplementary Fig. 28). Second, GC cell response was assessed post three-dose immunizations by detecting the 3 key GC cell populations: GCB (CD19^+^ CD38^-^ CD95^+^)^32, 36^, follicular dendritic cell (FDC, CD45^-^ CD31^-^ CD21/35^+^ PDPN^+^)^37, 38^, and T follicular helper cell (TFH, CD4^+^ CXCR5^+^ PD1^+^)^32, 36^ (see Supplementary Fig. 29 and Supplementary Fig. 30 for gating strategy). Upon immunization, GCB differentiated into antibody-secreting B cell, FDC retained antigen for sustained B cell activation, and TFH provided co-stimulatory signals (such as CD40L and IL-21) to facilitate antibody affinity maturation of GCB, thereby synergistically orchestrating humoral immunity^39^. The results showed that YM3.7 induced higher frequencies of GCB, FDC, and TFH in the PMLN (Fig. 6d-f) and spleen (Supplementary Fig. 31), indicating an enhanced GC cell response for YM3.7. Third, B cell response (see Supplementary Fig. 32 for gating strategy) was evaluated post three-dose immunizations by analyzing effector B cell (EBC, CD44^+^ CD138^+^)^32, 36^ and memory B cell (MBC, CD19^+^ PD-L2^+^ CD80^+^)^40^ in the lung, PMLN, and spleen, as well as tissue-resident memory B cell (Brm, CD19^+^ PD-L2^+^ CD80^+^ CD69^+^)^41, 42^ in the lung. When exposure to antigen, MBC^43^ proliferated and differentiated into EBC^44^ that was responsible for antibody production, while Brm, as a tissue-localized immune surveillant, would respond to antigen faster and more robustly in barrier tissues where antigen was previously encountered^45^. The results showed that YM3.7 induced greater frequencies of EBC and MBC in all examined organs (Fig. 6g-h,j-k, Supplementary Fig. 33), and also a higher frequency of Brm in the lung (Fig. 6i), indicating an enhanced B cell response for YM3.7. Fourth, recirculating Th1/2 and Tc1/2 cell response (see Supplementary Fig. 34 for gating strategy) post three-dose immunizations was measured. Herein, Th1 or Th2 cell was defined as CD4^+^ helper T cell secreting IFN-γ or IL-13, respectively, while Tc1 or Tc2 cell as CD8^+^ Tc cell secreting IFN-γ or IL-13, respectively^36^. Following ex vivo stimulation with a VacPAE1 peptide pool, YM3.7 induced higher levels of all these T cell populations in the lung (Fig. 7a-b, d-e), PMLN (Fig. 7g-j), and spleen (Supplementary Fig. 35). Fifth, CD4^+^ tissue-resident memory T cell (Trm, CD3^+^ CD4^+^ CD44^+^ CD69^+^ CD103^+^) and CD8^+^ Trm (CD3^+^ CD8^+^ CD44^+^ CD69^+^ CD103^+^)^46, 47^ were quantified in the lung post three-dose immunizations. Trm, strategically positioned in barrier tissues without recirculating, would provide a first response against infections reencountered at barrier tissues^48^. The results showed that YM3.7 induced a greater expansion of CD4^+^/8^+^ Trm (Fig. 7c, f). Taken together, compared to YM3.3, YM3.7 exhibited a better efficacy in orchestrating a coordinated antigen-specific immune response, including GC cell response, recirculating B/Th/Tc cell response, and lung-resident memory B/Th/Tc cell response. The above constructed a multi-layered immune network in both lymphatic system and mucosal lung tissue.

**Fig. 6.**
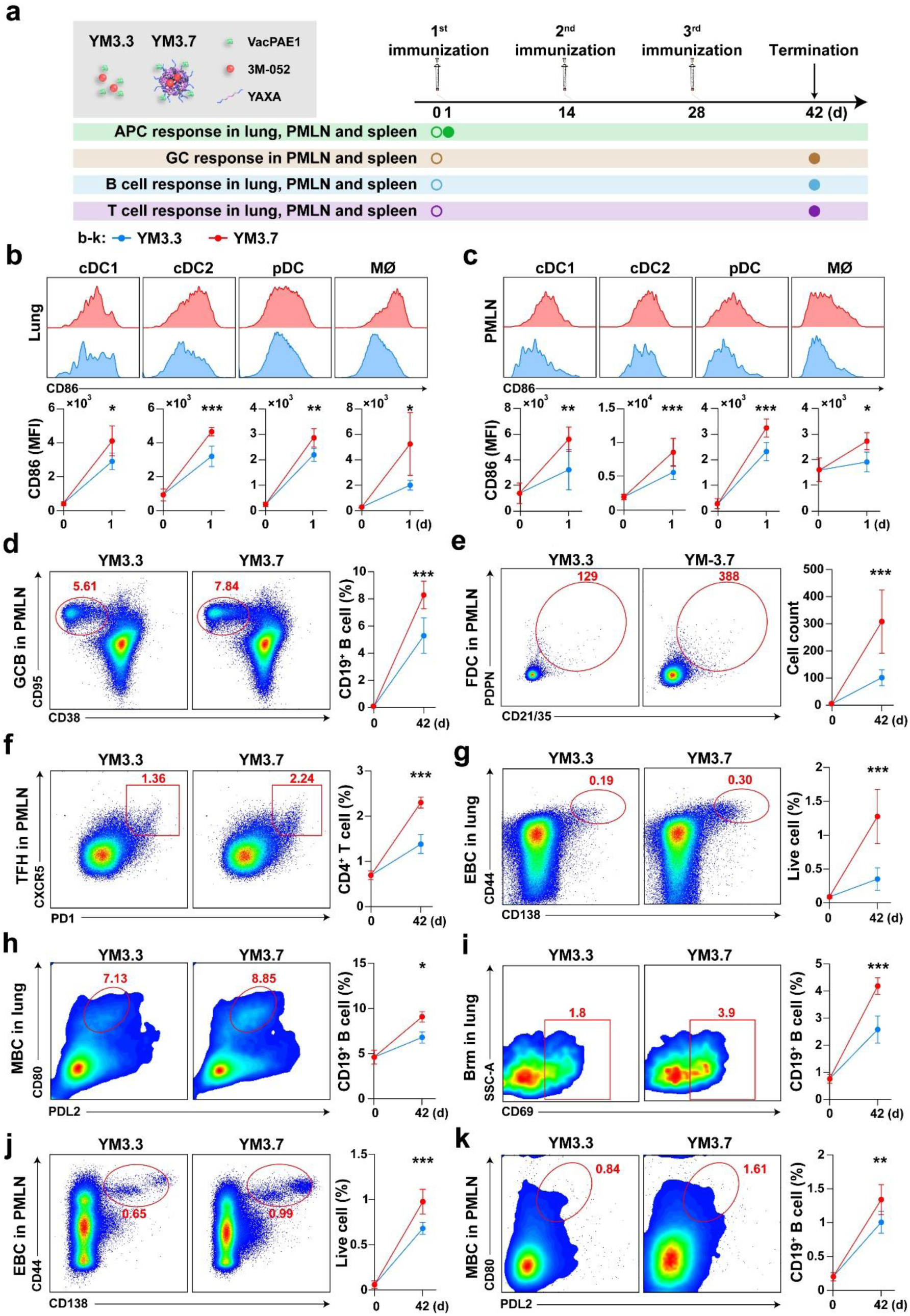
Flow cytometry analysis of APC, GC, and B cell response induced by YM3.7. **a)** Schematic diagram of experimental design. Mice are immunized with YM3.7 (nanoparticle containing VacPAE1, 3M-052, and YAXA) or YM3.3 (mixture of VacPAE1 and 3M-052). Single-dose immunization is conducted for APC response, while three-dose immunizations for GC and B cell response. 20 μg of VacPAE1, 18 μg of 3M-052, or 500 μg of YAXA are used per inoculation. Hollow circles indicate timepoints pre-immunization. n=5 biologically independent replicates. APC in the **b)** lung and **c)** PMLN. **d)** GCB, **e)** FDC, and **f)** TFH in the PMLN. **g)** EBC, **h)** MBC, and **i)** Brm in the lung. **j)** EBC and **k)** MBC in the PMLN. Data are presented as mean ± SD. **Statistical significance test:** Two-way ANOVA with Dunn’s post hoc test (YM3.7 v.s. YM3.3). *: *P*<0.05, **: *P*<0.01, ***: *P*<0.001.

**Fig. 7.**
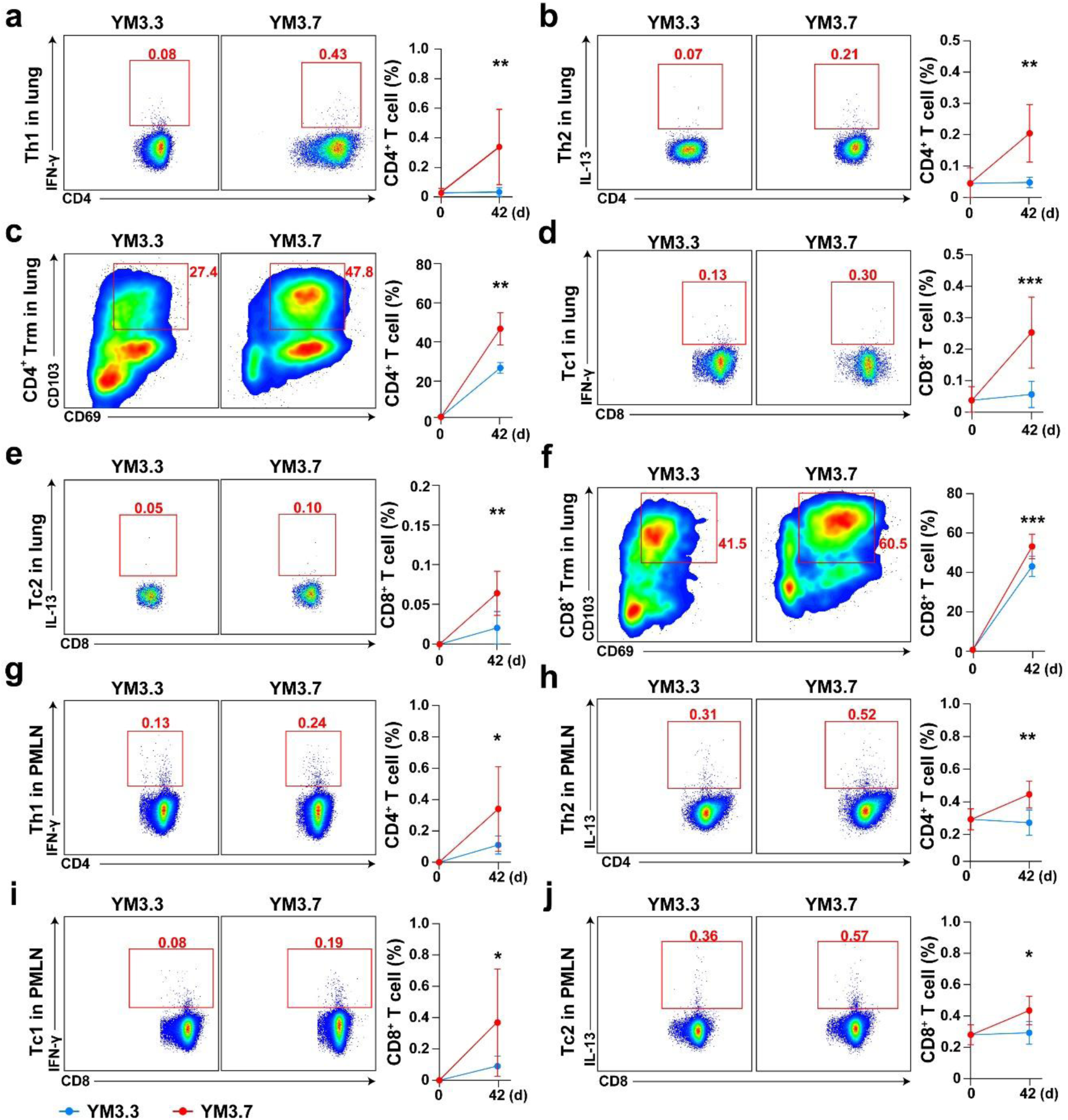
Flow cytometry analysis of VacPAE1-specific T cell response induced by YM3.7. Mice are immunized 3 times at 14-d intervals with YM3.7 (nanoparticle containing VacPAE1, 3M-052, and YAXA) or YM3.3 (mixture of VacPAE1 and 3M-052). 20 μg of VacPAE1, 18 μg of 3M-052, or 500 μg of YAXA are used per inoculation. Hollow circles indicate timepoints pre-immunization. n=5 biologically independent replicates. **a)** Th1 cell, **b)** Th2 cell, and **c)** CD4^+^ Trm in the lung. **d)** Tc1 cell, **e)** Tc2 cell, and **f)** CD8^+^ Trm in the lung. **g)** Th1 cell and **h)** Th2 cell in the PMLN. **i)** Tc1 cell and **j)** Tc2 cell in the PMLN. Data are presented as mean ± SD. **Statistical significance test:** Two-way ANOVA with Dunn’s post hoc test (YM3.7 v.s. YM3.3). *: *P*<0.05, **: *P*<0.01, ***: *P*<0.001.

### Spatiotemporally ordered immune response for YM3.7

In this study, the lung was the primary organ targeted for immunization and infection. Additionally, the lung was rich in lymphatic vessels that connected to the lymphatic system^29,49^. To characterize temporal immune response in the lung, we utilized an integrated strategy combining bulk RNA sequencing (bRNA-seq) and single-cell RNA-seq (scRNA-seq) before (0 d) and after (1, 7, and 14 d) single-dose immunization of YM3.7 (Fig. 8a). An enrichment-mixing strategy for collecting lung cells was implemented for RNA-seq: CD45^+^ immune cells and CD45^-^ cells were sorted from the lung single-cell suspensions and recombined at a 9:1 ratio^50^.

**Fig. 8.**
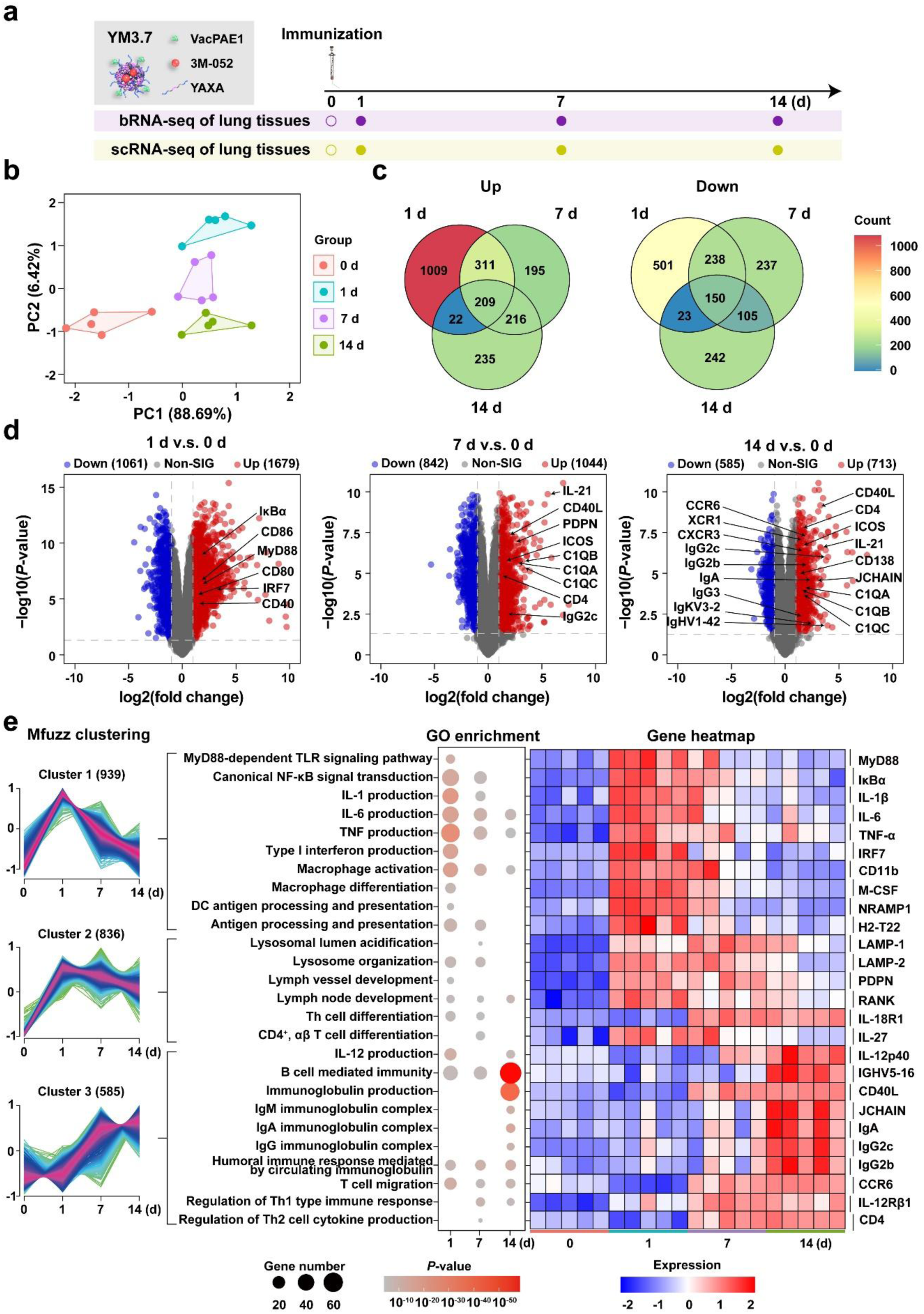
Time-course bRNA-seq transcriptomic response in the lung induced by YM3.7. **a)** Schematic diagram of experimental design. Mice are immunized with a single-dose YM3.7 (nanoparticle containing VacPAE1, 3M-052, and YAXA). 20 μg of VacPAE1, 18 μg of 3M-052, and 500 μg of YAXA are used. Lung tissues are collected on 0 d (pre-immunization) and 1/7/14 d post single-dose immunization for RNA-seq. **b)** PCA plot of sequenced samples (n=5 biologically independent replicates) at 4 timepoints. **c)** Venn diagram of DEGs. Up: up-regulated; Down: down-regulated. **d)** Volcano plots of DEGs. **e)** Time-course gene expression clusters (left) analyzed by Mfuzz, along with GO-enriched up-regulated pathways (middle) and selected up-regulated genes (right).

bRNA-seq revealed the following major results. First, principal component analysis (PCA) plot showed a high level of data reproducibility within each group and a high degree of data difference across groups (Fig. 8b). Pairwise-comparison Venn diagrams of all differentially expressed genes (DEGs) showed a total of 2197 up-regulated genes and 1496 down-regulated ones across all timepoints (Fig. 8c), illustrating the marked activation of various biological processes. Volcano plot analysis revealed distinct patterns of gene expression across the following 3 phases of immune response (Fig. 8d). At the early phase (1 d), there was significantly up-regulation of the genes encoding key TLR7/NF-κB signaling components (such as MyD88^51^, IκBα^52^, and IRF7^53^), and antigen presentation determinants (CD86, CD80, and CD40^54^). At the middle phase (7 d), there was elevated expression of GC markers (PDPN^37^), antibody secretion markers (such as IgG2c^25^), B cell activation regulators (C1QA, C1QB, and C1QC^55^), T-B cell interaction mediators (CD40L^56^ and IL-21^57^), and T cell activation markers (CD4, and ICOS^58^). At the late phase (14 d), there was sustained up-regulation of the above B cell activation regulators (C1QA, C1QB, and C1QC), T-B cell interaction mediators (CD40L and IL-21), and T cell activation markers (CD4 and ICOS). Additionally, there was newly up-regulated expression of genes encoding T cell migration receptors (CCR6^59^, CXCR3^60^, and XCR1^61^), plasma cell-associated antibody-secretion molecules (CD138^62^, JCHAIN^62^), immunoglobulin repertoire (53 heavy-chain variable regions such as IgHV1-42, and 48 light-chain variable regions such as IgKV3-2^63^), and antibody isotypes (IgA^10^, IgG2c^36^, IgG2b^64^, and IgG3^64^). Second, time-course clustering analysis categorized the 2197 up-regulated genes into 3 distinct patterns: Cluster 1 to Cluster 3 (Fig. 8e). In addition, Gene Ontology (GO) enriched pathways, along with arbitrarily selected major up-regulated genes, were built for each of these clusters to manifest their dynamic expression trends (Fig. 8e). Cluster 1 was characterized by an initial rise followed by a decline, and mainly associated with innate immunity, including TLR7/NF-κB signaling transduction as well as antigen presentation. Cluster 2 showed a sustained upward trend over time, and was primarily linked to lysosome biogenesis, lymphoid node development, and Th cell-mediated immunity. Cluster 3 was significantly up-regulated at the late phase and highly related to adaptive immunity including B cell-mediated immunity, immunoglobulin production, and regulation of Th1/Th2-type immune response. Overall, there was a temporal transition from innate immunity to adaptive immunity during the period from 1 d to 14 d post single-dose immunization of YM3.7.

We performed scRNA-seq data mining on the 46699 lung cells across 4 timepoints. This identified a total of 13 distinct cell clusters (Fig. 9a), including 6 APC subsets (cDC1, cDC2, pDC, alveolar macrophage [AM], interstitial macrophage [IM], and monocyte [Mono]), 3 lymphocyte subsets (B, T, and NK cells), neutrophil (Neu), endothelial cell (EC), and alveolar type I/II epithelial cells (AT1/2). Cell markers for clustering^65^ were detailed in Supplementary Fig. 36. Temporal dynamics of these cell types were depicted in Supplementary Fig. 37, and their overall trajectory trends were summarized in Supplementary Fig. 38. Numbers of DEGs were presented in Supplementary Fig. 39. Due to the crucial roles of APC, B cell, and T cell, scRNA-seq data mining was concentrated on these cell populations regarding their heterogeneity, dynamic proportion shifts, gene transcriptional signatures, and pathway enrichment profiles.

**Fig. 9.**
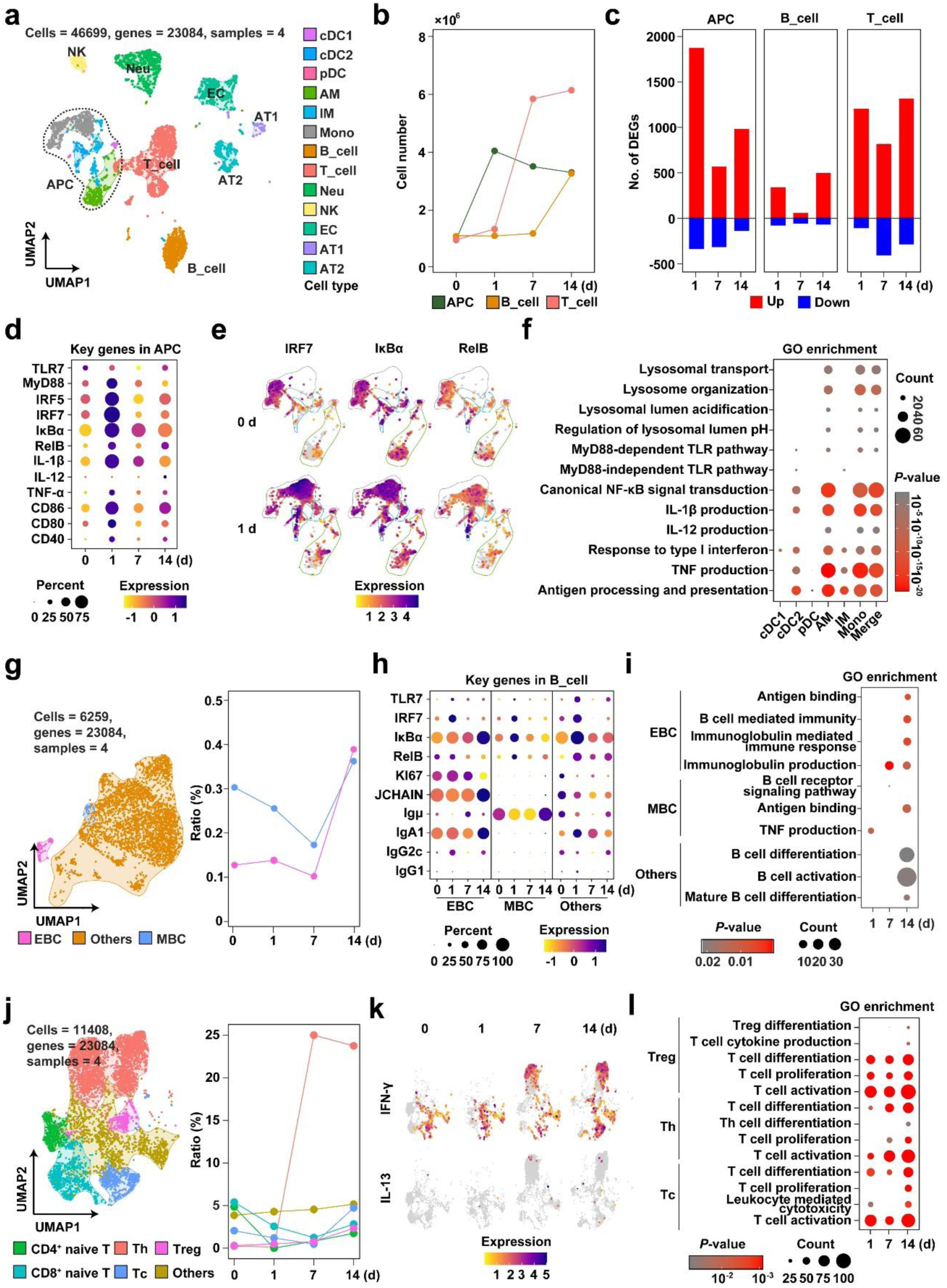
Time-course scRNA-seq transcriptomic response in the lung induced by YM3.7. Mice are immunized with a single-dose YM3.7 (nanoparticle containing VacPAE1, 3M-052, and YAXA). 20 μg of VacPAE1, 18 μg of 3M-052, and 500 μg of YAXA are used. Lung tissues are collected on 0 d (pre-immunization) and 1/7/14 d post single-dose immunization for scRNA-seq. **a)** UMAP plot of cell clusters. **b)** Line chart showing absolute counts of various cell clusters over time. **c)** Histogram showing frequencies of DEGs in various cell clusters over time. Up: up-regulated; Down: down-regulated. **d)** Dot plot showing expression levels of key TLR7/NF-κB signaling genes in APC subsets. **e)** UMAP plot showing expression patterns of IRF7, IκBα, and RelB in APC subsets on 0 d and 1 d. **f)** GO-enriched pathways for APC subsets on 1 d relative to 0 d. **g)** UMAP plot of B cell subsets (left), and line chart showing changes in proportion of B cell subsets over time (right). **h)** Dot plot showing expression levels of key genes in B cell subsets. **i)** GO-enriched pathways for B cell subsets. **j)** UMAP plot of T cell subsets (left), and line chart showing changes in proportion of T cell subsets over time. **k)** UMAP plot of expression patterns of cytokines IFN-γ and IL-13 in T cell subsets. **l)** GO-enriched pathways for T cell subsets.

First, temporal trend analysis showed that APC count peaked on 1 d and then gradually declined, while B and T cell counts steadily rose from 1 d to 14 d (Fig. 9b). DEG frequency analysis revealed that up-regulated genes in APC peaked on 1 d, whereas those in B and T cells were most pronounced on 14 d (Fig. 9c). These findings highlight the early predominance of APC activation, followed by a later shift toward adaptive B and T cell dominance.

Second, the entire APC population displayed a significant up-regulation of key TLR7/NF-κB signaling components on 1 d (Fig. 9d). A uniform transcriptional profile was confirmed across all APC subtypes (Supplementary Fig. 40). This was consistent with the above results of flow cytometry (Fig. 6b) and bRNA-seq (Fig. 8d). Temporal UMAP plots demonstrated a significant up-regulation of key TLR7/NF-κB signaling components across all APC subsets on 1 d relative to 0 d (Fig. 9e). GO-enriched up-regulated pathways primarily consisted of lysosome biogenesis, TLR7/NF-κB signaling transduction, antigen presentation (Fig. 9f). These findings delineated an integrated cascade of APC activation on 1 d, linking innate pattern recognition to adaptive immune priming.

Third, by using established specific markers (Supplementary Fig. 41), B cells were divided into the following 3 functionally distinct clusters: EBC, MBC, and other B cells (Fig. 9g). Remarkably, the proportion of EBC plus MBC showed a significant increase, reaching 0.4% of total cell count on 14 d (Fig. 9g). There was a significant up-regulation of key TLR7/NF-κB signaling components across B cell subsets (Fig. 9h), with EBC exhibiting a unique molecular signature of co-expressing cell proliferation marker Ki67^66^ and antibody-secretion-related molecules such as JCHAIN^36^, Igμ^67^, IgA1^67^, IgG2c^36^, and IgG1^36^. GO enrichment analysis revealed that EBC was primarily involved in antigen binding, B cell-mediated immunity, immunoglobulin-mediated immune response, and immunoglobulin production on 7 d and 14 d, whereas MBC was mainly associated with B cell receptor signaling pathway and antigen binding (Fig. 9i).

Fourth, by using specific markers (Supplementary Fig. 42), T cell could be classified into the following 6 distinct subsets: CD4^+^ naïve T cell, CD8^+^ naïve T cell, Th cell, Tc cell, regulatory T cell (Treg), and other T cells (Fig. 9j). Th cell exhibited a marked increase in its proportion on 7 d and 14 d, while Tc cell greatly arose on 14 d, alongside a progressive upward trend for Treg over time (Fig. 9j). Consistent with flow cytometry results (Fig. 7a-e), a significant increase in the production of cytokines IFN-γ and IL-13 was observed on 7 d and 14 d, respectively (Fig. 9k). IFN-γ and IL-13 represented the hallmarks of Th1 and Th2 response, respectively^36^. As shown by GO enrichment analysis, almost all these T cell subsets exhibited a gradual up-regulation of multiple genes responsible for differentiation, proliferation, and activation of T cell (Fig. 9l, Supplementary Fig. 43). In addition, distinct functional specializations could be observed for different T cell subsets on 14 d: Treg demonstrated Treg differentiation and T cell cytokine production, Th cell exhibited enhanced Th differentiation, while Tc cell displayed activated leukocyte-mediated cytotoxicity (Fig. 9l). This illustrated the temporal progression of functional commitments specific to various T cell subsets.

Collectively, YM3.7 orchestrated a spatiotemporally ordered induction of APC, B cell, and T cell, and moreover this process of cell activation and differentiation was specific to both the stages and the cell subsets, thereby establishing a comprehensive mechanistic framework for its superior immunoprotection efficacy.

### TLR7-stimilated immune response for YM3.7

Since 3M-052 adjuvant functioned as a TLR7 agonist^21^, we evaluated the impact of *Tlr7* knockout on immune response at both cellular (Fig. 10a) and animal (Fig. 10b) levels. First, BMDC from wild-type (WT) and *Tlr7*^-/-^ mice was treated with a total of 5 groups, namely Mock, YM3.1, YM3.4, YM3.5, and YM3.7. Following the treatments, we analyzed cell maturation, protein expression, and cytokine production using flow cytometry, Western blot, and ELISA, respectively (Fig. 10a). Compared to all other groups, both YM3.5 and YM3.7 treatments induced significantly higher levels of CD80 and CD86 (Fig. 10c), as well as key TLR7/NF-κB signaling components including p-IκBα and p-p65 (Fig. 10d) and IL-12p70 and TNF-α (Fig. 10e), in WT BMDC. CD80 and CD86 represented BMDC maturation markers^54^, while phosphorylated proteins p-IκBα and p-p65, alongside cytokine effectors IL-12p70 and TNF-α, belonged to key TLR7/NF-κB signaling components^52^. In contrast, expressions of all these proteins were markedly attenuated in *Tlr7*^-/-^ BMDC (Fig. 10c-e). Consequently, *Tlr7* deficiency in mice significantly impaired cell maturation and TLR7/NF-κB signaling transduction in BMDC treated with the above 5 groups, which was dependent on 3M-052 (but not VacPAE1 and YAXA) in relevant groups. Second, VacPAE1-specific antibody titers in sera, as well as T cell response in the lung, PMLN, and spleen, were measured by ELISA and flow cytometry, respectively, in WT and *Tlr7*^-/-^ mice received three-dose immunizations of YM3.7 (Fig. 10b). Compared to WT mice, *Tlr7*^-/-^ mice exhibited a reduced IgG titer in serum (Fig. 10f), lower frequencies of Th1/2 and Tc1/2 cells across all examined organs (Fig. 10g-i), and impaired expansion of CD4^+^/8^+^ Trm cells in the lung (Fig. 10g). Therefore, *Tlr7* deficiency in mice significantly affected VacPAE1-specific IgG production and recirculating/non-recirculating T cell response after YM3.7 immunization. Overall, the above findings highlighted the essential role of TLR7 in orchestrating YM3.7-induced immune response.

**Fig. 10.**
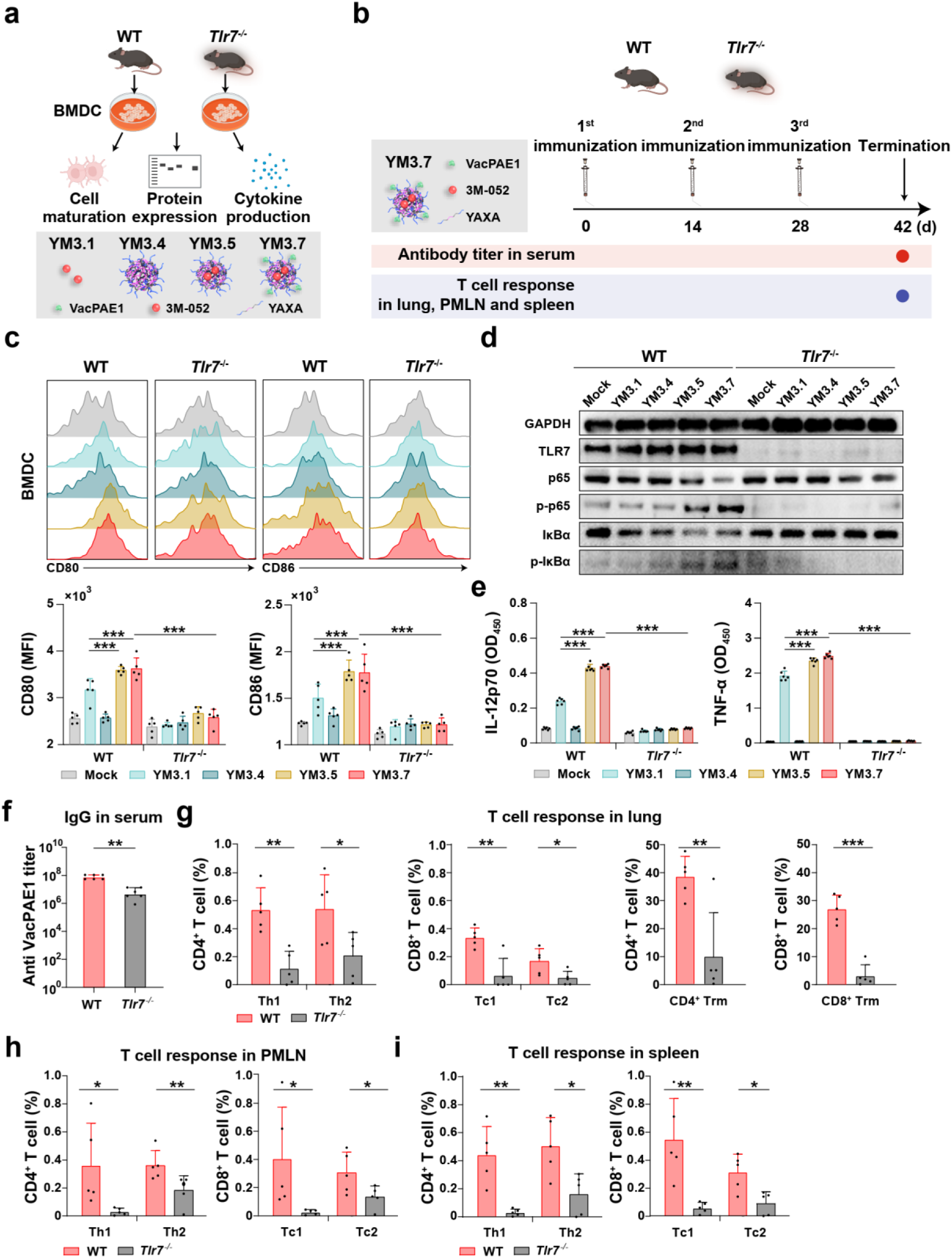
Requirement of TLR7 signaling pathway for YM3.7-induced immune response. **a)** Schematic of in vitro experimental design. BMDC isolated from WT and *Tlr7*^-/-^ mice are incubated with Mock (PBS), YM3.1 (3M-052), YM3.4 (nanoparticle containing YAXA), YM3.5 (nanoparticle containing 3M-052, and YAXA), and YM3.7 (nanoparticle containing 3M-052, YAXA, and VacPAE1). 5.56 μg/mL of VacPAE1, 138.89 μg/mL of YAXA, or 5 μg/mL 3M-052 are used for treatment. **b)** Schematic of in vivo experimental design. WT and *Tlr7*^-/-^ mice are immunized 3 times at 14-d intervals with YM3.7 (nanoparticle containing VacPAE1, 3M-052, and YAXA). 20 μg of VacPAE1, 18 μg of 3M-052, and 500 μg of YAXA are used for per inoculation. **c)** Expression of CD86 and CD80 in BMDC after 6-h incubation (n=5 biologically independent replicates). **d)** Western blot analysis of protein levels in BMDC after 3-h incubation. Experiments are repeated for 2 individual batches. **e)** ELISA measurement of cytokine levels in BMDC after 6-h incubation (n=5 biologically independent replicates). **f)** ELISA measurement of VacPAE1-specific IgG titer in sera from WT and *Tlr7*^-/-^ mice (n=6 biologically independent replicates). Flow cytometry analysis of VacPAE1-specific T cell response in the **g)** lung, **h)** PMLN and **i)** spleen from WT and *Tlr7*^-/-^ mice (n=5 biologically independent replicates). Data are presented as mean ± SD. **Statistical significance test: c, e**: YM3.7 group in WT mice is compared with each of the other groups using one-way ANOVA with Dunn’s multiple comparison test. **f-i**: Unpaired two-tailed Student’s *t*-test is used to compare WT and *Tlr7*^-/-^ groups. *: *P*<0.05, **: *P*<0.01, ***: *P*<0.001.

## 4. Conclusion

Polymeric nanoparticle vaccine (YM3.7) employed an innovative approach to modular design of mucosal vaccines by incorporating several key features: an engineered polymer backbone suitable for acid-sensitive degradation and protein conjugation, assembly of nanoparticle vaccine from polymer backbone together with antigen and adjuvant through hydrophilic-hydrophobic interactions, respective loading of antigen and adjuvant onto its surface and into its core, targeted co-delivery and subsequent co-release of antigen and adjuvant within the lysosome, and direct administration into the lung via aerosolized intratracheal inoculation. YM3.7 induced a coordinated innate, humoral, mucosal, and cell-mediated immune response in mice. In this case, cell-mediated immunity included not only recirculating B/Th/Tc cell response but also lung-resident memory B/Th/Tc cell response. The above constructed a multi-layered immune network in both lymphatic system and mucosal lung tissue. Compared to conventional antigen and adjuvant mixture (YM3.3), YM3.7 demonstrated better efficiency of accumulation within secondary lymphoid organ, uptake by APC, and intracellular trafficking to the lysosome. Once released in the lysosome, VacPAE1 was processed for MHC-II-mediated antigen presentation, while 3M-052 activated TLR7/NF-κB-dependent production of IL-12 and TNF-α. These two cytokines, in turn, enhanced APC maturation and antigen presentation. After that, there was greatly improved capacity for YM3.7 to induce multiple immune response including GC formation, recirculating B/Th/Tc cell response, and lung-resident memory B/Th/Tc cell response, and IgG/sIgA production, thereby achieving an excellent immunoprotection potency. Transcriptomic analyses revealed a spatiotemporally ordered sequence of immune response following single-dose immunization: innate immunity (such as APC activation and TLR7/NF-κB signaling transduction) peaked on 1 d, followed by adaptive immunity (GC reaction, antibody production, and B and T cell response) from 7 d to 14 d. TLR7 deficiency attenuated cell maturation and TLR7/NF-κB signaling in BMDC, and also IgG production and T cell response in mice, which was attributed to action of 3M-052 as a TLR7 agonist. The modular design property of YM3.7 would allow for substitution of antigens (e.g., viral or fungal protective antigens) and adjuvants (e.g., TLR3 or TLR9 agonists), thereby extending its potential applications from *P. aeruginosa* to diverse infectious diseases and cancers. This inhalable nanoparticle vaccine platform would pave the way for developing next-generation mucosal vaccines that offer enhanced efficacy and safety.

## Supplementary Information

Supplementary Information is available online or from the author.

## Data availability

bRNA-seq and scRNA-seq data have been deposited in the NCBI Gene Expression Omnibus repository under accession number GSE313435.

## Acknowledgements

This work is financially supported by the National Key R&D Program of China (Grant No. 2022YFC2603900).

## References

1. Curran CS, Bolig T, Torabi-Parizi P. Mechanisms and targeted therapies for Pseudomonas aeruginosa lung infection *Am J Respir Crit Care Med* **197**, 708–727 (2018).

2. Self WH, et al. Staphylococcus aureus Community-acquired Pneumonia: Prevalence, Clinical Characteristics, and Outcomes. Clin Infect Dis 63, 300–309 (2016).

3. Kulkarni D, Wang X, Sharland E, Stansfield D, Campbell H, Nair H. The global burden of hospitalisation due to pneumonia caused by Staphylococcus aureus in the under-5 years children: A systematic review and meta-analysis. EClinicalMedicine 44, 101267 (2022).

4. Torres A, et al. Adult vaccinations against respiratory infections. Expert Rev Anti Infect Ther 23, 135–147 (2025).

5. Moyle PM, Toth I. Modern subunit vaccines: development, components, and research opportunities. ChemMedChem 8, 360–376 (2013).

6. Zhu M. Immunological perspectives on spatial and temporal vaccine delivery. Adv Drug Deliv Rev 178, 113966 (2021).

7. Roth GA, Picece V, Ou BS, Luo W, Pulendran B, Appel EA. Designing spatial and temporal control of vaccine responses. Nat Rev Mater 7, 174–195 (2022).

8. Pulendran B, P SA, O’Hagan DT. Emerging concepts in the science of vaccine adjuvants. Nat Rev Drug Discov 20, 454–475 (2021).

9. Lavelle EC, Ward RW. Mucosal vaccines - fortifying the frontiers. Nat Rev Immunol 22, 236–250 (2022).

10. Mettelman RC, Allen EK, Thomas PG. Mucosal immune responses to infection and vaccination in the respiratory tract. Immunity 55, 749–780 (2022).

11. Fries CN, Curvino EJ, Chen JL, Permar SR, Fouda GG, Collier JH. Advances in nanomaterial vaccine strategies to address infectious diseases impacting global health. Nat Nanotechnol 16, 1–14 (2021).

12. Singh A. Eliciting B cell immunity against infectious diseases using nanovaccines. Nat Nanotechnol 16, 16–24 (2021).

13. Gupta A, Rudra A, Reed K, Langer R, Anderson DG. Advanced technologies for the development of infectious disease vaccines. Nat Rev Drug Discov 23, 914–938 (2024).

14. Pishesha N, Harmand TJ, Ploegh HL. A guide to antigen processing and presentation. Nat Rev Immunol 22, 751–764 (2022).

15. Kaur A, Baldwin J, Brar D, Salunke DB, Petrovsky N. Toll-like receptor (TLR) agonists as a driving force behind next-generation vaccine adjuvants and cancer therapeutics. Curr Opin Chem Biol 70, 102172 (2022).

16. Beach MA, et al. Polymeric Nanoparticles for Drug Delivery. Chem Rev 124, 5505–5616 (2024).

17. Cahn D, Amosu M, Maisel K, Duncan GA. Biomaterials for intranasal and inhaled vaccine delivery. Nat Rev Bioeng 1, 83–84 (2023).

18. Wilson JT, et al. pH-Responsive nanoparticle vaccines for dual-delivery of antigens and immunostimulatory oligonucleotides. ACS Nano 7, 3912–3925 (2013).

19. Ni Q, et al. A bi-adjuvant nanovaccine that potentiates immunogenicity of neoantigen for combination immunotherapy of colorectal cancer. Sci Adv 6, eaaw6071 (2020).

20. Qiu C, et al. Advanced strategies for overcoming endosomal/lysosomal barrier in nanodrug delivery. Research 6, 0148 (2023).

21. Kasturi SP, et al. 3M-052, a synthetic TLR-7/8 agonist, induces durable HIV-1 envelope-specific plasma cells and humoral immunity in nonhuman primates. Sci Immunol 5, eabb1025 (2020).

22. Kanamala M, Wilson WR, Yang M, Palmer BD, Wu Z. Mechanisms and biomaterials in pH-responsive tumour targeted drug delivery: A review. Biomaterials 85, 152–167 (2016).

23. Kalkhof S, Sinz A. Chances and pitfalls of chemical cross-linking with amine-reactive N-hydroxysuccinimide esters. Anal Bioanal Chem 392, 305–312 (2008).

24. Chen F, et al. Treatment of Acute Wound Infections by Degradable Polymer Nanoparticle with a Synergistic Photothermal and Chemodynamic Strategy. Adv Sci (Weinh) 11, e2309624 (2024).

25. Nazeri S, Zakeri S, Mehrizi AA, Sardari S, Djadid ND. Measuring of IgG2c isotype instead of IgG2a in immunized C57BL/6 mice with Plasmodium vivax TRAP as a subunit vaccine candidate in order to correct interpretation of Th1 versus Th2 immune response. Exp Parasitol 216, 107944 (2020).

26. Simonis A, et al. Discovery of highly neutralizing human antibodies targeting Pseudomonas aeruginosa. Cell 186, 5098–5113 e5019 (2023).

27. Boero E, et al. A flow cytometry-based assay to determine the ability of anti-Streptococcus pyogenes antibodies to mediate monocytic phagocytosis in human sera. J Immunol Methods 528, 113652 (2024).

28. Ledger EVK, Edwards AM. Host-induced cell wall remodeling impairs opsonophagocytosis of Staphylococcus aureus by neutrophils. mBio 15, e0164324 (2024).

29. Iyer H, et al. Mediastinal lymphadenopathy: a practical approach. Expert Rev Respir Med 15, 1317–1334 (2021).

30. Bronte V, Pittet MJ. The spleen in local and systemic regulation of immunity. Immunity 39, 806–818 (2013).

31. Randall TD, Carragher DM, Rangel-Moreno J. Development of secondary lymphoid organs. Annu Rev Immunol 26, 627–650 (2008).

32. Yin Q, et al. A TLR7-nanoparticle adjuvant promotes a broad immune response against heterologous strains of influenza and SARS-CoV-2. Nat Mater 22, 380–390 (2023).

33. Victora GD, Nussenzweig MC. Germinal Centers. Annu Rev Immunol 40, 413–442 (2022).

34. Ye T, et al. Inhaled SARS-CoV-2 vaccine for single-dose dry powder aerosol immunization. Nature 624, 630–638 (2023).

35. Gillespie EJ, et al. Selective inhibitor of endosomal trafficking pathways exploited by multiple toxins and viruses. Proc Natl Acad Sci U S A 110, E4904–4912 (2013).

36. Li C, et al. Mechanisms of innate and adaptive immunity to the Pfizer-BioNTech BNT162b2 vaccine. Nat Immunol 23, 543–555 (2022).

37. Aguzzi A, Kranich J, Krautler NJ. Follicular dendritic cells: origin, phenotype, and function in health and disease. Trends Immunol 35, 105–113 (2014).

38. Martinez-Riano A, et al. Long-term retention of antigens in germinal centers is controlled by the spatial organization of the follicular dendritic cell network. Nat Immunol 24, 1281–1294 (2023).

39. De Silva NS, Klein U. Dynamics of B cells in germinal centres. Nat Rev Immunol 15, 137–148 (2015).

40. Zuccarino-Catania GV, et al. CD80 and PD-L2 define functionally distinct memory B cell subsets that are independent of antibody isotype. Nat Immunol 15, 631–637 (2014).

41. Allie SR, et al. The establishment of resident memory B cells in the lung requires local antigen encounter. Nat Immunol 20, 97–108 (2019).

42. Onodera T, et al. Memory B cells in the lung participate in protective humoral immune responses to pulmonary influenza virus reinfection. Proc Natl Acad Sci U S A 109, 2485–2490 (2012).

43. Inoue T, Kurosaki T. Memory B cells. Nat Rev Immunol 24, 5–17 (2024).

44. Nutt SL, Hodgkin PD, Tarlinton DM, Corcoran LM. The generation of antibody-secreting plasma cells. Nat Rev Immunol 15, 160–171 (2015).

45. Samiea A, Celis G, Yadav R, Rodda LB, Moreau JM. B cells in non-lymphoid tissues. Nat Rev Immunol 25, 483–496 (2025).

46. D’Agostino MR, Afkhami S, Kang A, Marzok A, Miller MS, Xing Z. Protocol for isolation and characterization of lung tissue resident memory T cells and airway trained innate immunity after intranasal vaccination in mice. STAR Protoc 3, 101652 (2022).

47. Mao T, et al. Unadjuvanted intranasal spike vaccine elicits protective mucosal immunity against sarbecoviruses. Science 378, eabo2523 (2022).

48. Christo SN, Park SL, Mueller SN, Mackay LK. The multifaceted role of tissue-resident memory T cells. Annu Rev Immunol 42, 317–345 (2024).

49. Trivedi A, Reed HO. The lymphatic vasculature in lung function and respiratory disease. Front Med (Lausanne) 10, 1118583 (2023).

50. Lee A, et al. A molecular atlas of innate immunity to adjuvanted and live attenuated vaccines, in mice. Nat Commun 13, 549 (2022).

51. Deguine J, Barton GM. MyD88: a central player in innate immune signaling. F1000Prime Rep 6, 97 (2014).

52. Liu T, Zhang L, Joo D, Sun SC. NF-kappaB signaling in inflammation. Signal Transduct Target Ther 2, 17023- (2017).

53. Honda K, et al. IRF-7 is the master regulator of type-I interferon-dependent immune responses. Nature 434, 772–777 (2005).

54. Van Gool SW, Vandenberghe P, de Boer M, Ceuppens JL. CD80, CD86 and CD40 provide accessory signals in a multiple-step T-cell activation model. Immunol Rev 153, 47–83 (1996).

55. Kouser L, et al. Emerging and Novel Functions of Complement Protein C1q. Front Immunol 6, 317 (2015).

56. Karnell JL, Rieder SA, Ettinger R, Kolbeck R. Targeting the CD40-CD40L pathway in autoimmune diseases: Humoral immunity and beyond. Adv Drug Deliv Rev 141, 92–103 (2019).

57. Yoshizaki A, et al. Regulatory B cells control T-cell autoimmunity through IL-21-dependent cognate interactions. Nature 491, 264–268 (2012).

58. Tafuri A, et al. ICOS is essential for effective T-helper-cell responses. Nature 409, 105–109 (2001).

59. Yamazaki T, et al. CCR6 regulates the migration of inflammatory and regulatory T cells. J Immunol 181, 8391–8401 (2008).

60. Hickman HD, et al. CXCR3 chemokine receptor enables local CD8(+) T cell migration for the destruction of virus-infected cells. Immunity 42, 524–537 (2015).

61. Dorner BG, et al. Selective expression of the chemokine receptor XCR1 on cross-presenting dendritic cells determines cooperation with CD8+ T cells. Immunity 31, 823–833 (2009).

62. Scharer CD, Patterson DG, Mi T, Price MJ, Hicks SL, Boss JM. Antibody-secreting cell destiny emerges during the initial stages of B-cell activation. Nat Commun 11, 3989 (2020).

63. Glanville J, et al. Precise determination of the diversity of a combinatorial antibody library gives insight into the human immunoglobulin repertoire. Proc Natl Acad Sci U S A 106, 20216–20221 (2009).

64. Roco JA, et al. Class-Switch Recombination Occurs Infrequently in Germinal Centers. Immunity 51, 337–350 e337 (2019).

65. Su D, et al. Spatiotemporal single-cell transcriptomic profiling reveals inflammatory cell states in a mouse model of diffuse alveolar damage. Exploration (Beijing) 3, 20220171 (2023).

66. Coffey F, Alabyev B, Manser T. Initial clonal expansion of germinal center B cells takes place at the perimeter of follicles. Immunity 30, 599–609 (2009).

67. Macpherson AJ, et al. IgA production without mu or delta chain expression in developing B cells. Nat Immunol 2, 625–631 (2001).

