## Supplementary Information for "Inhalable Polymeric Nanoparticle Vaccine for Lysosome-targeting Co-delivery of Antigen and Adjuvant with Enhanced Immunoprotection"

**Experimental section**

**Materials.** Terephthalaldehyde (Cat# 623-27-8, Aladdin, China), 2-aminoethanol hydrochloride (Cat# A110257, Aladdin, China), L-lysine diisocyanate (Cat# L857594, Aladdin, China), 3M-052 (Cat# HY-19573, MedChemExpress, USA), anhydrous methanol (Cat# CM060050, Concord, China), N,N-dimethylformamide (DMF, Cat# CD061010, Concord, China), dimethyl sulfoxide (DMSO, Cat# CD061015, Concord, China), PBS (Cat# 70011044, Thermo Fisher Scientific, USA), and dialysis membranes (3.5 kDa: Cat# AC131192; 8-14 kDa: Cat# AC131276; 100 kDa: Cat# AC131408, Thermo Fisher Scientific, USA) were used for nanoparticle preparation. Reagents for measuring protein loading included Pierce BCA Protein Assay Kit (Cat# 23225, Thermo Fisher Scientific, USA). Cell culture reagents included NIH-3T3 cell line (Cat# CRL-1658, American Type Culture Collection, USA), BEAS-2B cell line (Cat# CRL-3588, American Type Culture Collection, USA), RAW264.7 cell line (Cat# TIB-71, American Type Culture Collection, USA), Dulbecco's Modified Eagle Medium (DMEM, Cat# 10-013-CV, Corning, USA), RPMI 1640 medium (Cat# 10-040-CV, Corning, USA), fetal bovine serum (FBS, Cat# 10437028, Thermo Fisher Scientific, USA), penicillin-streptomycin (Cat# P4333, Sigma-Aldrich, USA), trypsin-EDTA (Cat# T4049, Sigma-Aldrich, USA), L-glutamine (Cat# G8540, Sigma-Aldrich, USA), β-mercaptoethanol (Cat# M6250, Sigma-Aldrich, USA), recombinant mouse GM-CSF (Cat# 415-ML, R&D Systems, USA), CCK-8 kit (Cat# HY-K0301, MedChemExpress, USA), meropenem (Cat# HY-13678, MedChemExpress, USA), cefepime (Cat# HY-B0692, MedChemExpress, USA), methicillin (Cat# HY-121544, MedChemExpress, USA), and 4% paraformaldehyde fixative (Cat# G1102, Servicebio, China). Bacterial culture utilized Brain Heart Infusion broth (Cat# 211065, BD Biosciences, USA). Detection of antibody titer employed high-binding 96-well plates (Cat# 1030011, Dakewe, China), TMB substrate (Cat# DKW-TMB-0020, Dakewe, China), Stop solution (Cat# DKW-SULF-0010, Dakewe, China), and HRP-conjugated anti-mouse antibodies (IgG: Cat# ab6561204; IgA: Cat# ab97226; IgG1: Cat# ab6736056; IgG2c: Cat# ab6736058, Abcam, UK). Reagents for fluorescence imaging included Sulfo-Cy5.5 NHS ester (Cy5.5, Cat# CY5.5-001, Qiyue Xi'An, China), LysoTracker Green (Cat# L7526, Thermo Fisher Scientific, USA), and Hoechst 33342 (Cat# H1399, Nuobold, China). 4',6-diamidino-2-phenylindole (DAPI, Cat# G1012, Servicebio, China), anti-B220 rabbit pAb (Cat# GB113886, Servicebio, China), anti-Ki67 rabbit pAb (Cat# GB111141, Servicebio, China), CFSE (Cat# HY-D0938, MedChemExpress, USA), HRP conjugated Goat Anti-Rabbit IgG (H+L) (Cat# GB23303, Servicebio, China), Tyramide Signal Amplification (TSA, Cat# G1226, Servicebio, China). Reagents for flow cytometry comprised 100-μm strainer (Cat# CLS431752, Corning, USA), collagenase type IV (Cat# LS004188, Worthington, USA), Fixable Viability Dye 780 (FVD780, Cat# 565388, BD Biosciences, USA), Fc-blocker (Clone 2.4G2, Cat# 553141, BD Biosciences, USA), and Cytofix/CytopermTM Fixation and Permeabilization buffer (Cat# 554714, BD Biosciences, USA), ethylenediaminetetraacetic acid (EDTA, Cat# 20158, Sigma-Aldrich, USA), sodium azide (Cat# S2002, Sigma-Aldrich, USA). Western blotting reagents were categorized as follows: electrophoresis and transfer reagents, membranes and transfer reagents, buffers and blocking agents, protease inhibitors and detection, primary antibodies, and secondary antibodies. Electrophoresis and transfer reagents included Tris-glycine gels (Cat# PG113, Yaozyme, China), PAGE gel rapid preparation kit (Cat# PG213, Epizyme Biotech, China), SDS-PAGE protein buffer (Cat# P0015, Beyotime, China), SDS-PAGE sample loading buffer (Cat# P0015, Beyotime, China), and prestained protein ladder (Cat# 26616, Thermo Fisher Scientific, USA). Membranes and transfer reagents included PVDF membranes (Cat# IPVH00010, Merck, Germany), fast transfer powder (Cat# G2028, Servicebio, China). Buffers and blocking agents included cell lysis buffer for Western/IP (Cat# P0013, Beyotime, China), Tris-buffered saline (Cat# G0001-2L, Servicebio, China), blocking buffer (Cat# P0023B, Beyotime, China), primary antibody dilution buffer (Cat# P0023A, Beyotime, China), tween-20 (Cat# ST825, Beyotime, China). Protease inhibitors and detection included protease/phosphatase inhibitor cocktail (Cat# ab201119 and ab141272, Abcam, UK), chemiluminescent HRP substrate kit (Cat# P06M31, Gene-Protein Link, China). Primary antibodies included TLR7 (Cat# 82658, Cell Signaling Technology, USA), IκBα (Cat# 9242S, Cell Signaling Technology, USA), p-IκBα (Cat# 2859S, Cell Signaling Technology, USA), p65 (Cat# 6956S, Cell Signaling Technology, USA), p-p65 (Cat# 3033S, Cell Signaling Technology, USA), GAPDH (Cat# 60004-1, Proteintech, USA). Secondary antibodies included Goat Anti-Rabbit IgG-HRP (Cat# P03S02, Gene-Protein Link, China) and Goat Anti-Mouse IgG-HRP (Cat# P03S01, Gene-Protein Link, China). Reagents for measuring cytokine expression comprised ElaBoXTM Mouse IL-12p70 (Cat# SEKM-0013, Solarbio, China) and TNF-α (Cat# SEKM-0034, Solarbio, China). Reagents for bRNA-seq and scRNA-seq sample preparation involved biotinylated anti-CD45 (Cat# 130-097-153, Miltenyi Biotec, Germany), streptavidin-labeled magnetic beads (Cat# 130-048-101, Miltenyi Biotec, Germany), bovine serum albumin (BSA, Cat# A9418, Sigma-Aldrich, USA), and RNase inhibitor (Cat# R1158, Sigma-Aldrich, USA).

**Synthesis of monomer M1.** Terephthalaldehyde (134 mg) and 2-aminoethanol hydrochloride (194 mg) were dissolved in 15 mL of anhydrous methanol. The mixture was stirred at room temperature for 3 h, during which a precipitate gradually formed. The precipitate was collected by centrifugation and washed with anhydrous methanol. The monomer M1 was obtained after drying the precipitate under vacuum. $^{1}H$ NMR and $^{13}C$ NMR (400 MHz, DMSO-$d_6$) was shown in Supplementary Fig. 2.

**Synthesis of polymer YAXA.** Monomers M1 (220 mg) and M2 (249 mg) were dissolved in 10 mL of anhydrous DMF. The mixture was stirred at room temperature for 24 h. Subsequently, monomer M3 (1 g) was dissolved in 10 mL of anhydrous DMF and added to the reaction mixture, which was then stirred for an additional 24 h. The reaction mixture was purified by dialysis using a dialysis membrane with a molecular weight cutoff (MWCO) of 8-14 kDa for 48 h. Finally, YAXA was obtained as a white solid by freeze-drying. $^{1}H$ NMR (400 MHz, DMSO-$d_6$) was shown in Supplementary Fig. 3.

**Preparation of VacPAE1 and VacSAU4 protein antigens.** VacPAE1 was designed as a chimeric antigen comprising the following protein fragments (amino acid positions denoted by subscripts) from *P. aeruginosa*: $PcrV_{1-127}$ (NP_250397, 13.9 kDa)[1], $PcrV_{251-294}$ (NP_250397, 5.2 kDa)[1], $OprF_{311-341}$ (NP_250468, 3.4 kDa)[2], $OprI_{25-83}$ (NP_251543, 6.6 kDa)[3], full-length Hcp1 (NP_248775, 17.4 kDa)[3]. VacSAU4 was designed as a chimeric antigen containing 2 distinct attenuated toxins from *S. aureus*: $SpA_{KKAA}$[4] and $Hla_{H35L/H48L}$[5]. Both VacPAE1 and VacSAU4 were recombinantly expressed in *Escherichia coli* BL21(DE3) using pET28a vector[6]. VacPAE1 construct included the following sequential elements: His-SUMO tag, $PcrV_{1-127}$, GGGGS linker, $PcrV_{251-294}$, GGGGSGGGG linker, $OprF_{311-341}$, GGGGSGGGG linker, $OprI_{25-83}$, GSGGSG linker, and Hcp1. VacSAU4 construct contained: His-SUMO tag, $SpA_{KKAA}$, GGGGSGGGGSGGGGS linker, and $Hla_{H35L/H48L}$. After induction with isopropyl β-D-1-thiogalactopyranoside (0.5 mM, 15°C, 16 h), the cells were lysed by sonication, and then the lysate was purified sequentially through ion-exchange chromatography (Q column, 0-1 M NaCl gradient) and Ni-NTA affinity chromatography (500 mM imidazole elution). Following affinity purification, an endotoxin removal step was performed using a specialized endotoxin-removing resin to minimize potential pyrogen contamination. His-SUMO tag was cleaved overnight at 4°C with SUMO protease. Untagged VacPAE1 or VacSAU4 protein was isolated via reverse Ni-NTA. Final polishing used Superdex75 gel filtration, followed by dialysis into PBS with 300 mM NaCl/10% glycerol, sterilization, and quantification by BCA assay. Endotoxin levels were measured using Limulus Amebocyte Lysate assay, confirming a final concentration below the detection limit of 0.25 EU/mL, substantially lower than the accepted safety standard[7].

**Pre-experiment to measure 3M-052 loading during YM3.5 formulation optimization.** 3M-052 concentration was fixed at 360 μg/mL. YAXA:3M-052 mass ratios were set at 15:1, 20:1, 25:1, 27.78:1, 30:1, 35:1, and 40:1. Accordingly, various amounts of YAXA were co-dissolved with 3M-052 (2.4 mg) in DMSO (400 μL) via sonication. Each mixture was added dropwise into 3.6 mL of PBS under vigorous stirring. Purified YM3.5 was obtained after dialyzing for 24 h against PBS using a 3.5 kDa MWCO membrane, and final volume was adjusted to 6.67 mL. 3M-052 concentrations were quantified by 1260 Infinity II HPLC (Agilent, Germany) equipped with C18 Reverse-phase Column, and loading efficiencies and loading contents of 3M-052 were calculated (Supplementary Fig. 4).

**Pre-experiment to measure VacPAE1 loading during YM3.7 formulation optimization.** 3M-052 concentration and YAXA:3M-052 mass ratio was fixed at 360 μg/mL and 27.78:1, respectively. To measure loading capacities at varying amounts of VacPAE1, 600 μL of YM3.5 (16.668 mg/mL YAXA and 600 μg/mL 3M-052) was mixed with 80, 160, or 320 μL of VacPAE1 solution (2.5 mg/mL), and the volume was adjusted to 1 mL with PBS. Each mixture was stirred at room temperature for 3 h, then dialyzed against PBS using a 100 kDa MWCO membrane to remove unbound VacPAE1, and final volume was adjusted to 1 mL. Then, 50 μL of each nanoparticle suspension was incubated with 200 μL of BCA working reagent, and absorbance at 562 nm was recorded on SpectraMax i3x Microplate Reader (Molecular Devices, USA), followed by calculation of VacPAE1 loading contents and loading efficiencies (Supplementary Fig. 5).

**Preparation of final-version nanoparticle YM3.4/5/6/7.** YAXA (66.672 mg) was fully dissolved in 400 μL of DMSO by bath sonication. The solution was then added dropwise to vigorously stirred PBS (3.6 mL). Residual DMSO was removed by dialysis using a membrane with a MWCO of 3.5 kDa for 24 h, yielding YM3.4 (nanoparticle containing pure

YAXA). YAXA (66.672 mg) and 3M-052 (2.4 mg) were co-dissolved in 400 μL of DMSO through bath sonication. The resulting mixture was slowly dripped into vigorously stirred PBS (3.6 mL). Subsequent dialysis with the same conditions (3.5 kDa, 24 h) eliminated residual DMSO, resulting in YM3.5 (nanoparticle containing YAXA and 3M-052). A mixture of 178 μL of VacPAE1 (2.5 mg/mL), 800 μL of YM3.4 (13.889 mg/mL YAXA), and 24 μL of PBS was stirred at room temperature for 3 h. Unbound protein was separated via dialysis with a MWCO of 100 kDa, yielding YM3.6. A mixture of 178 μL of VacPAE1 (2.5 mg/mL), 800 μL of YM3.5 (13.889 mg/mL YAXA and 500 μg/mL 3M-052), and 24 μL of PBS was stirred at room temperature for 3 h. Dialysis with a MWCO of 100 kDa was performed to remove excess protein, generating YM3.7 (nanoparticle containing YAXA, 3M-052, and VacPAE1). To achieve fluorescence labeling, VacPAE1, 3M-052, YAXA, and Cy5.5 were co-assembled to generate nanoparticle YM3.7-Cy5.5, followed by dialysis to remove excess dye, while VacPAE1, 3M-052, and Cy5.5 are directly mixed to produce YM3.3-Cy5.5. The concentration of Cy5.5 was determined by absorbance at its specific maximum wavelength 695 nm using SpectraMax i3x Microplate Reader. The block concentrations were as follows: YAXA at 10 mg/mL, 3M-052 at 360 μg/mL, VacPAE1 at 400 μg/mL, and Cy5.5 at 20 μg/mL.

**Characterizations.** 5 mg monomer M1 or polymer YAXA was dissolved in deuterated DMSO, and $^{13}$C NMR and $^{1}$H NMR spectra were subsequently measured using AVANCE NEO 400 NMR spectrometer (Bruker, Germany) at room temperature. A maximum of 6 experimental groups were as follows: YM3.2 (VacPAE1), YM3.3 (mixture of VacPAE1 and 3M-052), YM3.4 (nanoparticle containing pure YAXA), YM3.5 (nanoparticle containing YAXA and 3M-052), YM3.6 (nanoparticle containing YAXA and VacPAE1), and YM3.7 (nanoparticle containing YAXA, 3M-052, and VacPAE1). TEM and SEM images of YM3.4/5/6/7 solution, containing 100 μg/mL YAXA, 3.6 μg/mL 3M-052, or 4 μg/mL VacPAE1, were acquired using HT-7700 TEM (Hitachi, Japan) and JEM-ARM200F SEM (Hitachi, Japan), respectively. YM3.4/5/6/7 solution was used at its block concentration for the following experiments. To quantify hydrodynamic diameter and zeta potential, 1 mL of YM3.4/5/6/7 solution was analyzed using Zetasizer Nano ZS90 (Malvern, England). To determine 3M-052 concentration, 100 μL of YM3.5/7 solution was analyzed by HPLC. To determine VacPAE1 concentration, 50 μL of YM3.2/3/6/7 solution was mixed with 200 μL of BCA working reagent and then quantified at 562 nm using SpectraMax i3x Microplate Reader. To verify VacPAE1's structural integrity, 10 μL of YM3.2/3/6/7 solution was resolved on 12.5% Tris-glycine gel under reducing conditions, followed by Coomassie Blue staining for band intensity quantification. To assess physiological stability in murine blood, 100 μL of YM3.7 solution was incubated with 900 μL of PBS containing 10% FBS and then subjected to hydrodynamic size monitoring over 24 h to measure in vitro stability.

**Preparation and characterization of YM3.7 loaded with VacSAU4.** A maximum of 6 experimental groups were as follows: YM3.2 (VacSAU4), YM3.3 (mixture of VacSAU4 and 3M-052), YM3.4 (nanoparticle containing pure YAXA), YM3.5 (nanoparticle containing YAXA and 3M-052), YM3.6 (nanoparticle containing YAXA and VacSAU4), and YM3.7 (nanoparticle containing YAXA, 3M-052, and VacSAU4). YM3.2/3/4/5/6/7 formulation was prepared with the following component concentrations: YAXA at 10 mg/mL, 3M-052 at 360 μg/mL, and VacSAU4 at 400 μg/mL. To quantify hydrodynamic diameter and zeta potential, 1 mL of YM3.4/5/6/7 solution was analyzed using Zetasizer Nano ZS90. To determine VacSAU4 concentration, 50 μL of YM3.2/3/6/7 solution was mixed with 200 μL of BCA working reagent and then quantified at 562 nm using SpectraMax i3x Microplate Reader. To verify VacSAU4's structural integrity, 10 μL of YM3.2/3/6/7 solution was resolved on 12.5% Tris-glycine gel under reducing conditions, followed by Coomassie Blue staining for band intensity quantification.

**Measurement of nanoparticle dissociability triggered by acidic conditions.** There were the following 3 experimental groups: YM3.5 (nanoparticle containing YAXA and 3M-052), YM3.6 (nanoparticle containing YAXA and VacPAE1), and YM3.7 (nanoparticle containing YAXA, 3M-052, and VacPAE1). The working concentrations were as follows: YAXA at 1 mg/mL, 3M-052 at 36 μg/mL, and VacPAE1 at 40 μg/mL. 1 mL of YM3.5/6/7 solution was dialyzed in pH 5.0/7.4 PBS using dialysis tube with a MWCO of 100 kDa, under gentle agitation at 100 rpm. To quantify 3M-052 release, 50 μL of YM3.5/3.7 solution collected at various time intervals (0, 12, 24, 36, 48, 60, and 72 h) was analyzed by HPLC. To quantify VacPAE1 release, 50 μL of YM3.6/7 solution was mixed with 200 μL of BCA working reagent, and then subjected to BCA assay. To assess hydrodynamic diameter, 1 mL of YM3.7 solution was analyzed by DLS after 72-h dialysis at pH 5.0/7.4. To evaluate morphological changes, 1 mL of YM3.7 solution was subjected to SEM.

**Determination of cell cytotoxicity of nanoparticle YM3.4.** To assess cell cytotoxicity of YM3.4 (nanoparticle containing pure YAXA), a total of 3 different cell types were employed: NIH-3T3 cell line (https://www.atcc.org/products/crl-1658), BEAS-2B cell line (https://www.atcc.org/products/crl-3588), and BMDC. NIH-3T3 and BEAS-2B cells were cultured in DMEM supplemented with 10% FBS, 100 U/mL penicillin-streptomycin, and 2 mM L-glutamine at 37°C under 5% $CO_2$. When reaching 80-90% confluence, they were detached using 0.25% trypsin-EDTA, washed, and prepared for further use. BMDC was isolated from WT mice and maintained in RPMI 1640 medium containing 10% FBS, 100 U/mL penicillin-streptomycin, 50 μM β-mercaptoethanol, and 10 ng/mL recombinant mouse GM-CSF. Fresh medium (10 mL) was added on 3 d, and half of the medium was replaced on 6 d. On 8 d, the medium was gently pipetted, and immature BMDC that was suspended in the medium and loosely adhered to the flask was collected for further use. All 3 cell types were seeded in 96-well plates at a density of $1\times10^4$ cells per well, and allowed to adhere for 24 h. YM3.4 was then added at 0, 50, 100, 250, or 500 μg/mL. After 24-h treatment, the medium was removed, and 100 μL of fresh medium containing 10% CCK-8 solution was added to each well. Following 4-h incubation, cell viability was quantified by measuring absorbance at 450 nm using SpectraMax i3x Microplate Reader.

**Determination of hemolytic activity of nanoparticle YM3.4.** To evaluate hemolytic activity of YM3.4 (nanoparticle containing pure YAXA), blood samples were collected from WT mice via retro-orbital bleeding and centrifuged at 350 g for 10 min to isolate red blood cell. The harvested red blood cell samples were subsequently washed and resuspended in

PBS to a final concentration of $2\times10^6$ cells per mL. For hemolysis assessment, 100 μL red blood cell suspension was mixed with 900 μL PBS containing YM3.4 at concentrations of 50, 100, 250, or 500 μg/mL, establishing the corresponding 4 experimental groups. Deionized water and PBS served as positive and negative controls, respectively. After 4-h incubation at room temperature, samples were centrifuged at 3000 rpm for 10 min, and hemolysis was visually documented. Supernatants were further centrifuged at 11000 rpm for 10 min, and absorbance at 550 nm was measured using SpectraMax i3x Microplate Reader. The hemolysis rate was calculated using the following formula: hemolysis rate % = (experimental group - negative control) / (positive control - negative control) ×100%.

**Single-dose or three-dose immunizations in mice.** To achieve aerosolized intratracheal inoculation, each liquid formulation (50 μL) was delivered into the lung of each mouse using MicroSprayer Aerosolizer (Huironghe, China). For single-dose immunization, mice were immunized only once. For three-dose immunizations, mice were immunized 3 times at 14-d intervals. 20 μg of VacPAE1, 18 μg of 3M-052, 500 μg of YAXA, or 1 μg of Cy5.5 were used per inoculation.

**Determination of biosafety of nanoparticle YM3.7 in mice.** WT mice were subjected to single-dose immunization with the following 2 groups: Mock (PBS), and YM3.7 (nanoparticle containing VacPAE1, 3M-052, and YAXA). On 7 d and 28 d post single-dose immunization, mice were euthanized, and the tissues of the lung, spleen, heart, kidney, and liver were harvested and fixed in 4% paraformaldehyde. Fixed tissues were dehydrated through a graded ethanol series (70%, 80%, 95%, and 100%), cleared in xylene, and embedded in paraffin. Sections (4 μm) were stained with H&E and then examined using DM500 Light Microscope (Leica, Germany) for pathological analysis. Blood samples were collected from mice on 7 d post single-dose immunization as above and then centrifuged at 1000 rpm for 10 min, and serum was analyzed for hepatic function markers (alanine aminotransferase and aspartate aminotransferase) and renal function markers (blood urea nitrogen and creatinine) by Compass Biotechnology (China).

**Determination of VacPAE1-specific antibody production in mice.** WT mice received three-dose immunizations in 2 independent experiments. In the first experiment, mice were randomly divided into 3 groups: YM3.2 (VacPAE1), YM3.3 (mixture of VacPAE1 and 3M-052), and YM3.7 (nanoparticle containing VacPAE1, 3M-052, and YAXA). In the second experiment, the following 2 groups were compared: YM3.7 and YM3.5+YM3.6 [physical mixture of YM3.5 (nanoparticle containing YAXA and 3M-052) and YM3.6 (nanoparticle containing YAXA and VacPAE1)]. Blood samples were collected via the tail vein route on 7, 14, 21, 28, 35, and 42 d post the first immunization for the first experiment, and on 14, 28, and 42 d post the first immunization for the second experiment. To promote coagulation, whole blood sample was incubated at room temperature for 30 min, followed by centrifugation at 3000 g for 10 min to isolate serum. On 14, 28, and 42 d post the first immunization for both experiments, BALF was collected by perfusing each mouse with ice-cold PBS, followed by intratracheal lavage with 1 mL of PBS containing 0.5% BSA. Both serum and BALF samples were stored at -20°C for subsequent analysis. Titers of anti-

VacPAE1 antibody subsets, including IgG, IgG2c, and IgG1 in serum, along with IgG and IgA in BALF, were quantified by ELISA. High-binding 96-well plates were coated overnight with 0.1 μg of VacPAE1 per well. After treatment with blocking buffer for 2 h, diluted samples were incubated for 30 min. Bound antibodies were detected using horseradish peroxidase-conjugated secondary antibodies specific for mouse IgG, IgA, IgG1, or IgG2c, respectively. Reactions were developed with 50 μL of TMB substrate, followed by treatment with 50 μL of Stop solution, and absorbance at 450 nm was measured using SpectraMax i3x Microplate Reader.

***P. aeruginosa* strain and cultivation.** Two *P. aeruginosa* strains were used in this study: the multidrug-resistant clinical isolate F291007 (serotype O11, ST235 clone)[8, 9], which displayed a hypervirulent phenotype and was associated with global nosocomial infections, and the laboratory reference strain PAO1 (serotype O5)[10]. Both strains were cultured using a standardized three-step protocol. For F291007, primary culture (F1) was initiated by inoculating 20 μL of bacterial stock into 20 mL of BHI broth supplemented with 4 μg/mL meropenem, followed by incubation at 37 °C for 15 h with shaking at 200 rpm to reach stationary phase. Subsequently, 100 μL of F1 culture was transferred into 20 mL of fresh meropenem-containing BHI broth and incubated for 4 h under identical conditions to obtain mid-logarithmic phase culture (F2). F2 culture was then diluted 1:100 in meropenem-free BHI broth and incubated for 3 h to generate third-generation mid-logarithmic phase culture (F3). For PAO1, the same three-step protocol was followed except that all media were meropenem-free. The optical density ($OD_{600}$) of the F3 culture was adjusted to 0.2 for F291007 (~$1\times10^7$ CFU/mL) and to 1.0 for PAO1 (~$8\times10^8$ CFU/mL) using Multiskan FC Microplate Reader (Thermo Fisher Scientific, USA) prior to experimental use.

**In vitro neutralizing activity assessment in serum and BALF against *P. aeruginosa*.** WT mice that received three-dose immunizations were divided into 3 groups: YM3.2 (VacPAE1), YM3.3 (mixture of VacPAE1 and 3M-052), and YM3.7 (nanoparticle containing VacPAE1, 3M-052, and YAXA). Serum and BALF samples were collected on 42 d post the first immunization. For neutralization assay, RAW264.7 macrophages were seeded in a 96-well plate at a density of $10^5$ cell/mL in RPMI-1640 medium supplemented with 10% FBS and cultured at 37 °C under 5% $CO_2$. In neutralization step, *P. aeruginosa* F291007 ($10^5$ CFUs) or PAO1 ($10^6$ CFUs) was pre-incubated with 1 μL serum (1:200 dilution) or 20 μL BALF (1:50 dilution) in 200 μL of RPMI-1640 medium supplemented with 10% FBS for 30 min at 4 °C to generate the pre-opsonized bacterial suspension[11]. Subsequently, cell culture supernatant was removed, and cells were infected with the pre-opsonized bacterial suspension (200 μL/well). Following 240-min incubation, cells were gently washed with PBS and then treated with 100 μL of RPMI-1640 medium containing 10% FBS and 40 μg/mL of cefepime for 18 h to kill extracellular bacteria. Cell culture supernatant was then replaced with 100 μL of fresh medium containing 10% CCK-8 solution. After 4-h incubation, cell viability was quantified by measuring absorbance at 450 nm using SpectraMax i3x Microplate Reader. The resulting cell viability served as the indicator of neutralizing activity of serum and BALF samples.

**In vitro determination of opsonophagocytic activity in serum and BALF against *P. aeruginosa*.** WT mice that received three-dose immunizations were divided into 3 groups: YM3.2 (VacPAE1), YM3.3 (mixture of VacPAE1 and 3M-052), and YM3.7 (nanoparticle containing VacPAE1, 3M-052, and YAXA). Serum and BALF samples were collected on 42 d post the first immunization. RAW264.7 macrophages were seeded in flow cytometry tubes at a density of $5\times10^5$ cell/tube in RPMI-1640 medium supplemented with 10% FBS and cultured at 37 °C under 5% $CO_2$. *P. aeruginosa* F291007 or PAO1 was stained with CFSE for 60 min and washed twice to remove excess dye[12, 13]. For each opsonophagocytosis assay, CFSE-labeled *P. aeruginosa* ($10^5$ CFUs of F291007 or $10^6$ CFUs of PAO1) was incubated with 1 μL of serum (1:200 dilution) or 20 μL of BALF (1:50 dilution), together with 1 μL of heat-inactivated serum from unimmunized mice as the complement source. The final reaction volume was adjusted to 200 μL with RPMI-1640/10% FBS. The opsonized bacterial suspension was added to the macrophage pellet and incubated for 60 min at 37 °C under 5% $CO_2$ to permit phagocytosis[12, 13]. Then, cells were fixed with 4% paraformaldehyde and washed twice with PBS to remove extracellular bacteria. Opsonophagocytic activity was quantified by measuring CFSE fluorescence intensity of the macrophage population using LSRFortessa Flow Cytometer (BD Biosciences, USA) operated by FACSDiva software (v8.0.1).

***P. aeruginosa* challenge, survival monitoring, bacterial load determination, and pathological examination in mice.** WT mice were assigned to 2 independent experiments. In the first experiment, 4 groups received three-dose immunizations as follows: Mock (PBS), YM3.2 (VacPAE1), YM3.3 (mixture of VacPAE1 and 3M-052), and YM3.7 (nanoparticle containing VacPAE1, 3M-052, and YAXA). In the second experiment, 2 groups were compared: YM3.7 and YM3.5+YM3.6 [physical mixture of YM3.5 (nanoparticle containing YAXA and 3M-052) and YM3.6 (nanoparticle containing YAXA and VacPAE1)]. On 42 d post the first immunization, each mouse was anesthetized and subjected to aerosolized intratracheal challenge with a predetermined $LD_{100}$ dose (~$1.4\times10^6$ CFUs) of *P. aeruginosa* F291007. Survival rates, as well as body weights were monitored daily for 7 d following the challenge. At 6 h post *P. aeruginosa* challenge, the blood, lungs, spleens, and livers were collected from mice of the first experimental groups to quantify bacterial load by using plate counting method. Additionally, mice from the YM3.3 and YM3.7 groups in the first experiment were euthanized at 48 h post *P. aeruginosa* challenge for pathological assessment of the lungs, spleens, and livers.

**Determination of VacSAU4-specific antibody production in mice.** WT mice were subjected to three-dose immunizations with the following 4 groups: YM3.2 (VacSAU4), YM3.3 (mixture of VacSAU4 and 3M-052), YM3.6 (nanoparticle containing VacSAU4 and YAXA), and YM3.7 (nanoparticle containing VacSAU4, 3M-052, and YAXA). Serum and BALF samples were collected on 14, 28, and 42 d post the first immunization. Titers of anti-VacSAU4 antibody subsets, including IgG in serum, along with IgG and IgA in BALF, were quantified by ELISA.

***S. aureus* strain and cultivation.** The methicillin-resistant clinical isolate *S. aureus* USA300-FPR3757[14], which belonged to the high-risk ST8/CC8 clone and was characterized by a hypervirulent phenotype associated with global nosocomial infections and poor clinical outcome, was cultured through a three-step process. For primary culture (F1), 10 μL of bacterial stock was inoculated into 20 mL of BHI broth supplemented with 5 μg/mL methicillin and incubated at 37 °C with shaking at 200 rpm for 12 h. Secondary culture (F2) was initiated by transferring 100 μL of F1 into 20 mL of fresh BHI broth containing 5 μg/mL methicillin, followed by incubation under identical conditions for 3.5 h. Then, 100 μL of F2 was inoculated into 20 mL of methicillin-free BHI broth and incubated at 37 °C with shaking at 200 rpm for 3.5 h until $OD_{600}$ reached 1.8 (F3). The $OD_{600}$ value of the F3 culture was then adjusted to 0.2 using Multiskan FC Microplate Reader, corresponding to approximately $1\times10^7$ CFU/mL for subsequent experimental use.

***S. aureus* challenge and survival monitoring in mice.** WT mice were subjected to three-dose immunizations with the following 5 groups: Mock (PBS), YM3.2 (VacSAU4), YM3.3 (mixture of VacSAU4 and 3M-052), YM3.6 (nanoparticle containing VacSAU4 and YAXA), and YM3.7 (nanoparticle containing VacSAU4, 3M-052, and YAXA). On 42 d post the first immunization, each mouse was anesthetized and subjected to aerosolized intratracheal challenge with a predetermined $LD_{100}$ dose (~$4\times10^8$ CFUs) of *S. aureus* USA300-FPR3757. Survival rates for all groups, as well as body weights, were monitored daily for 7 d following the challenge. At 6 h post *S. aureus* challenge, the blood, lungs, spleens, and livers were collected to quantify bacterial load by using plate counting method.

**Fluorescence imaging of whole-body of mice.** WT mice were subjected to single-dose immunization with the following 2 groups: YM3.3-Cy5.5 (mixture of VacPAE1, 3M-052, and Cy5.5), and YM3.7-Cy5.5 (nanoparticle containing VacPAE1, 3M-052, YAXA, and Cy5.5). Whole-body fluorescence imaging was measured longitudinally from 0 d to 8 d using IVIS Lumina III imaging system (PerkinElmer, USA) at 675 nm excitation/695 nm emission. Data are expressed as total integrated fluorescence intensity.

**Fluorescence imaging of secondary lymphoid organs of mice.** WT mice were subjected to single-dose immunization with the following 2 groups: YM3.3-Cy5.5 (mixture of VacPAE1, 3M-052, and Cy5.5), and YM3.7-Cy5.5 (nanoparticle containing VacPAE1, 3M-052, YAXA, and Cy5.5). The PMLN and spleen were excised at 24 h post single-dose immunization, and then fluorescence intensities were measured using IVIS Lumina III imaging system at 675 nm excitation/695 nm emission. Data are expressed as total integrated fluorescence intensity.

**Preparation of single-cell suspensions.** The lung, PMLN, or spleen was collected from each mouse and digested with 1 mg/mL of collagenase type IV for 20 min at 37 °C, followed by smashing with 100-μm strainers to make single-cell suspension. Red blood cells in the lung or spleen were lysed before further processing.

**Flow cytometry analysis of APC uptake in mice.** WT mice were subjected to single-dose immunization with the following 2 groups: YM3.3-Cy5.5 (mixture of VacPAE1, 3M-052, and Cy5.5), and YM3.7-Cy5.5 (nanoparticle containing VacPAE1, 3M-052, YAXA, and Cy5.5). At 24 h post single-dose immunization, single-cell suspensions were prepared from the lung, PMLN, and spleen. As described previously[15], these suspensions were stained with FVD780 and washed, and then cells were subsequently blocked with Fc-blocker prior to staining with the markers listed in Supplementary Table 1. Cy5.5 expression levels in cDC1, cDC2, pDC, and Mø were analyzed by LSRFortessa Flow Cytometer.

**Immunofluorescence staining of GC formation in mice.** WT mice were subjected to three-dose immunizations with the following 2 groups: YM3.3 (mixture of VacPAE1 and 3M-052), and YM3.7 (nanoparticle containing VacPAE1, 3M-052, and YAXA). On 42 d post the first immunization, the PMLN was isolated and snap frozen in molds containing OCT medium, and then dropped into 2-methyl butane cooled with liquid nitrogen. Frozen tissues were sectioned at 5 μm, fixed in ice-cold acetone for 10 min, air dried, and stored at -80°C. As described previously[16], sections were sequentially stained with DAPI, primary antibodies anti-mouse B220, and anti-mouse Ki67, followed by incubating by HRP conjugated Goat Anti-Rabbit IgG (H+L), and then TSA dyes were added: PE (red) for B220, FITC (green) for Ki67. Images were captured using Eclipse C1 Upright Fluorescence Microscope (Nikon, Japan) and Pannoramic MIDI Digital Slide Scanner (3DHISTECH, Hungary).

**Confocal laser fluorescence imaging of intracellular localization within BMDC.** BMDC was seeded at a density of $2.5 \times 10^5$ cells per mL of in confocal dishes and cultured for 10 min or 120 min with the following 2 groups: YM3.3-Cy5.5 (mixture of VacPAE1, 3M-052, and Cy5.5), and YM3.7-Cy5.5 (nanoparticle containing VacPAE1, 3M-052, YAXA, and Cy5.5). 5.56 μg/mL VacPAE1, 138.89 μg/mL YAXA, 5 μg/mL 3M-052, and 0.278 μg/mL Cy5.5 were used. Following treatment, cells were incubated with LysoTracker Green (100 nM) and Hoechst 33342 (2 μg/mL) in serum-free medium at 37°C for 30 min to label the lysosome and nucleus, respectively, as described previously[17]. Cells were washed 3 times with PBS and imaged using SP8 STED 3X Confocal Laser Scanning Microscope (Leica, Germany).

**Flow cytometry analysis of APC response in mice.** WT mice were subjected to single-dose immunization with the following 2 groups: YM3.3 (mixture of VacPAE1 and 3M-052), and YM3.7 (nanoparticle containing VacPAE1, 3M-052, and YAXA). Single-cell suspensions were prepared from the tissues of lung, PMLN, and spleen collected at the following 2 timepoints: 0 d (pre-immunization baseline), and 1 d post single-dose immunization[15]. Prior to staining, cells were incubated with anti-CD16/32 for 10 min on ice to block Fc receptors, thereby minimizing nonspecific antibody binding. As described previously[15], surface marker staining was performed using fluorochrome-conjugated antibodies (Supplementary Table 2) in FACS buffer (PBS supplemented with 3% FBS, 1 mM EDTA, and 0.02% sodium azide) for 30 min at 4°C under light-protected conditions. Following surface staining, cells were treated with FVD780 for 10 min. After twice washes with cold PBS, the maturation of cDC1, cDC2, pDC, and Mø was analyzed by LSRFortessa Flow Cytometer as above.

**Flow cytometry analysis of GC cell response in mice.** WT mice were subjected to three-dose immunizations with the following 2 groups: YM3.3 (mixture of VacPAE1 and 3M-052), and YM3.7 (nanoparticle containing VacPAE1, 3M-052, and YAXA). Single-cell suspensions were prepared from the PMLN and spleen collected on 0 d (pre-immunization baseline), and 42 d post the first immunization. Cells were blocked with anti-CD16/32, and then stained with the below specific markers for 30 min at 4°C under light-protected conditions. GCB and TFH cell markers[18] and corresponding antibodies were listed in Supplementary Table 3, while FDC markers[19] and corresponding antibodies in Supplementary Table 4. After surface staining, cells were incubated with FVD780 for 10 min at 4°C. Cells were then washed twice with ice-cold PBS and analyzed immediately by LSRFortessa Flow Cytometer as above.

**Flow cytometry analysis of B cell response in mice.** WT mice were subjected to three-dose immunizations with the following 2 groups: YM3.3 (mixture of VacPAE1 and 3M-052), and YM3.7 (nanoparticle containing VacPAE1, 3M-052, and YAXA). Single-cell suspensions were prepared from the lung, PMLN, and spleen collected on 0 d (pre-immunization baseline), and 42 d post the first immunization. Cells were blocked with anti-CD16/32, and then stained with corresponding antibodies for 30 min at 4°C in the dark. Antibody panels of EBC[18], MBC[20], and Brm[21] were listed in Supplementary Table 5. After surface staining, cells were incubated with FVD780 for 10 min at 4°C. Cells were then washed twice with ice-cold PBS, and immediately analyzed by LSRFortessa Flow Cytometer as above.

**Preparation of VacPAE1-specific peptide pool.** To generate a VacPAE1-specific peptide pool, overlapping 11-mer peptides (4-amino-acid overlap) were designed to achieve 1.6-fold sequence coverage. Peptide synthesis was performed by Genscript Biotech Corporation (China) via Fmoc-based solid-phase chemistry. Each synthesized peptide was purified (>70% purity) using reversed-phase HPLC, lyophilized, and stored at -80°C. For peptide pool preparation, individual peptides were reconstituted in DMSO, pooled at equimolar ratios, and diluted to working concentrations (1 μg/mL per peptide).

**Flow cytometry analysis of antigen-specific T cell response in mice.** WT mice were subjected to three-dose immunizations with the following 2 groups: YM3.3 (mixture of VacPAE1 and 3M-052), and YM3.7 (nanoparticle containing VacPAE1, 3M-052, and YAXA). Single-cell suspensions were prepared from lung, PMLN, and spleen collected on 0 d (pre-immunization baseline), and 42 d post the first immunization[18]. Cells were adjusted to $2\times10^6$ cells per tube and restimulated overnight at 37°C with VacPAE1-specific peptide pools (1 μg/mL per peptide) in RPMI 1640 medium with protein transport inhibitor containing brefeldin A (10 μg/mL). Following 18-h cultivation, cells were sequentially stained with FVD780 on ice for 10 min, washed, and blocked with anti-CD16/32 for 5 min. As described previously[18], surface staining was performed using antibodies (Supplementary

Table 6) for 30 min, and then cells were washed, fixed and permeabilized with Cytofix/Cytoperm buffer, and further stained with intracellular antibodies (Supplementary Table 6). Th1, Th2, Tc1, Tc2, $CD4^+$ Trm, and $CD8^+$ Trm cell subsets were identified by LSRFortessa Flow Cytometer as above.

**bRNA-seq and scRNA-seq experiments.** WT mice were subjected to single-dose immunization with YM3.7 (nanoparticle containing VacPAE1, 3M-052, and YAXA). As described previously[18], the following 4 sampling timepoints were set: 0 d prior to immunization, and 1/7/14 d post single-dose immunization. The lung of each mouse was collected and digested into single-cell suspension. Cells were incubated with biotinylated anti-CD45 antibody on ice for 20 min. Subsequently, streptavidin-labeled magnetic beads were added and incubated on ice for 30 min. As described previously[16], $CD45^+$ bound and $CD45^-$ unbound cells were collected and combined at a ratio of 9:1. Cells were resuspended in PBS containing 1% BSA and 0.5 U/μL RNase inhibitor and submitted to GENEWIZ (Azenta, China) for bulk RNA-seq and scRNA-seq.

**bRNA-seq data mining.** bRNA-seq data were processed using R (v4.4.1). DEGs were identified using edgeR (v3.42.4)[22]. DEGs were defined as genes with an adjusted $P < 0.05$ and $|\log2(\text{fold change})| > 1$ when compared to 0 d. Dynamic expression patterns of DEGs were clustered using Mfuzz package (v2.58.0) with fuzzy C-means (parameters: 3 clusters, fuzzy coefficient $m=2$)[23]. To further analyze the gene clusters, GO enrichment analysis was performed using clusterProfiler (v4.10.0)[24].

**scRNA-seq data mining.** Raw sequencing data were aligned to the GRCh38 reference genome (10×Genomics, v3.0.0) using Cell Ranger (v3.1.0). Raw count data were filtered to remove cells with a mitochondrial RNA fraction of >5% of total RNA counts per cell, cells with <100 unique features or cells with < 200 total reads. The filtered count matrix from 4 timepoints (0 d pre-immunization, and 1/7/14 d post single-dose immunization) was used to create Seurat S4 object (v4.3.0) and was scaled by a factor of 10000 and log-transformed. The remaining 46699 cells were processed with the default Seurat pipeline. Specifically, the most variable 5000 RNA features were used to perform PCA on the log-transformed counts. The first 50 components were used for further downstream analyses, including clustering and UMAP projections. Clusters were identified with Seurat SNN graph construction followed by Louvain community detection on the resultant graph with a resolution of 2, yielding 64 clusters, which were subsequently consolidated into 13 biologically meaningful cell populations[25]. Absolute APC/B/T cell counts were computed by scaling each subset's proportion among immune cells (cDC1, cDC2, pDC, AM, IM, Mono, B, T, NK, and Neu) to the total measured $CD45^+$ cell count. Differential expression for each timepoint compared to 0 d was calculated with Wilcoxon rank-sum test. GO enrichment analysis was further used to analyze the differential expressed genes[24]. All analysis was performed in R (v4.4.1).

**Immune response of BMDC treated in vitro.** BMDC samples isolated from WT and *Tlr7*$^{-/-}$ mice were incubated with the following 5 treatment groups: Mock (PBS), YM3.1 (3M-052), YM3.4 (nanoparticles containing YAXA), YM3.5 (nanoparticles containing 3M-052 and YAXA), and YM3.7 (nanoparticles containing 3M-052, YAXA, and VacPAE1). The concentrations used were 5.56 μg/mL VacPAE1, 138.89 μg/mL YAXA, or 5 μg/mL 3M-052. First, cell maturation was assessed by flow cytometry. As described previously[26], cells were incubated for 6 h and then stained with FVD780 as well as antibodies (Supplementary Table 7). Identification of cell markers was then performed by LSRFortessa Flow Cytometer as above. Second, protein levels in whole cell lysates were assessed by Western blotting. At 3 h post incubation, cells were lysed by cell lysis buffer with 1× protease/phosphatase inhibitor followed by immunoblotting with the following primary antibodies: TLR7 (1:1000), IκBα (1:1000), p-IκBα (1:1000), p65 (1:1000), p-p65 (1:1000), and anti-GAPDH (1:5000). Then, the mixtures were incubated with appropriate secondary antibodies. Blots were developed with enhanced chemiluminescence kit. Third, cytokine levels in cell supernatants were measured by ELISA. Cells were stimulated for 6 h and then culture supernatants were collected to measure the concentrations of IL-12p70 and TNF-α by ELISA.

**IgG production and T cell response in YM3.7-immunized WT and *Tlr7*$^{-/-}$ mice.** WT and *Tlr7*$^{-/-}$ mice were subjected to three-dose immunizations with YM3.7 (nanoparticle containing VacPAE1, 3M-052, and YAXA). Serum was collected on 42 d post the first immunization, and anti-VacPAE1 IgG titers were determined by ELISA. Single-cell suspensions were prepared from the lung, PMLN, and spleen collected on 42 d post the first immunization. Th1, Th2, Tc1, and Tc2 cell subsets in the lung, PMLN, and spleen, along with CD4$^+$ Trm and CD8$^+$ Trm cell subsets in the lung, were analyzed by LSRFortessa Flow Cytometer as above.

**Statistical analysis.** All statistical analyses were performed using GraphPad Prism (v8.0), with experimental data presented as mean ± standard deviation (SD). Normality of data distribution was first assessed using Shapiro-Wilk test, followed by parametric or non-parametric testing: for comparisons between 2 groups with normal distribution, unpaired two-tailed Student's *t*-test was applied, while non-normally distributed data were analyzed using Mann-Whitney U test; comparisons among 3 or more groups under normal distribution were subjected to one-way ANOVA with Dunn's honestly significant difference post hoc test for pairwise comparisons, whereas non-parametric Kruskal-Wallis test with Dunn's multiple comparisons correction was used for skewed distributions. For experimental designs involving 2 independent variables (e.g., treatment groups and timepoints), two-way ANOVA was performed, followed by Dunn's or Sidak's multiple comparisons test to evaluate main effects, interactions, and pairwise differences. Mouse survival outcomes were analyzed using Kaplan-Meier survival curves, with between-group differences assessed via log-rank (Mantel-Cox) test. Correlation analyses between continuous variables were conducted using Spearman's rank correlation coefficient. All statistical tests were two-sided, and significance thresholds were defined as follows: $P < 0.05$ (*), $P < 0.01$ (**), and $P < 0.001$ (***).

**Animal ethical clearance.** Female WT C57BL/6J mice (6-8 weeks old) were purchased from Vital River (Beijing, China). Female *Tlr7*$^{-/-}$ mice (6-8 weeks old) in C57BL/6J background were purchased from Cyagen (Suzhou, China). The use of female mice was based on their relative docility and lower procedural variability, rather than a sex-specific research focus. All the animals were cared under pathogen-free conditions, 12-h light/12-h dark cycle, temperatures of 18-23°C with 40-60% humidity. The study protocol was reviewed and approved by Institute of Animal Care and Use Committee.

**Supplementary Table 1** Antibodies used in flow cytometry analysis of APC uptake

| Marker | Dilution | Dye | Manufacturer | Catalogue no. |
|---|---|---|---|---|
| CD45 | 1:100 | BUV395 | BD Biosciences | 564279 |
| CD3e | 1:100 | BV605 | BD Biosciences | 563004 |
| CD11b | 1:125 | FITC | BD Biosciences | 557396 |
| F4/80 | 1:125 | AF647 | BioLegend | 123122 |
| MHC-II | 1:125 | PE | BioLegend | 107608 |
| CD11c | 1:100 | BV785 | BioLegend | 117336 |
| CD103 | 1:100 | BV510 | BioLegend | 141223 |
| pDCA-1 | 1:100 | BV421 | BioLegend | 127023 |

**Supplementary Table 2** Antibodies used in flow cytometry analysis of APC response

| Marker | Dilution | Dye | Manufacturer | Catalogue no. |
|---|---|---|---|---|
| CD45 | 1:100 | PerCP-Cy5.5 | BD Biosciences | 550994 |
| CD3e | 1:100 | BV605 | BD Biosciences | 563004 |
| CD11b | 1:125 | FITC | BD Biosciences | 557396 |
| F4/80 | 1:125 | AF647 | BioLegend | 123122 |
| MHCII | 1:125 | AF700 | BioLegend | 107622 |
| CD11c | 1:100 | BV785 | BioLegend | 117336 |
| CD86 | 1:100 | PE-Cy7 | BioLegend | 105014 |
| CD103 | 1:100 | BV510 | BioLegend | 141223 |
| pDCA-1 | 1:100 | BV421 | BioLegend | 127023 |

**Supplementary Table 3** Antibodies used in flow cytometry analysis of GCB and TFH cells

| Marker | Dilution | Dye | Manufacturer | Catalogue no. |
|---|---|---|---|---|
| CD45 | 1:100 | BUV395 | BD Biosciences | 564279 |
| CD3e | 1:100 | BV421 | BD Biosciences | 100341 |
| CD19 | 1:100 | APC | BD Biosciences | 550992 |
| CD4 | 1:100 | BV510 | BD Biosciences | 563106 |
| CXCR5 | 1:100 | PE-Cy7 | BD Biosciences | 560617 |
| PD1 | 1:100 | BV605 | BD Biosciences | 563059 |
| CD95 | 1:125 | FITC | BD Biosciences | 554257 |
| CD38 | 1:100 | PE | BD Biosciences | 553764 |

**Supplementary Table 4** Antibodies used in flow cytometry analysis of FDC

| Marker | Dilution | Dye | Manufacturer | Catalogue no. |
|---|---|---|---|---|
| CD45 | 1:100 | BUV395 | BD Biosciences | 564279 |
| PDPN | 1:100 | PE | Thermo Fisher Scientific | 2866239 |
| CD31 | 1:100 | PE-Cy7 | BD Biosciences | 561410 |
| CD21/35 | 1:100 | BV421 | BioLegend | 123422 |

**Supplementary Table 5** Antibodies used in flow cytometry analysis of B cell response

| Marker | Dilution | Dye | Manufacturer | Catalogue no. |
|---|---|---|---|---|
| CD45 | 1:100 | BUV395 | BD Biosciences | 564279 |
| CD19 | 1:100 | AF488 | BioLegend | 115521 |
| CD3e | 1:100 | BV421 | BD Biosciences | 100341 |
| CD44 | 1:100 | BV605 | BioLegend | 103040 |
| CD138 | 1:100 | PE-Cy7 | BioLegend | 142514 |
| PDL2 | 1:100 | PE | BioLegend | 107205 |
| CD80 | 1:125 | AF647 | BioLegend | 104718 |
| CD69 | 1:100 | PerCP-Cy5.5 | BD Biosciences | 551113 |

**Supplementary Table 6** Flow cytometry surface staining of T cell response

| Marker | Dilution | Dye | Manufacturer | Catalogue no. | Surface or intracelluar staining |
|---|---|---|---|---|---|
| CD3e | 1:100 | BV650 | BD Biosciences | 564378 | Surface |
| CD8a | 1:100 | BV510 | BioLegend | 100751 | Surface |
| CD4 | 1:125 | FITC | BD Biosciences | 553046 | Surface |
| CD44 | 1:100 | BV650 | BD Biosciences | 740455 | Surface |
| CD69 | 1:100 | PE-Cy7 | BD Biosciences | 552879 | Surface |
| CD103 | 1:100 | BV421 | BD Biosciences | 562771 | Surface |
| IFN-γ | 1:100 | PerCP-Cy5.5 | BD Biosciences | 560660 | Intracellular |
| IL-13 | 1:100 | PE | BD Biosciences | 568551 | Intracellular |

**Supplementary Table 7** Antibodies used in flow cytometry analysis of BMDC maturation

| Marker | Dilution | Dye | Manufacturer | Catalogue no. |
|---|---|---|---|---|
| CD40 | 1:100 | PE | BioLegend | 124610 |
| CD80 | 1:100 | APC | BioLegend | 104714 |
| CD86 | 1:100 | PE-Cy7 | BioLegend | 105014 |
| CD11c | 1:125 | FITC | BioLegend | 117306 |
| MHC-II | 1:100 | PerCP-Cy5.5 | BD Biosciences | 562363 |

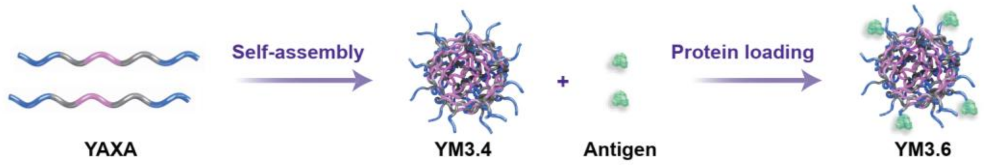

**Supplementary Fig. 1** Schematic illustration of preparation of YM3.4/6.

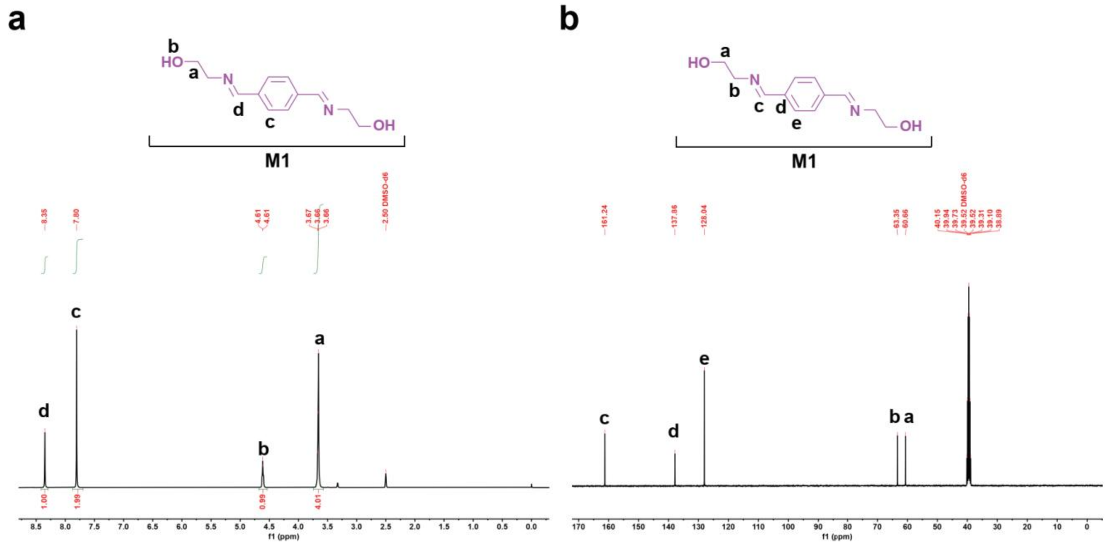

**Supplementary Fig. 2 a)** $^{1}$H NMR and **b)** $^{13}$C NMR spectra of monomer M1 (10 mg/mL). Experiments are repeated for 2 individual batches.

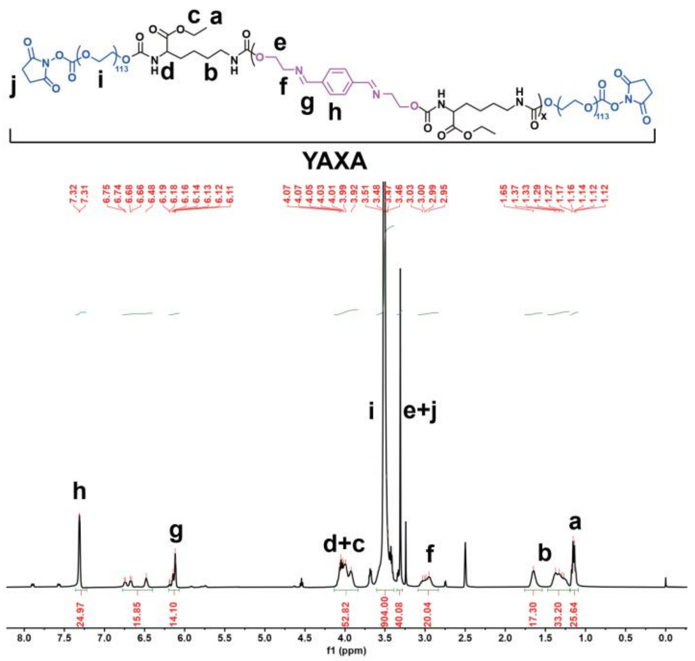

**Supplementary Fig. 3** $^{1}$H NMR spectrum of YAXA (10 mg/mL). Experiments are repeated for 2 individual batches.

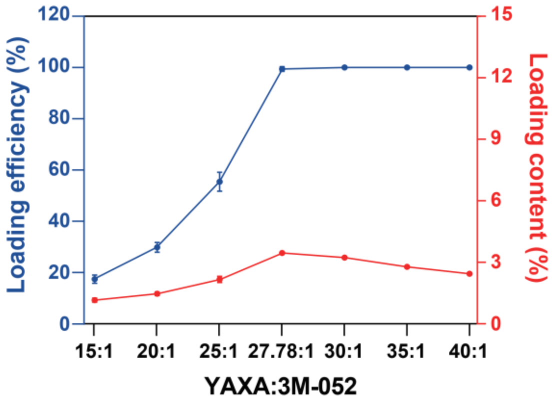

**Supplementary Fig. 4** Assessment of 3M-052 loading during preparation of YM3.5 (nanoparticle containing YAXA and 3M-052). 3M-052 is used at a fixed concentration of 360 µg/mL, while YAXA are systematically varied from 15:1 to 40:1. This is a pre-experiment to reveal the result that YAXA:3M-052 ratios ≥27.78:1 achieve nearly 100% 3M-052 loading. n=5 biologically independent replicates. Data are presented as mean ± SD.

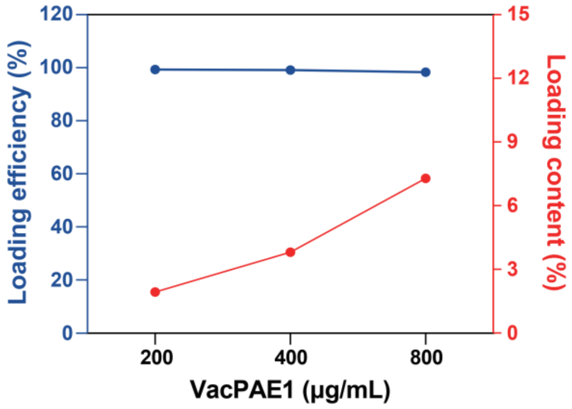

**Supplementary Fig. 5** Assessment of VacPAE1 loading during preparation of YM3.7 (nanoparticle containing YAXA, 3M-052, and VacPAE1). YAXA:3M-052 mass ratio is fixed at 27.78:1. 10 mg/mL of YAXA and 360 µg/mL of 3M-052 are kept constant, while VacPAE1 is tested at 200, 400, and 800 µg/mL. This is a pre-experiment to validate the fact that NHS-activated esters on YM3.7 are redundant for VacPAE1 loading. n=5 biologically independent replicates. Data are presented as mean ± SD.

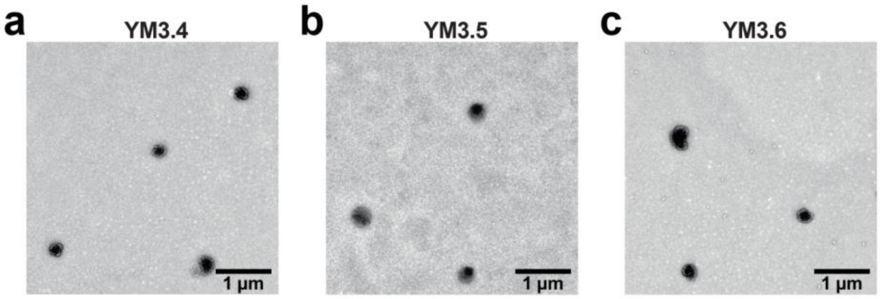

**Supplementary Fig. 6** TEM image of YM3.4/5/6. Shown are **a)** YM3.4 (nanoparticle containing pure YAXA), **b)** YM3.5 (nanoparticle containing YAXA and 3M-052), and **c)** YM3.6 (nanoparticle containing YAXA and VacPAE1). 4 µg/mL VacPAE1, 100 µg/mL YAXA, and 3.6 µg/mL 3M-052 are used. Experiments are repeated for 2 individual batches.

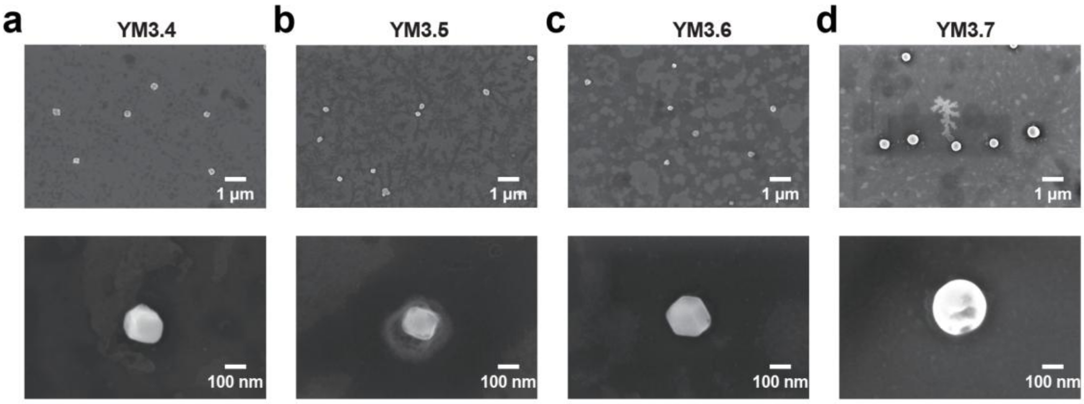

**Supplementary Fig. 7** SEM image of YM3.4/5/6. Shown are **a)** YM3.4 (nanoparticle containing pure YAXA), **b)** YM3.5 (nanoparticle containing YAXA and 3M-052), **c)** YM3.6 (nanoparticle containing YAXA and VacPAE1), and **d)** YM3.7 (nanoparticle containing YAXA, 3M-052 and VacPAE1). 4 µg/mL VacPAE1, 100 µg/mL YAXA, and 3.6 µg/mL 3M-052 are used. Experiments are repeated for 2 individual batches.

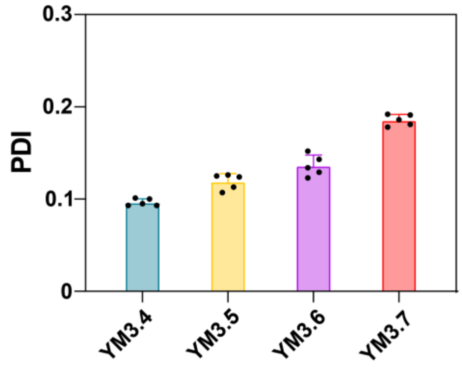

**Supplementary Fig. 8** PDI value of YM3.4/5/6/7 in PBS. YM3.4: nanoparticle containing pure YAXA. YM3.5: nanoparticle containing YAXA and 3M-052. YM3.6: nanoparticle containing YAXA and VacPAE1. YM3.7: nanoparticle containing YAXA, 3M-052, and VacPAE1. n=5 biologically independent replicates. Data are presented as mean ± SD.

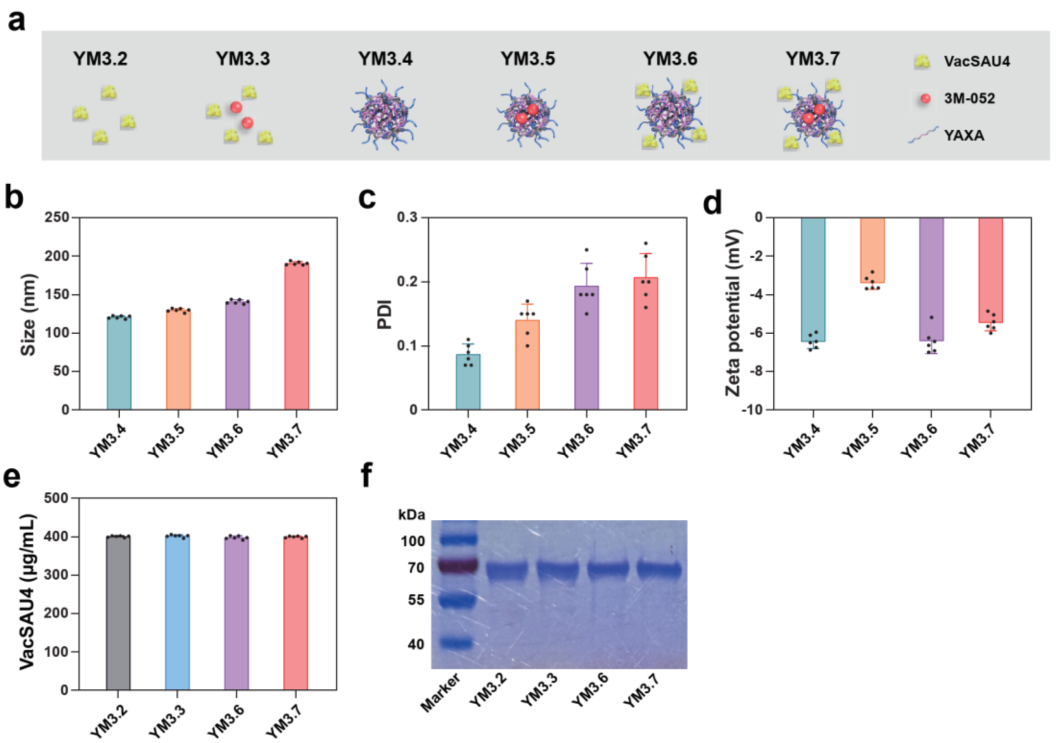

**Supplementary Fig. 9** Characterization of nanoparticle YM3.7 loading VacSAU4. **a)** Schematic illustration of the 6 experimental groups: YM3.2 (VacSAU4), YM3.3 (mixture of VacSAU4 and 3M-052), YM3.4 (nanoparticle containing pure YAXA), YM3.5 (nanoparticle containing YAXA and 3M-052), YM3.6 (nanoparticle containing YAXA and VacSAU4), and YM3.7 (nanoparticle containing YAXA, 3M-052, and VacSAU4). DLS measurement of **b)** hydrodynamic size, **c)** PDI, and **d)** zeta potential of YM3.4/5/6/7 (n=6 biologically independent replicates). **e)** BCA assay of VacSAU4 content in YM3.2/3/6/7 (n=6 biologically independent replicates). **f)** SDS-PAGE analysis of VacSAU4 in YM3.2/3/6/7. Experiments are repeated for 2 individual batches. Data are presented as mean ± SD.

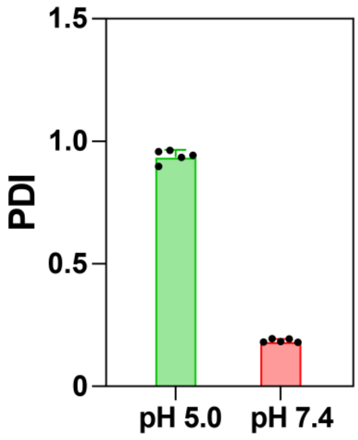

**Supplementary Fig. 10** PDI values of YM3.7 (nanoparticle containing YAXA, 3M-052, and VacPAE1) at pH 5.0/7.4 over 72 h. n=5 biologically independent replicates. Data are presented as mean ± SD.

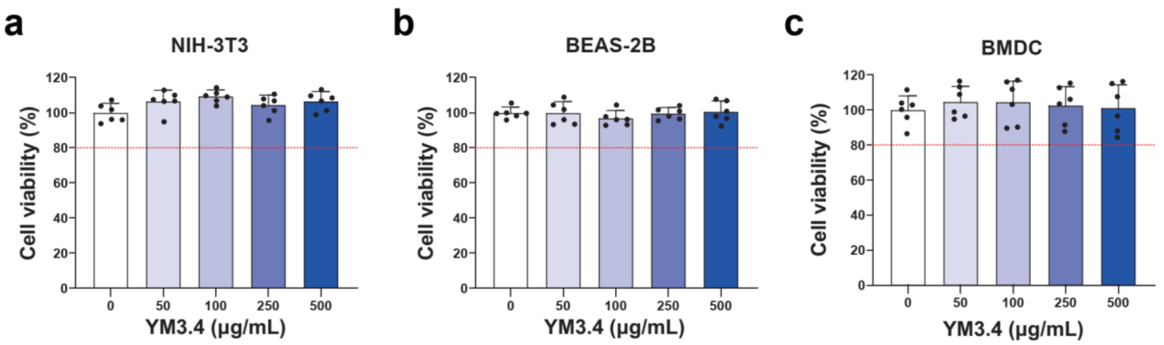

**Supplementary Fig. 11** Cytotoxicity assessment of YM3.4 (nanoparticle containing pure YAXA). Shown are viability of **a)** NIH-3T3, **b)** IOSE-80, and **c)** BMDC cells incubated with YM3.4 at varying concentrations for 24 h. n=6 biologically independent replicates. Data are presented as mean ± SD.

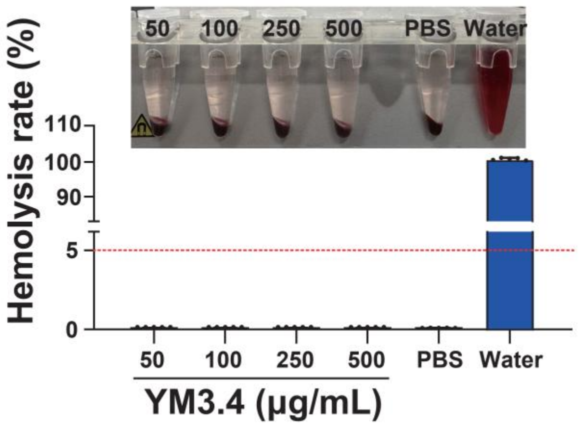

**Supplementary Fig. 12** Hemolysis rate of YM3.4 (nanoparticle containing pure YAXA). Red blood cells are incubated with varying concentrations of YM3.4. PBS and water are used as the negative and positive controls, respectively. n=5 biologically independent replicates. Data are presented as mean ± SD.

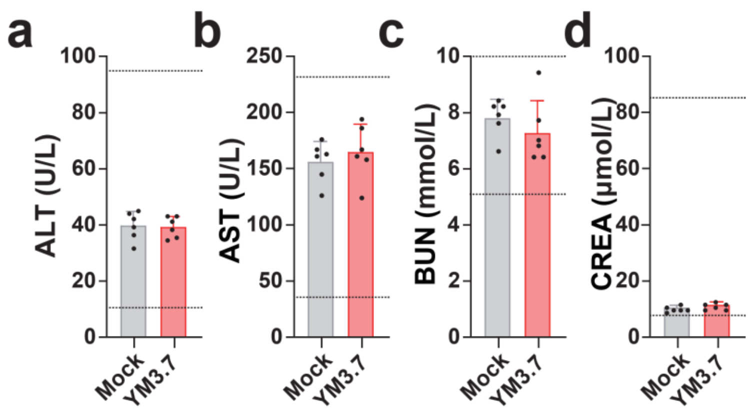

**Supplementary Fig. 13** Detection of biochemical blood indicators induced by YM3.7. Mice are immunized with single-dose YM3.7 (nanoparticle containing YAXA, 3M-052, and VacPAE1) or Mock (PBS). Each inoculation contained 20 μg of VacPAE1, 18 μg of 3M-052, and 500 μg of YAXA. Sera from the Mock and YM3.7 groups are analyzed by biochemical blood tests on 7 d post single-dose immunization. Shown are **a)** alanine aminotransferase (ALT), **b)** aspartate aminotransferase (AST), **c)** blood urea nitrogen (BUN), and **d)** creatinine (CREA). n=6 biologically independent replicates. Data are presented as mean ± SD.

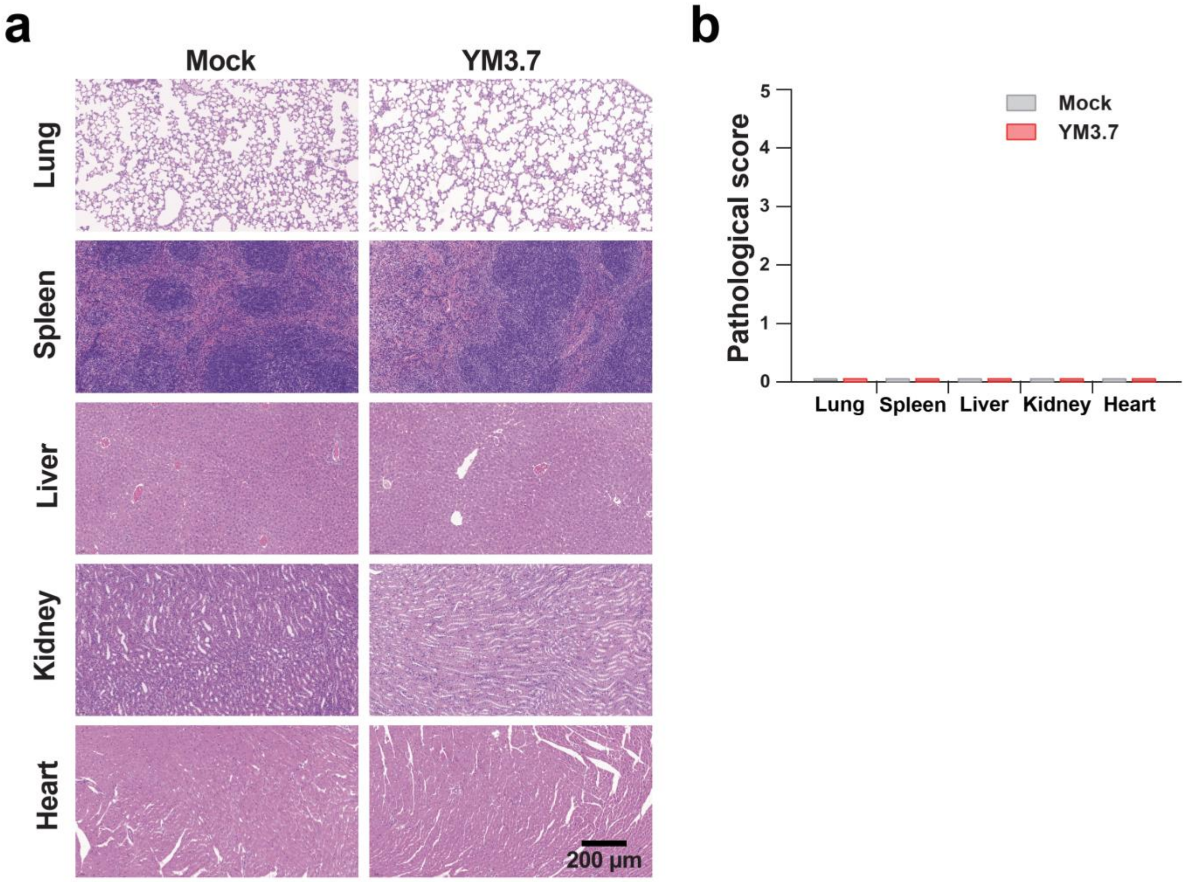

**Supplementary Fig. 14** H&E staining of major organs on 7 d post single-dose immunization of YM3.7. Mice are immunized with single-dose YM3.7 (nanoparticle containing YAXA, 3M-052, and VacPAE1) or Mock (PBS). Each inoculation contained 20 µg of VacPAE1, 18 µg of 3M-052, and 500 µg of YAXA. Major organs from the Mock and YM3.7 groups are analyzed by H&E staining on 7 d post single-dose immunization. Shown are **a)** representative H&E staining images and **b)** pathological scores. n=5 biologically independent replicates. Data are presented as mean ± SD.

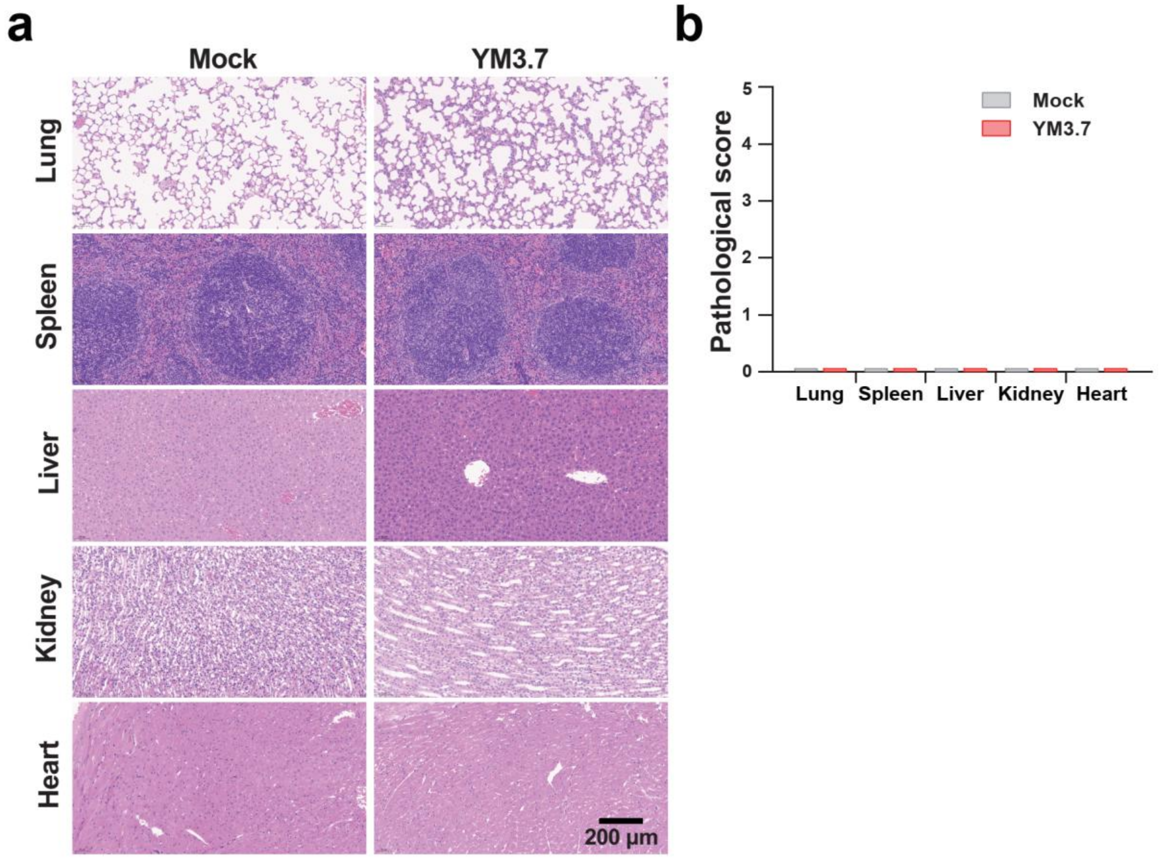

**Supplementary Fig. 15** H&E staining of major organs on 28 d post single-dose immunization of YM3.7. Mice are immunized with single-dose YM3.7 (nanoparticle containing YAXA, 3M-052, and VacPAE1) or Mock (PBS). Each inoculation contained 20 µg of VacPAE1, 18 µg of 3M-052, and 500 µg of YAXA. Major organs from the Mock and YM3.7 groups are analyzed by H&E staining on 28 d post single-dose immunization. Shown are **a)** representative H&E staining images and **b)** pathological scores. n=5 biologically independent replicates. Data are presented as mean ± SD.

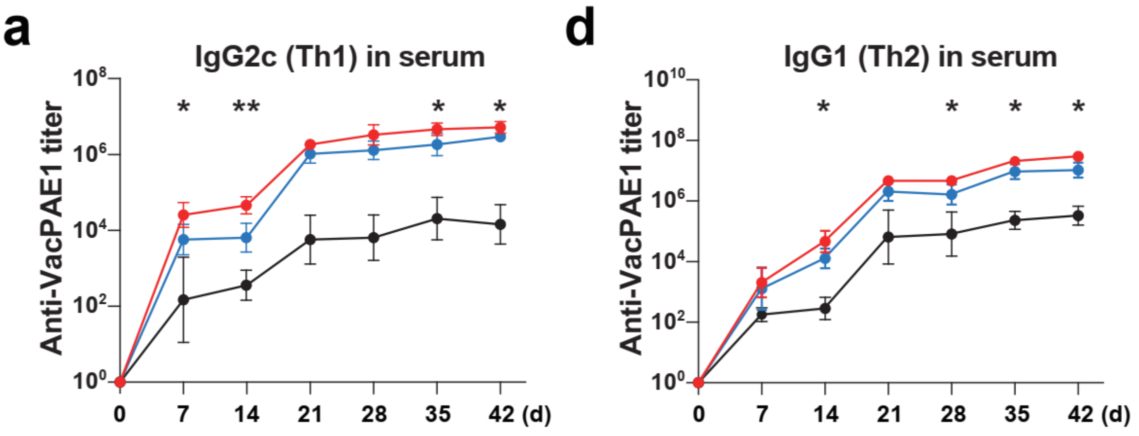

**Supplementary Fig. 16** ELISA measurement of VacPAE1-specific IgG subtypes in sera. Mice are immunized 3 times at 14-d intervals via aerosolized intratracheal inoculation with the following 3 experimental groups: YM3.2 (VacPAE1), YM3.3 (mixture of VacPAE1 and 3M-052), and YM3.7 (nanoparticle containing VacPAE1, 3M-052, and YAXA). 20 μg of VacPAE1, 18 μg of 3M-052, or 500 μg of YAXA are used per inoculation. Shown are **a)** IgG2c and **b)** IgG1. n=6 biologically independent replicates. Data are presented as mean ± SD. **Statistical significance test:** YM3.7 is compared with each of the other groups using two-way ANOVA with Dunn's multiple comparison test. *: *P*<0.05, **: *P*<0.01.

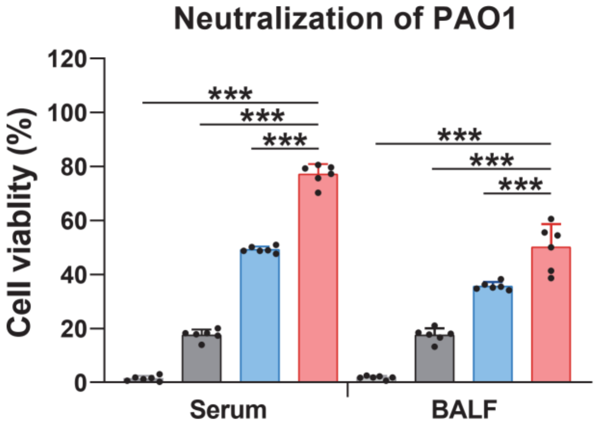

**Supplementary Fig. 17** Neutralization of *P. aeruginosa* PAO1 in sera and BALF assessed by CCK-8 cell viability assay. Mice are immunized 3 times at 14-d intervals via aerosolized intratracheal inoculation with the following 3 experimental groups: YM3.2 (VacPAE1), YM3.3 (mixture of VacPAE1 and 3M-052), and YM3.7 (nanoparticle containing VacPAE1, 3M-052, and YAXA). 20 μg of VacPAE1, 18 μg of 3M-052, or 500 μg of YAXA are used per inoculation. Serum and BALF samples are collected on 42 d post the first immunization. n=6 biologically independent replicates. Data are presented as mean ± SD. **Statistical significance test:** YM3.7 is compared with each of the other groups using one-way ANOVA with Dunn's multiple comparison test. ***: *P*<0.001.

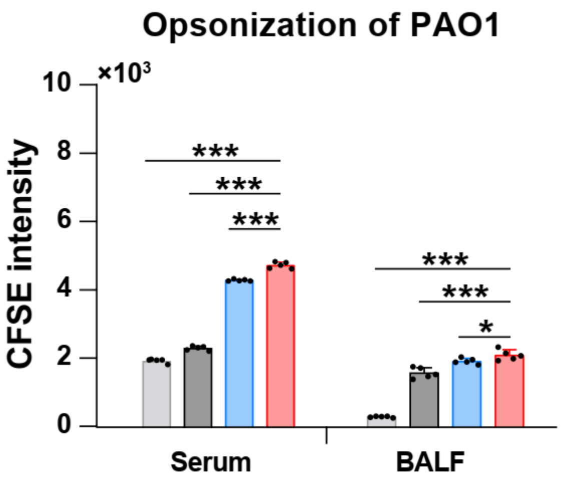

**Supplementary Fig. 18** Opsonophagocytic activity of sera and BALF measured using CFSE-labeled *P. aeruginosa* PAO1. Mice are immunized 3 times at 14-d intervals via aerosolized intratracheal inoculation with the following 3 experimental groups: YM3.2 (VacPAE1), YM3.3 (mixture of VacPAE1 and 3M-052), and YM3.7 (nanoparticle containing VacPAE1, 3M-052, and YAXA). 20 µg of VacPAE1, 18 µg of 3M-052, or 500 µg of YAXA are used per inoculation. Serum and BALF samples are collected on 42 d post the first immunization. n=5 biologically independent replicates. Data are presented as mean ± SD. **Statistical significance test:** YM3.7 is compared with each of the other groups using one-way ANOVA with Dunn's multiple comparison test. *: *P*<0.05, ***: *P*<0.001.

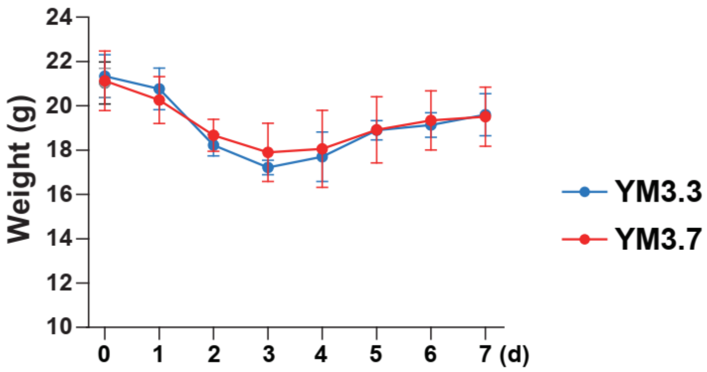

**Supplementary Fig. 19** Body weight changes of mice post *P. aeruginosa* challenge. Mice are immunized 3 times at 14-d intervals via aerosolized intratracheal inoculation with the following 4 experimental groups: Mock (PBS), YM3.2 (VacPAE1), YM3.3 (mixture of VacPAE1 and 3M-052), and YM3.7 (nanoparticle containing VacPAE1, 3M-052, and YAXA). 20 µg of VacPAE1, 18 µg of 3M-052, or 500 µg of YAXA are used per inoculation. The $LD_{100}$ value pre-determined at $1.4\times10^6$ CFUs of *P. aeruginosa* F291007 is used for aerosolized intratracheal challenge following three-dose immunizations. n=15 biologically independent replicates. Data are presented as mean ± SD. **Statistical significance test:** YM3.7 is compared with each of the other groups using two-way ANOVA with Dunn's multiple comparison test.

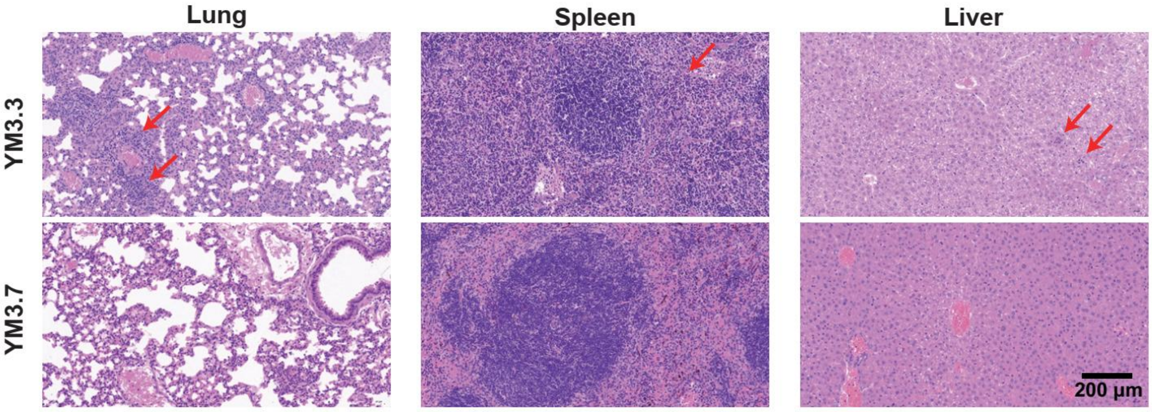

**Supplementary Fig. 20** Representative H&E staining images for the lung, spleen, and liver on 2 d post *P. aeruginosa* challenge. Mice are immunized 3 times at 14-d intervals via aerosolized intratracheal inoculation with the following 2 experimental groups: YM3.3 (mixture of VacPAE1 and 3M-052), and YM3.7 (nanoparticle containing VacPAE1, 3M-052, and YAXA). 20 μg of VacPAE1, 18 μg of 3M-052, or 500 μg of YAXA are used per inoculation. The $LD_{100}$ value pre-determined at $1.4\times10^6$ CFUs of *P. aeruginosa* F291007 is used for aerosolized intratracheal challenge following three-dose immunizations. Red arrows indicate inflammatory infiltration. n=6 biologically independent replicates.

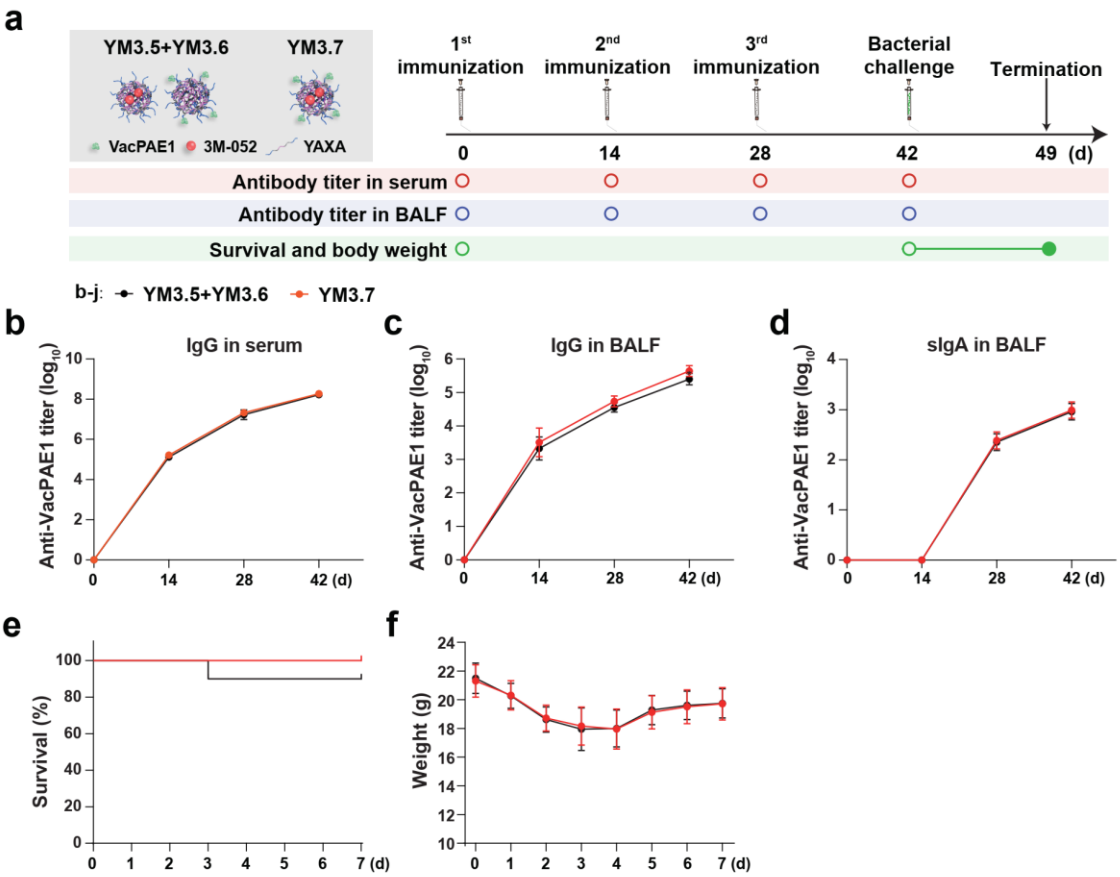

**Supplementary Fig. 21** Comparative evaluation of YM3.7 and YM3.5+YM3.6. **a)** Schematic diagram of experimental design. Mice are immunized 3 times at 14-d intervals via aerosolized intratracheal inoculation with the following 2 experimental groups: YM3.7 (nanoparticle containing VacPAE1, 3M-052, and YAXA), and YM3.5+YM3.6 [physical mixture of YM3.5 (nanoparticle containing 3M-052 and YAXA) and YM3.6 (nanoparticle containing VacPAE1 and YAXA)]. 20 µg of VacPAE1, 18 µg of 3M-052, or 500 µg of YAXA are used per inoculation. The $LD_{100}$ value pre-determined at $1.4\times10^{6}$ CFUs of *P. aeruginosa* F291007 is used for aerosolized intratracheal challenge following three-dose immunizations. Hollow circles denote timepoints pre-immunization or pre-challenge. **b)** ELISA measurement of VacPAE1-specific IgG in sera (n=6 biologically independent replicates). ELISA measurement of VacPAE1-specific **c)** IgG and **d)** sIgA in BALF (n=5 biologically independent replicates). **e)** Survival curves and **f)** body weight changes of mice post *P. aeruginosa* challenge (n=10 biologically independent replicates). Data are presented as mean ± SD. **Statistical significance test: b-d, f)** Two-way ANOVA with Sidak's multiple comparison tests is used to compare YM3.7 and YM3.5+YM3.6; **e)** survival analysis is conducted using Log-rank (Mantel-Cox) test.

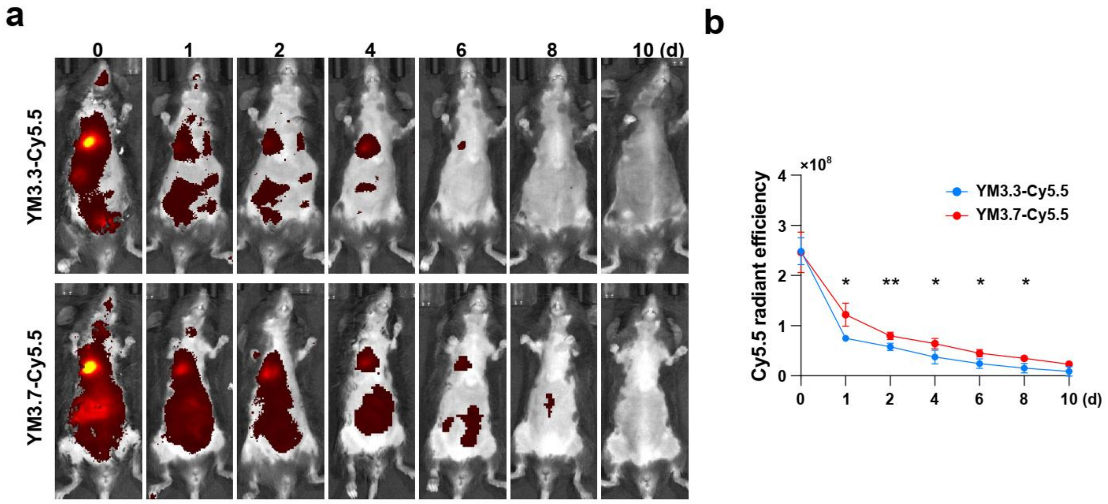

**Supplementary Fig. 22** Real-time IVFI assessment of YM3.3/7-Cy5.5 in mice. Mice are immunized with single-dose YM3.3-Cy5.5 (mixture of VacPAE1, 3M-052, and Cy5.5), or YM3.7-Cy5.5 (nanoparticle containing VacPAE1, 3M-052, YAXA, and Cy5.5). 20 μg of VacPAE1, 18 μg of 3M-052, 500 μg of YAXA, or 1 μg of Cy5.5 are used per inoculation. Shown are **a)** representative images and **b)** radiant efficiency. n=5 biologically independent replicates. Data are presented as mean ± SD. **Statistical significance test: b)** Two-way ANOVA followed by Dunn's post hoc test (YM3.7-Cy5.5 v.s. YM3.3-Cy5.5). *: $P < 0.05$, **: $P < 0.01$.

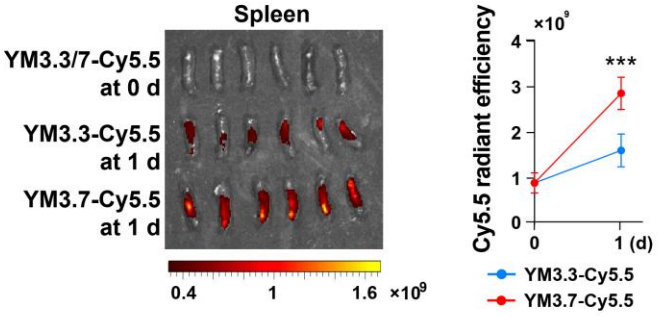

**Supplementary Fig. 23** IVFI assessment of accumulation of YM3.3/7-Cy5.5 in the spleen. Mice are immunized with single-dose YM3.3-Cy5.5 (mixture of VacPAE1, 3M-052, and Cy5.5), or YM3.7-Cy5.5 (nanoparticle containing VacPAE1, 3M-052, YAXA, and Cy5.5). 20 μg of VacPAE1, 18 μg of 3M-052, 500 μg of YAXA, or 1 μg of Cy5.5 are used per inoculation. n=6 biologically independent replicates. Data are presented as mean ± SD. **Statistical significance test:** Two-way ANOVA followed by Dunn's post hoc test (YM3.7-Cy5.5 v.s. YM3.3-Cy5.5). *: $P < 0.05$.

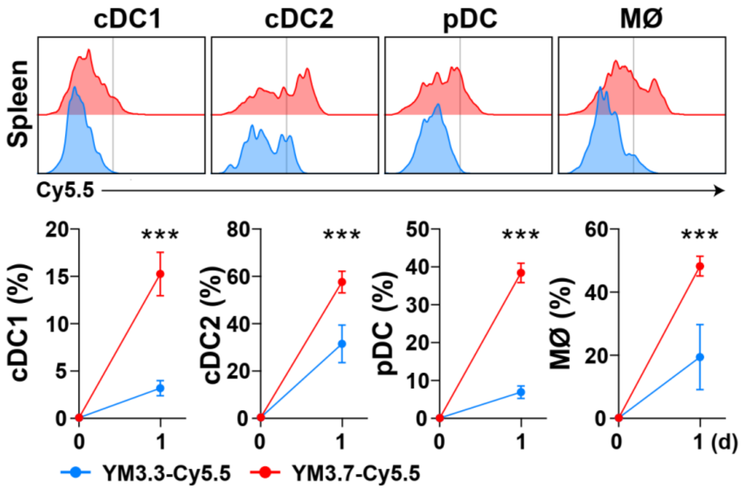

**Supplementary Fig. 24** Flow cytometry analysis of uptake of YM3.3/7-Cy5.5 by APC in the spleen. Mice are immunized with single-dose YM3.3-Cy5.5 (mixture of VacPAE1, 3M-052, and Cy5.5), or YM3.7-Cy5.5 (nanoparticle containing VacPAE1, 3M-052, YAXA, and Cy5.5). 20 μg of VacPAE1, 18 μg of 3M-052, 500 μg of YAXA, or 1 μg of Cy5.5 are used per inoculation. n=5 biologically independent replicates. Data are presented as mean ± SD. **Statistical significance test:** Two-way ANOVA followed by Dunn's multiple comparison test (YM3.7-Cy5.5 v.s. YM3.3-Cy5.5). ***: $P < 0.001$.

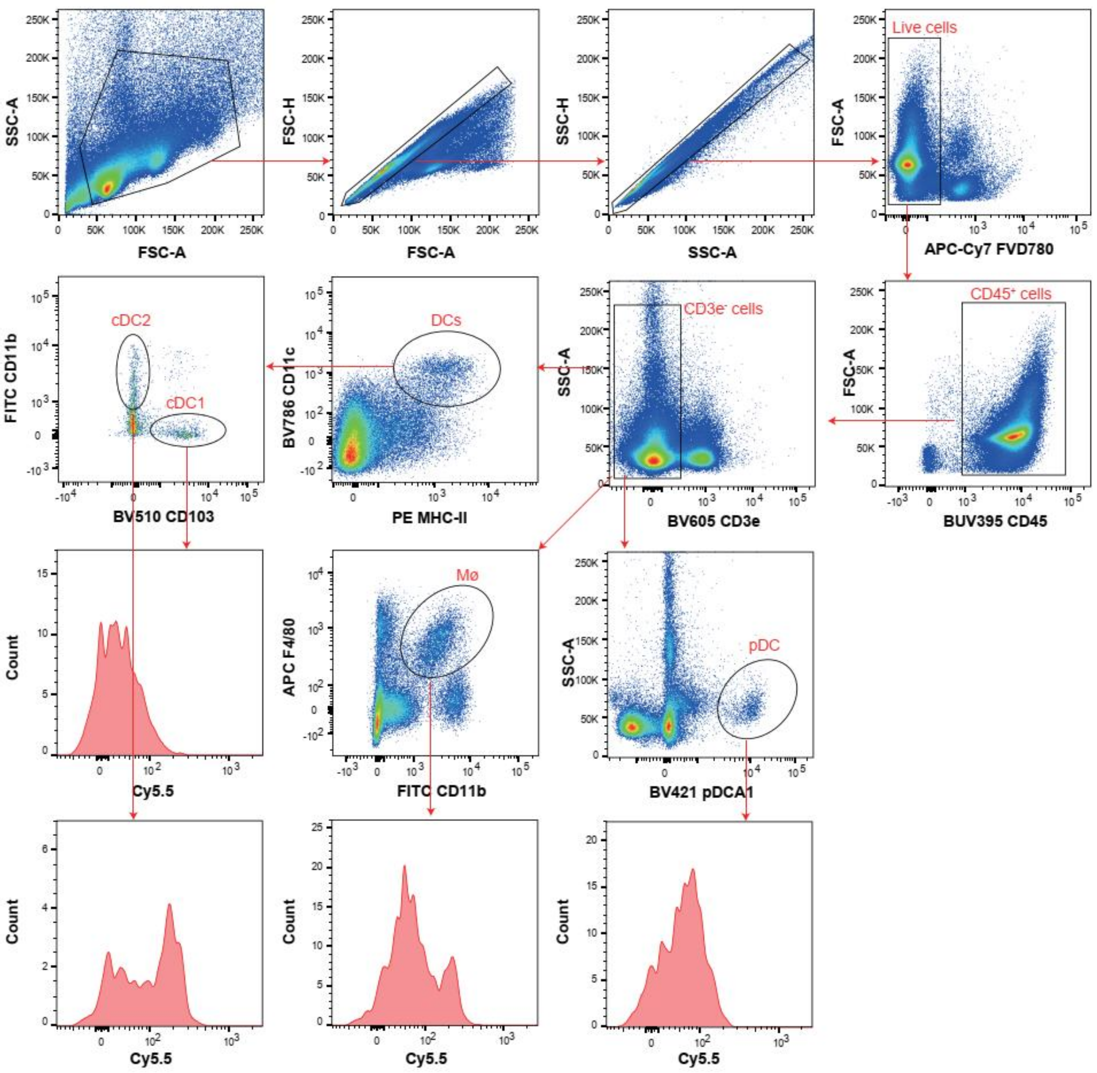

**Supplementary Fig. 25** Gating Strategy for flow cytometry analysis of APC with uptake of YM3.3/7-Cy5.5.

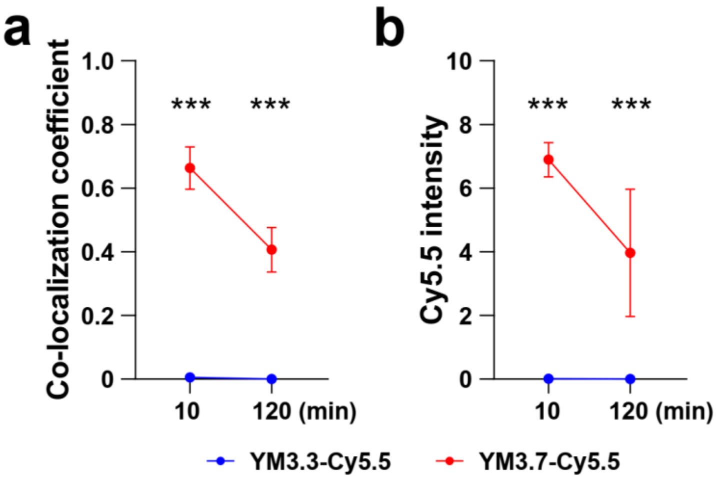

**Supplementary Fig. 26** Confocal laser fluorescence imaging of intracellular localization of YM3.3/7-Cy5.5 within BMDC. BMDC is incubated with YM3.3-Cy5.5 (mixture of VacPAE1, 3M-052, and Cy5.5) or YM3.7-Cy5.5 (nanoparticle containing VacPAE1, 3M-052, YAXA, and Cy5.5) for 10 min and 120 min. 5.56 μg/mL VacPAE1, 138.89 μg/mL YAXA, 5 μg/mL 3M-052, and Cy5.5 0.278 μg/mL are used. Shown are **a)** co-localization coefficient, and **b)** Cy5.5 intensity. n=5 biologically independent replicates. Data are presented as mean ± SD. **Statistical significance test: a)** Colocalization coefficient between Cy5.5 and LysoTraker is determined by Pearson correlation coefficient. **a-b)** Two-way ANOVA followed by Dunn's multiple comparison test (YM3.7-Cy5.5 v.s. YM3.3-Cy5.5). ***: $P < 0.001$.

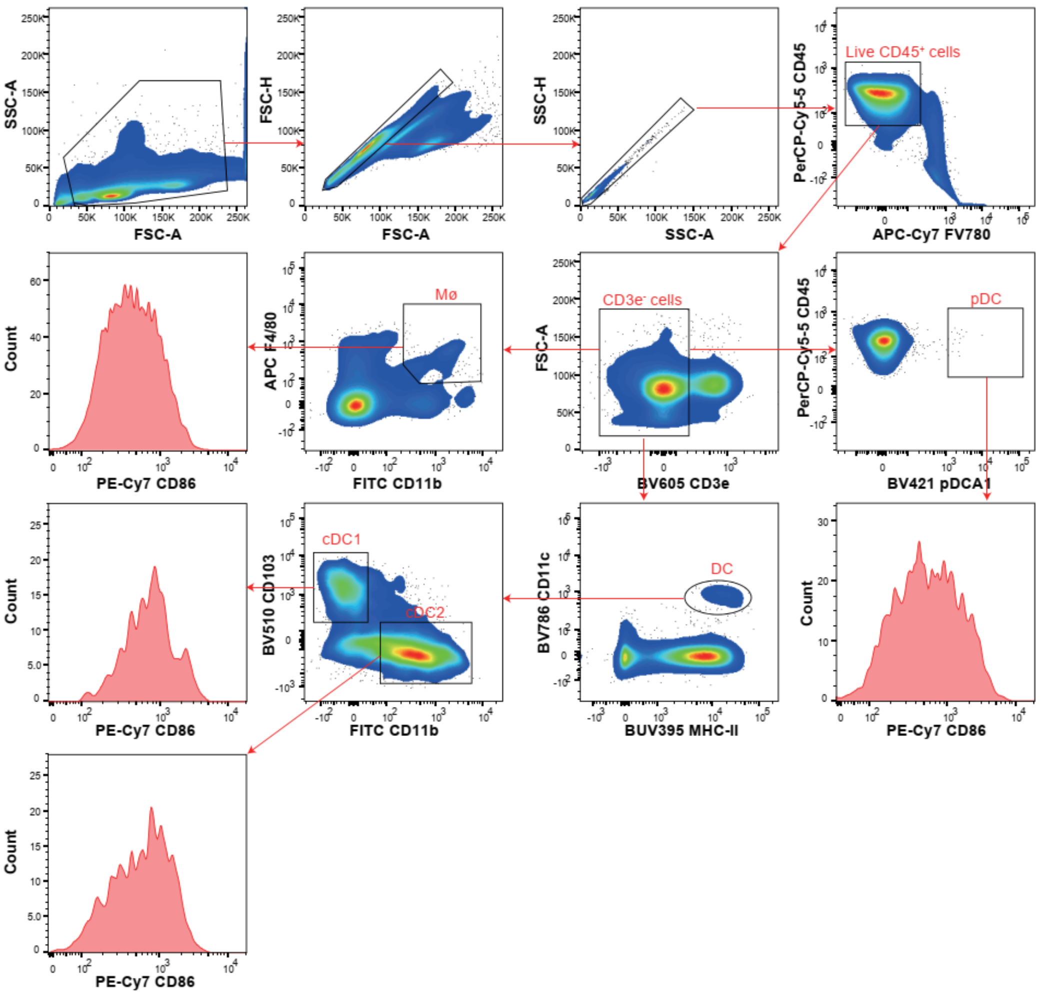

**Supplementary Fig. 27** Gating strategy for flow cytometry analysis of APC response.

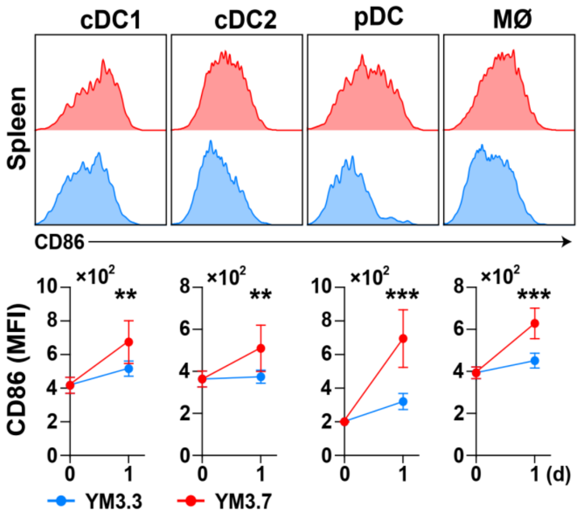

**Supplementary Fig. 28** Flow cytometry analysis of APC maturation in the spleen. n=5 biologically independent replicates. Data are presented as mean ± SD. **Statistical significance test:** Two-way ANOVA followed by Dunn's multiple comparison test (YM3.7 v.s. YM3.3). **: $P < 0.01$, ***: $P < 0.001$.

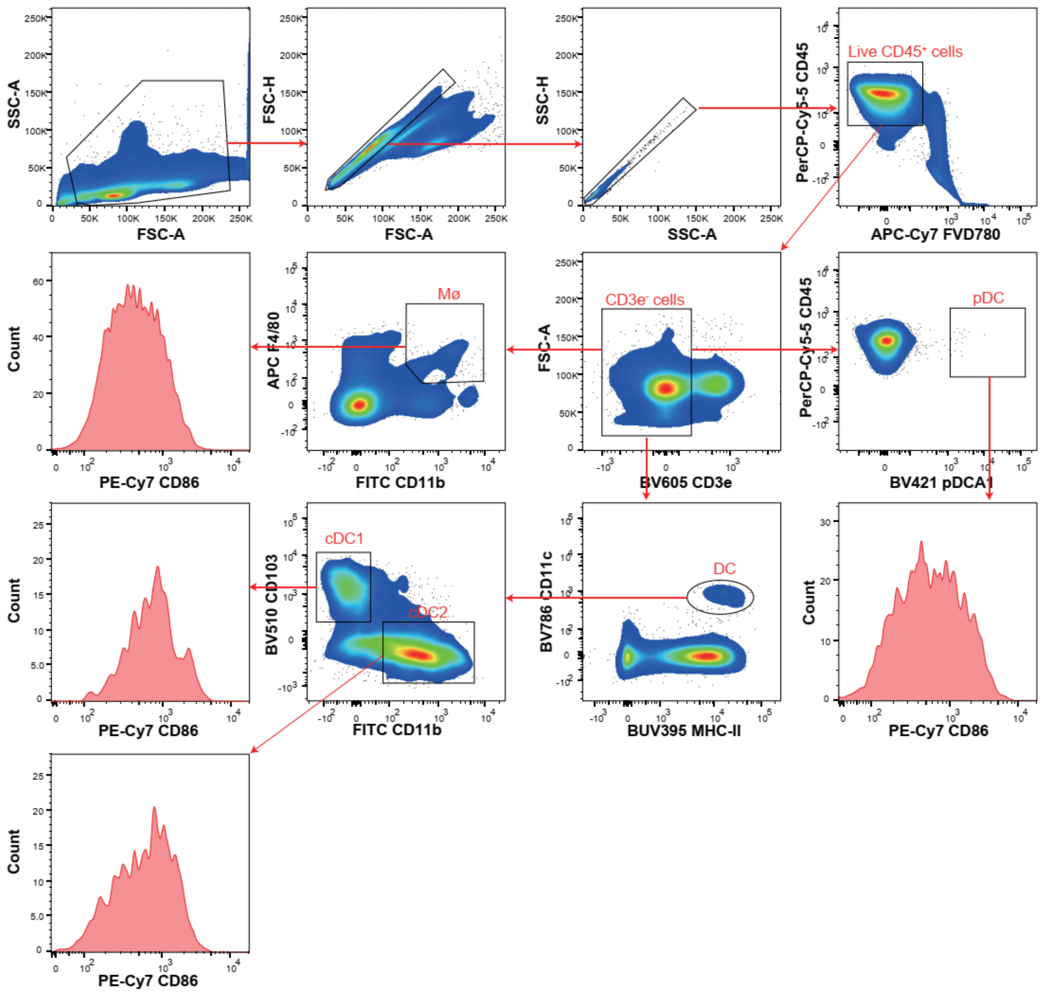

**Supplementary Fig. 29** Gating strategy for flow cytometry analysis of GCB and TFH.

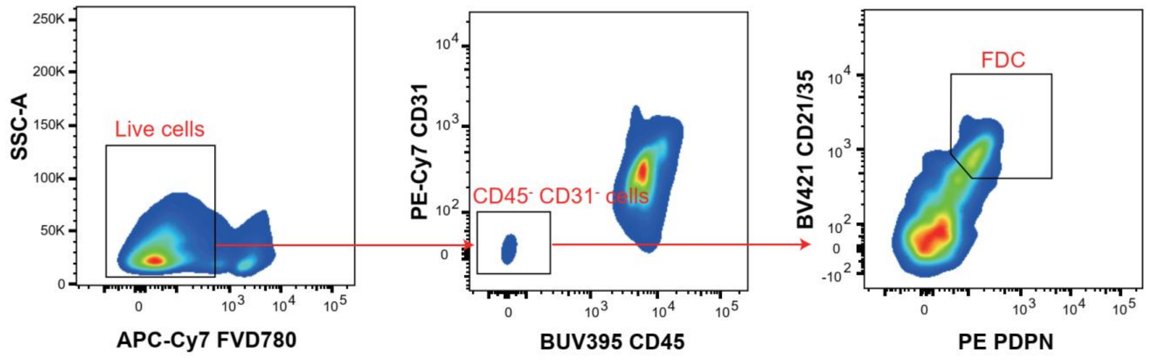

**Supplementary Fig. 30** Gating strategy for flow cytometry analysis of FDC.

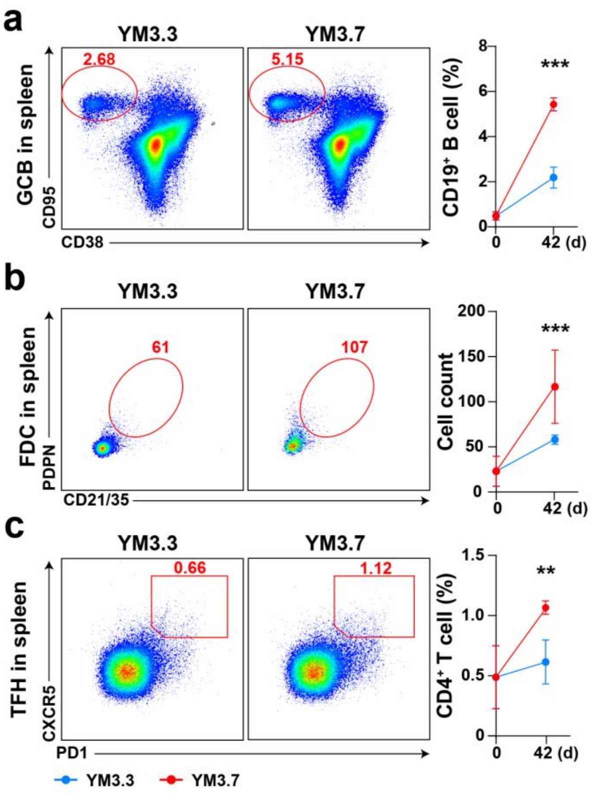

**Supplementary Fig. 31** Flow cytometry analysis of GC response in the spleen. Shown are **a)** GCB, **b)** FDC, and **c)** TFH. n=5 biologically independent replicates. Data are presented as mean ± SD. **Statistical significance test:** Two-way ANOVA followed by Dunn's multiple comparison test (YM3.7 v.s. YM3.3). **: $P < 0.01$, ***: $P < 0.001$.

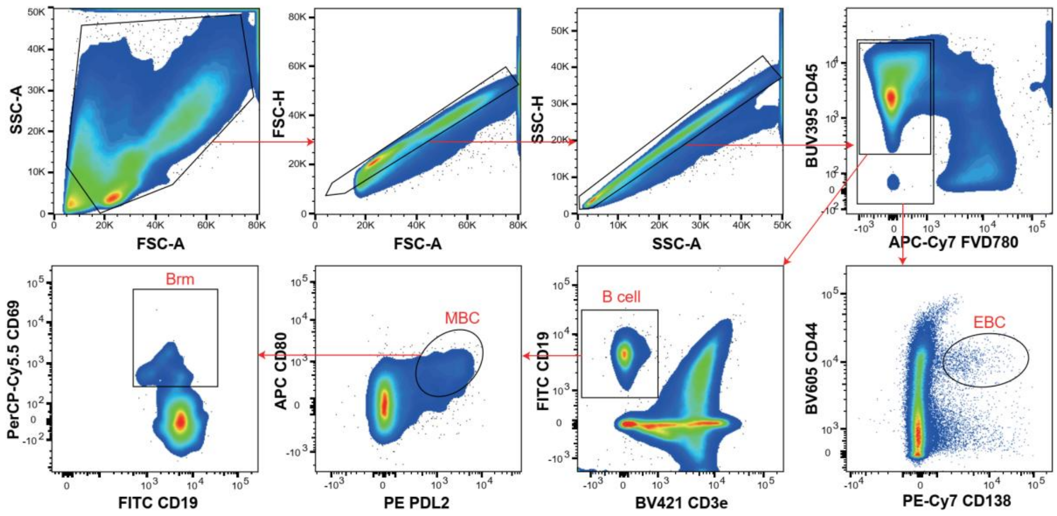

**Supplementary Fig. 32** Gating strategy for flow cytometry analysis of B cell response.

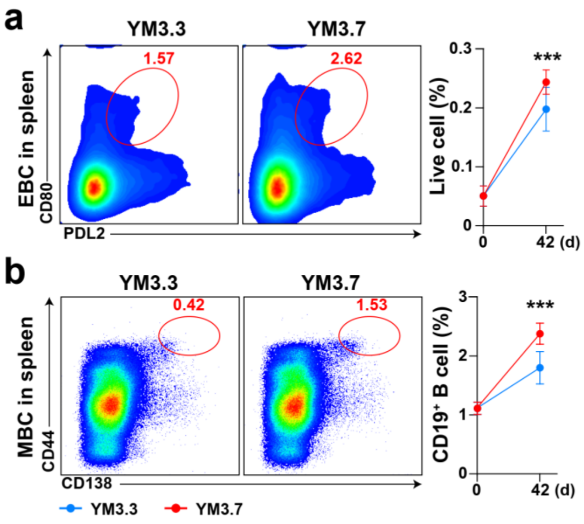

**Supplementary Fig. 33** Flow cytometry analysis of B cell response in the spleen. Shown are **a)** EBC and **b)** MBC. n=5 biologically independent replicates. Data are presented as mean ± SD. **Statistical significance test:** Two-way ANOVA followed by Dunn's multiple comparison test (YM3.7 v.s. YM3.3). ***: $P < 0.001$.

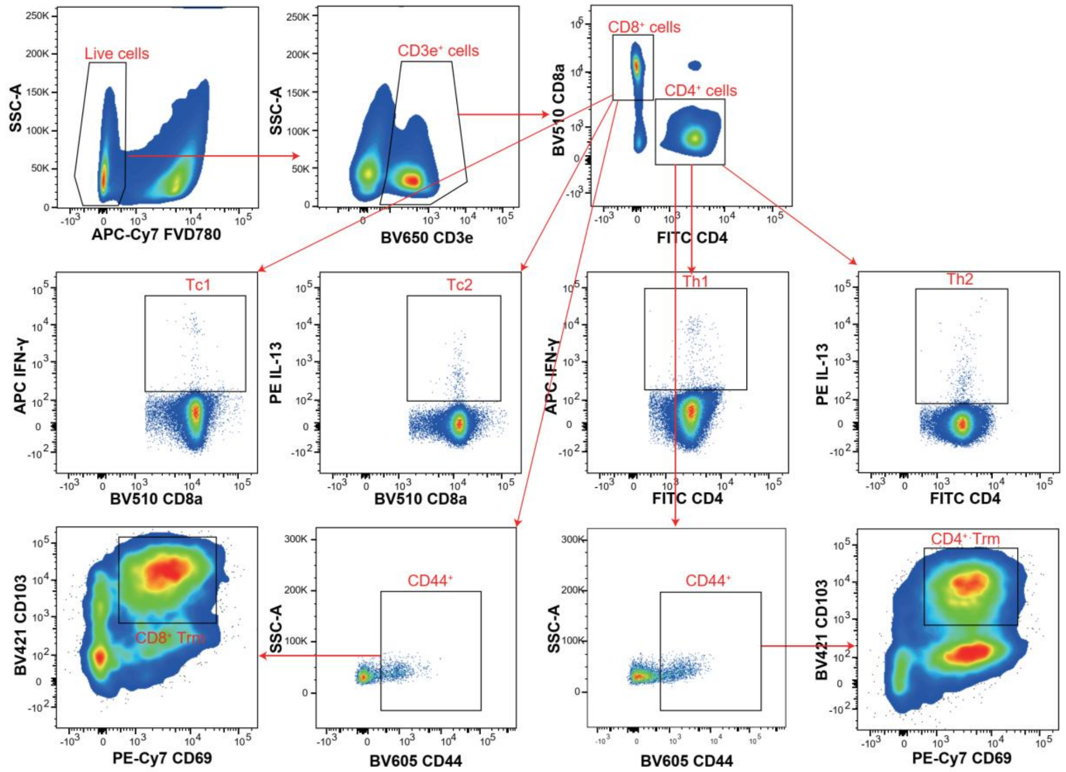

**Supplementary Fig. 34** Gating strategy for flow cytometry analysis of T cell response.

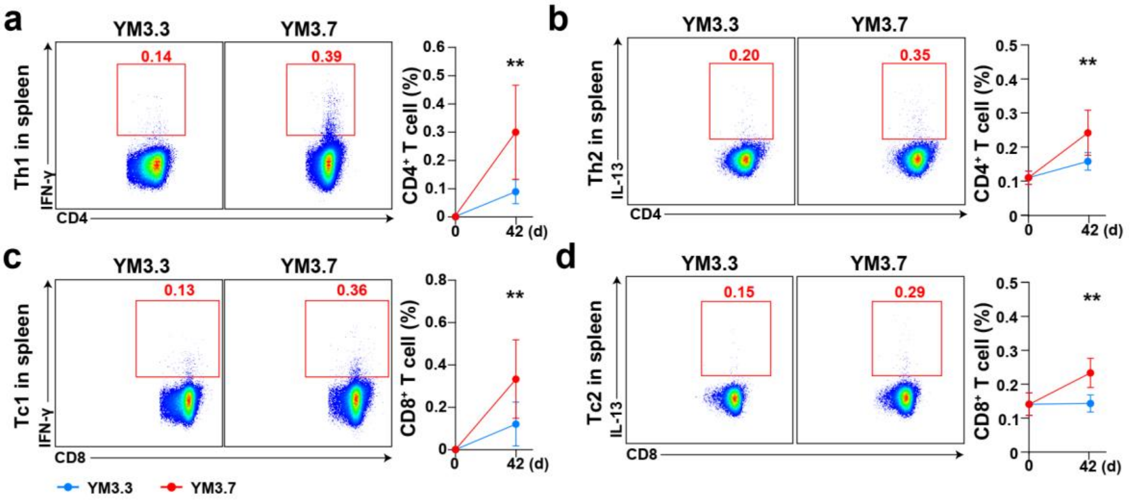

**Supplementary Fig. 35** Flow cytometry analysis of T cell response in the spleen. Shown are **a)** Th1, **b)** Th2, **c)** Tc1, and **d)** Tc2. n=5 biologically independent replicates. Data are presented as mean ± SD. **Statistical significance test:** Two-way ANOVA followed by Dunn's multiple comparison test (YM3.7 v.s. YM3.3). **: $P < 0.01$.

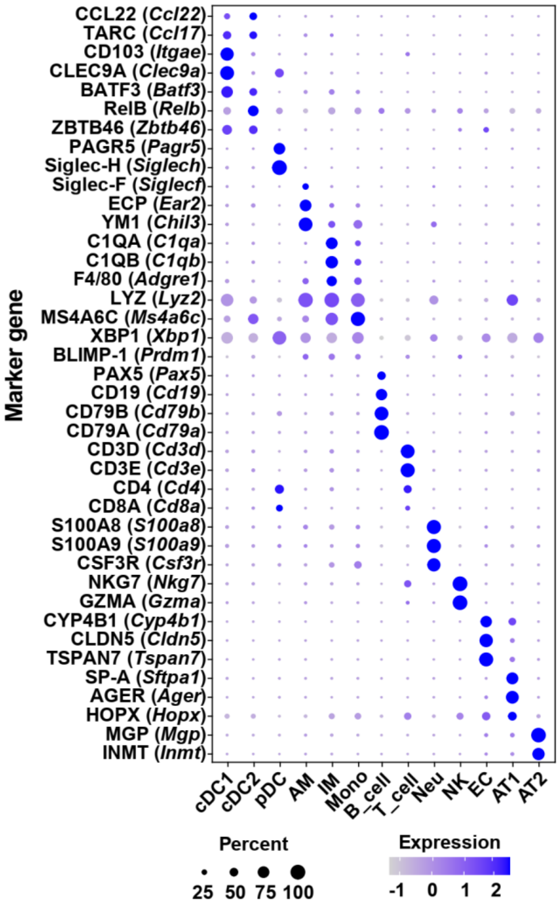

**Supplementary Fig. 36** Cell markers used for clustering of whole cell populations.

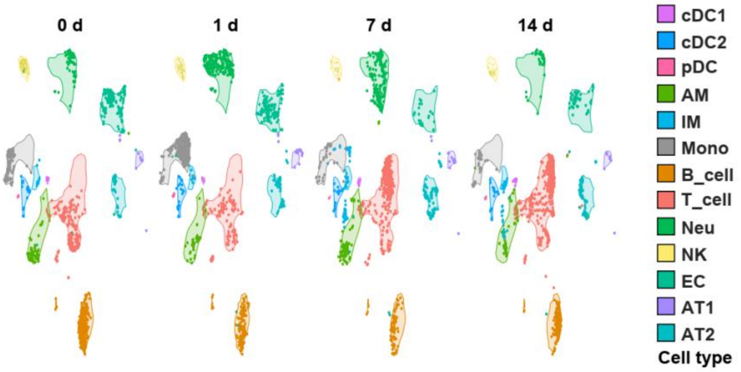

**Supplementary Fig. 37** UMAP plot showing temporal dynamics of cell subsets on 0 d pre-immunization and 1/7/14 d post single-dose immunization.

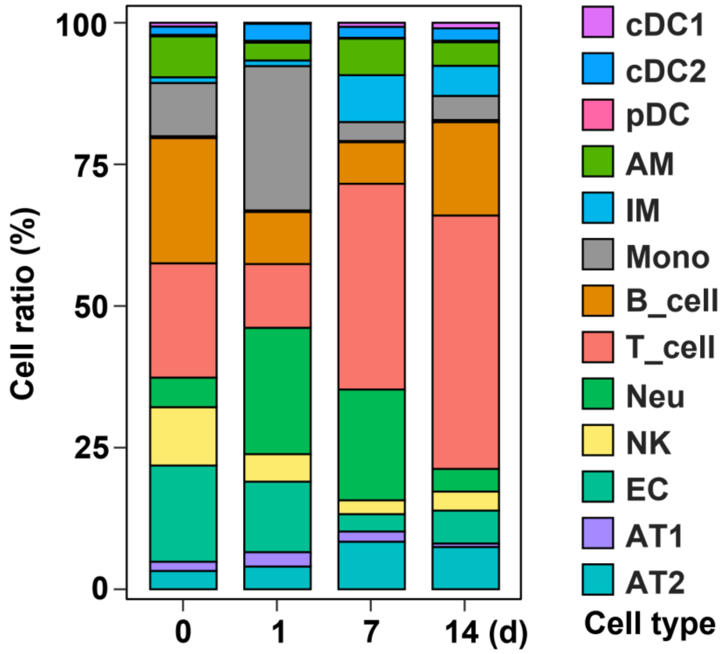

**Supplementary Fig. 38** Stacked bar chart showing changes in proportion of all cell subsets over time.

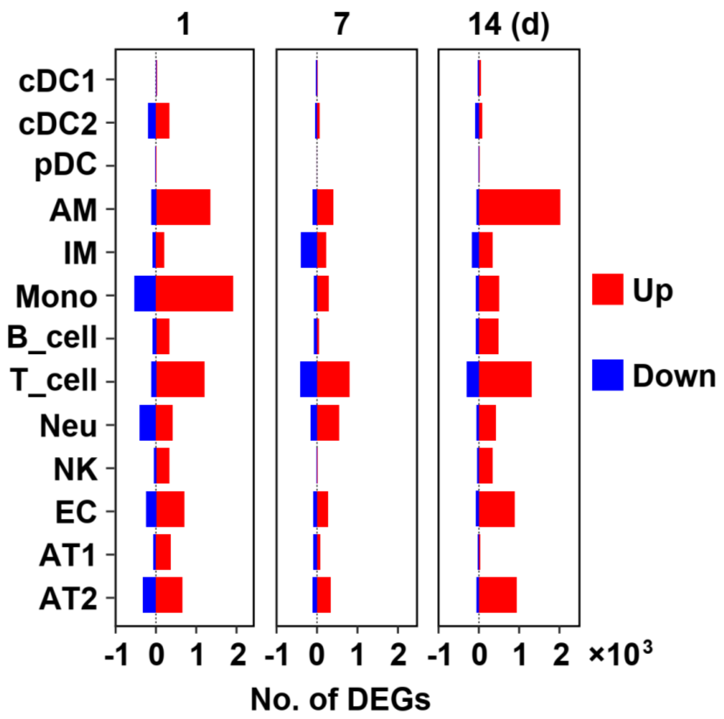

**Supplementary Fig. 39** Histogram showing counts of DEGs in all cell subsets over time.

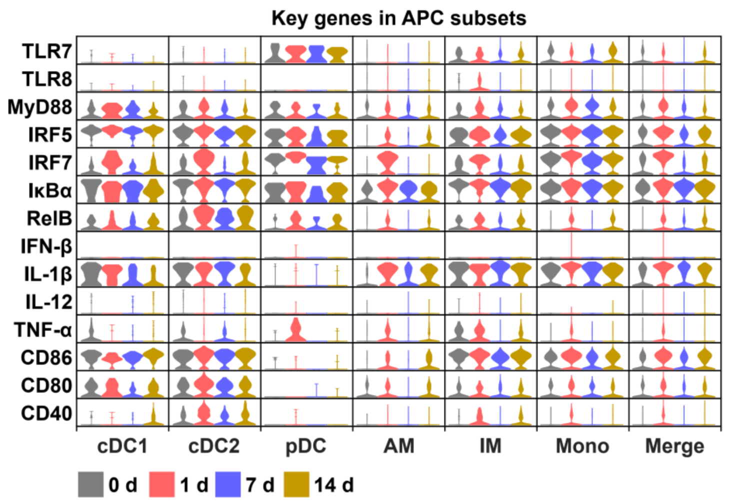

**Supplementary Fig. 40** Violin plot showing expression levels of key genes related to TLR7 signaling in APC subsets.

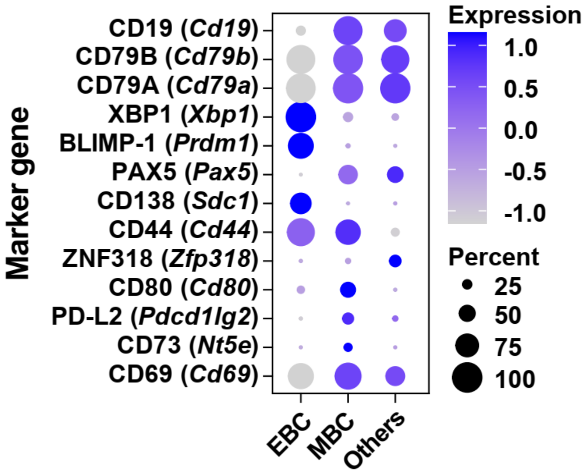

**Supplementary Fig. 41** Cell markers used for clustering of B cell subsets.

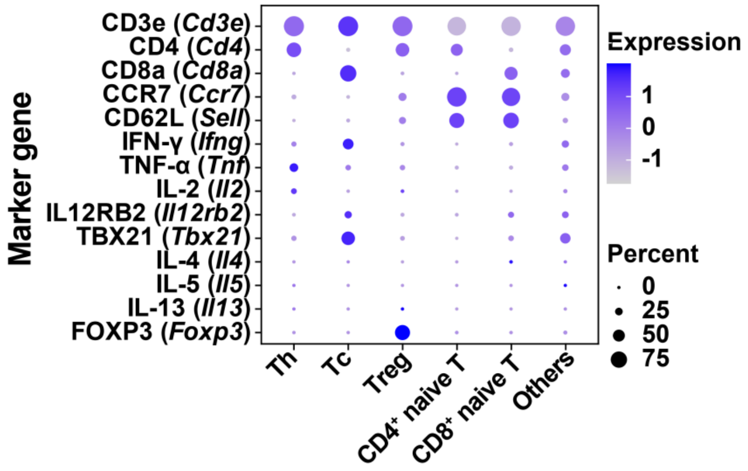

**Supplementary Fig. 42** Cell markers used for clustering of T cell subsets.

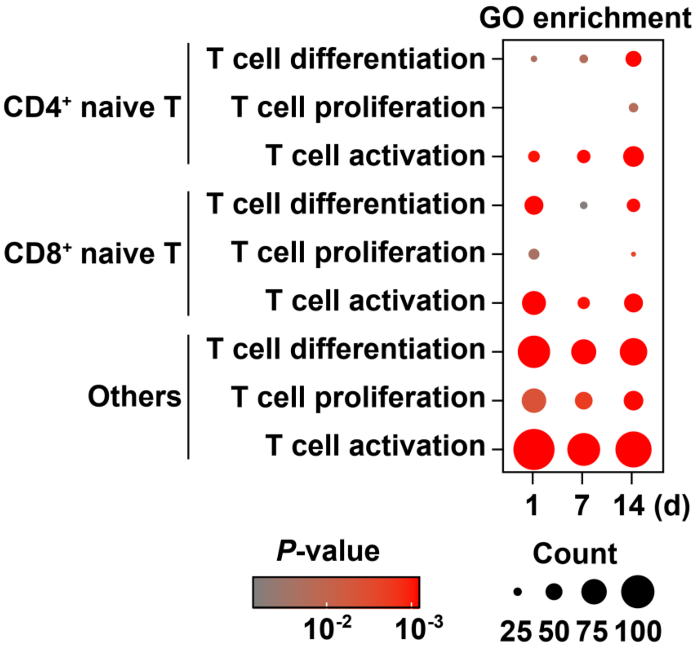

**Supplementary Fig. 43** GO enriched pathways for CD4$^+$ naïve T, CD8$^+$ naïve T, and other T cells.

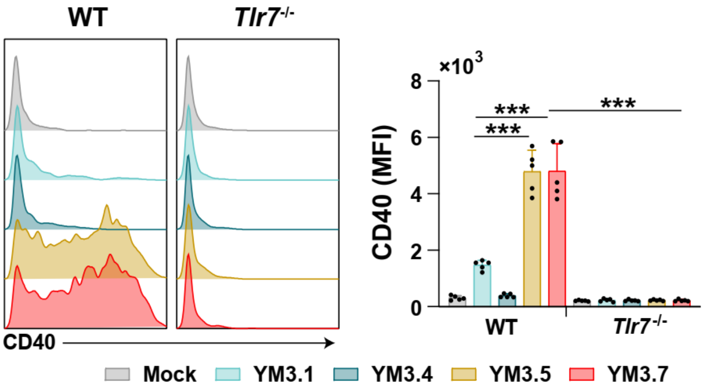

**Supplementary Fig. 44** Expression of CD40 in BMDC after 6-h incubation. BMDC isolated from WT and *Tlr7*$^{-/-}$ mice are incubated with Mock (PBS), YM3.1 (3M-052), YM3.4 (nanoparticle containing YAXA), YM3.5 (nanoparticle containing 3M-052 and YAXA), and YM3.7 (nanoparticle containing 3M-052, YAXA, and VacPAE1). 5.56 µg/mL of VacPAE1, 138.89 µg/mL of YAXA, or 5 µg/mL of 3M-052 are used for treatment. n=5 biologically independent replicates. Data are presented as mean ± SD. **Statistical significance test:** YM3.7 group in WT mice is compared with each of the other groups using one-way ANOVA with Dunn's multiple comparison test. ***: $P$<0.001.

**References**

1. Wan C*, et al.* Rational Design of a Chimeric Derivative of PcrV as a Subunit Vaccine Against Pseudomonas aeruginosa. *Front Immunol* **10**, 781 (2019).
2. Weimer ET, Lu H, Kock ND, Wozniak DJ, Mizel SB. A fusion protein vaccine containing OprF epitope 8, OprI, and type A and B flagellins promotes enhanced clearance of nonmucoid Pseudomonas aeruginosa. *Infect Immun* **77**, 2356-2366 (2009).
3. Yang F*, et al.* Protective Efficacy of the Trivalent Pseudomonas aeruginosa Vaccine Candidate PcrV-OprI-Hcp1 in Murine Pneumonia and Burn Models. *Sci Rep* **7**, 3957 (2017).
4. Kim HK, Cheng AG, Kim HY, Missiakas DM, Schneewind O. Nontoxigenic protein A vaccine for methicillin-resistant Staphylococcus aureus infections in mice. *J Exp Med* **207**, 1863-1870 (2010).
5. Tran VG*, et al.* Efficacy of Active Immunization With Attenuated alpha-Hemolysin and Panton-Valentine Leukocidin in a Rabbit Model of Staphylococcus aureus Necrotizing Pneumonia. *J Infect Dis* **221**, 267-275 (2020).
6. Shilling PJ, Mirzadeh K, Cumming AJ, Widesheim M, Kock Z, Daley DO. Improved designs for pET expression plasmids increase protein production yield in Escherichia coli. *Commun Biol* **3**, 214 (2020).
7. Brito LA, Singh M. Acceptable levels of endotoxin in vaccine formulations during preclinical research. *J Pharm Sci* **100**, 34-37 (2011).
8. Pan X*, et al.* Inhalable MOF-derived nanoparticles for sonodynamic therapy of bacterial pneumonia. *Adv Funct Mater* **32**, 2112145 (2022).
9. Del Barrio-Tofino E, Lopez-Causape C, Oliver A. Pseudomonas aeruginosa epidemic high-risk clones and their association with horizontally-acquired beta-lactamases: 2020 update. *Int J Antimicrob Agents* **56**, 106196 (2020).
10. Stover CK*, et al.* Complete genome sequence of Pseudomonas aeruginosa PAO1, an opportunistic pathogen. *Nature* **406**, 959-964 (2000).
11. Simonis A*, et al.* Discovery of highly neutralizing human antibodies targeting Pseudomonas aeruginosa. *Cell* **186**, 5098-5113 e5019 (2023).
12. Boero E*, et al.* A flow cytometry-based assay to determine the ability of anti-Streptococcus pyogenes antibodies to mediate monocytic phagocytosis in human sera. *J Immunol Methods* **528**, 113652 (2024).

13. Ledger EVK, Edwards AM. Host-induced cell wall remodeling impairs opsonophagocytosis of Staphylococcus aureus by neutrophils. *mBio* **15**, e0164324 (2024).
14. Diep BA*, et al.* Complete genome sequence of USA300, an epidemic clone of community-acquired meticillin-resistant Staphylococcus aureus. *Lancet* **367**, 731-739 (2006).
15. Yin Q*, et al.* A TLR7-nanoparticle adjuvant promotes a broad immune response against heterologous strains of influenza and SARS-CoV-2. *Nat Mater* **22**, 380-390 (2023).
16. Ye T*, et al.* Inhaled SARS-CoV-2 vaccine for single-dose dry powder aerosol immunization. *Nature* **624**, 630-638 (2023).
17. Zhang L*, et al.* A nanovaccine for immune activation and prophylactic protection of atherosclerosis in mouse models. *Nat Commun* **16**, 2111 (2025).
18. Li C*, et al.* Mechanisms of innate and adaptive immunity to the Pfizer-BioNTech BNT162b2 vaccine. *Nat Immunol* **23**, 543-555 (2022).
19. Martinez-Riano A*, et al.* Long-term retention of antigens in germinal centers is controlled by the spatial organization of the follicular dendritic cell network. *Nat Immunol* **24**, 1281-1294 (2023).
20. Zuccarino-Catania GV*, et al.* CD80 and PD-L2 define functionally distinct memory B cell subsets that are independent of antibody isotype. *Nat Immunol* **15**, 631-637 (2014).
21. Allie SR*, et al.* The establishment of resident memory B cells in the lung requires local antigen encounter. *Nat Immunol* **20**, 97-108 (2019).
22. Robinson MD, McCarthy DJ, Smyth GK. edgeR: a Bioconductor package for differential expression analysis of digital gene expression data. *Bioinformatics* **26**, 139-140 (2010).
23. Kumar L, M EF. Mfuzz: a software package for soft clustering of microarray data. *Bioinformation* **2**, 5-7 (2007).
24. Yu G, Wang LG, Han Y, He QY. clusterProfiler: an R package for comparing biological themes among gene clusters. *OMICS* **16**, 284-287 (2012).
25. Su D*, et al.* Spatiotemporal single-cell transcriptomic profiling reveals inflammatory cell states in a mouse model of diffuse alveolar damage. *Exploration (Beijing)* **3**, 20220171 (2023).

26. Yi M, *et al.* Combination of oral STING agonist MSA-2 and anti-TGF-beta/PD-L1 bispecific antibody YM101: a novel immune cocktail therapy for non-inflamed tumors. *J Hematol Oncol* **15**, 142 (2022).
